# Endogenous retroviruses mediate transcriptional rewiring in response to oncogenic signaling in colorectal cancer

**DOI:** 10.1101/2021.10.28.466196

**Authors:** Atma Ivancevic, David M. Simpson, Olivia M. Joyner, Stacey M. Bagby, Lily L. Nguyen, Ben G. Bitler, Todd M. Pitts, Edward B. Chuong

## Abstract

Cancer cells exhibit rewired transcriptional regulatory networks that promote tumor growth and survival. However, the mechanisms underlying the formation of these pathological networks remain poorly understood. Through a pan-cancer epigenomic analysis, we found that primate-specific endogenous retroviruses (ERVs) are a rich source of enhancers displaying cancer-specific activity. In colorectal cancer and other epithelial tumors, oncogenic AP1/MAPK signaling drives the activation of enhancers derived from the primate-specific ERV family LTR10. Functional studies in colorectal cancer cells revealed that LTR10 elements regulate tumor-specific expression of multiple genes associated with tumorigenesis, such as *ATG12* and *XRCC4*. Within the human population, individual LTR10 elements exhibit germline and somatic structural variation resulting from a highly mutable internal tandem repeat region, which affects AP1 binding activity. Our findings reveal that ERV-derived enhancers contribute to transcriptional dysregulation in response to oncogenic signaling and shape the evolution of cancer-specific regulatory networks.

## INTRODUCTION

Cancer cells undergo global transcriptional changes resulting from genetic and epigenetic alterations during tumorigenesis ^1^. While regulatory remodeling can arise from somatic non-coding mutations ^2^, epigenomic studies have revealed that transformation is associated with aberrant epigenetic activation of enhancer sequences that are typically silenced in normal tissues ^3–5^. Pathological enhancer activity is an established mechanism underlying tumorigenesis and therapy resistance, and therapeutic modulation of enhancer activity is an active area of investigation ^6–9^. However, we have a limited understanding of the molecular processes that shape and establish the enhancer landscapes of cancer cells.

Transposable elements (TEs) including endogenous retroviruses (ERVs) represent a potentially significant source of enhancers that could shape cancer-specific gene regulation ^10^. Many cancers exhibit genome-wide transcriptional reactivation of TEs, which can directly impact cells by promoting oncogenic mutations and stimulating immune signaling ^11–14^. In addition, the reactivation of TEs is increasingly recognized to have gene regulatory consequences in cancer cells ^15,16^. Several transcriptomic studies have uncovered TEs as a source of cancer-specific alternative promoters across many types of cancer, with some examples shown to drive oncogene expression ^17–21^. TEs also show chromatin signatures of enhancer activity in cancer cell lines ^22–24^, yet their functional relevance in patient tumors has remained largely unexplored. Recent studies have characterized TE-derived enhancers with oncogenic effects in acute myeloid leukemia ^25^ and prostate cancer ^26^ but the prevalence and mechanisms of TE-derived enhancer activity are unknown for most cancer types.

Here, we analyzed published cancer epigenome datasets to understand how TEs influence enhancer landscapes and gene regulation across cancer types. Our pan-cancer analysis revealed that elements from a primate-specific ERV named LTR10 show enhancer activity in many epithelial tumors, and this activity is regulated by MAPK/AP1 signaling. We conducted functional studies in HCT116 colorectal cancer cells, and found that LTR10 elements regulate AP1-dependent gene expression at multiple loci that include genes with established roles in tumorigenesis. Finally, we discovered that LTR10 elements contain highly mutable sequences that potentially contribute genomic variation affecting cancer-specific gene expression. Our work implicates ERVs as a source of pathological regulatory variants that facilitate transcriptional rewiring in cancer.

## RESULTS

To assess the contribution of TEs to cancer cell epigenomes, we analyzed aggregate chromatin accessibility maps from 21 human cancers generated by The Cancer Genome Atlas project ^27^. We defined cancer-specific subsets of accessible regions by subtracting regions that show evidence of regulatory activity in any healthy adult tissue profiled by the Roadmap Consortium (Fig 1A, Methods) ^28^. Out of 1315 total repeat subfamilies annotated in the human genome, we found 23 subfamilies that showed significant enrichment within the accessible chromatin in at least one cancer type (Fig 1B), of which 19 correspond to long terminal repeat (LTR) regions of primate-specific ERVs (Supp Table 1). These observations from chromatin accessibility data generated from primary tumors confirm previous reports of LTR-derived regulatory activity in cancer cell lines ^22,24,25^, and support a role for ERVs in shaping patient tumor epigenomes.

**Figure 1:**
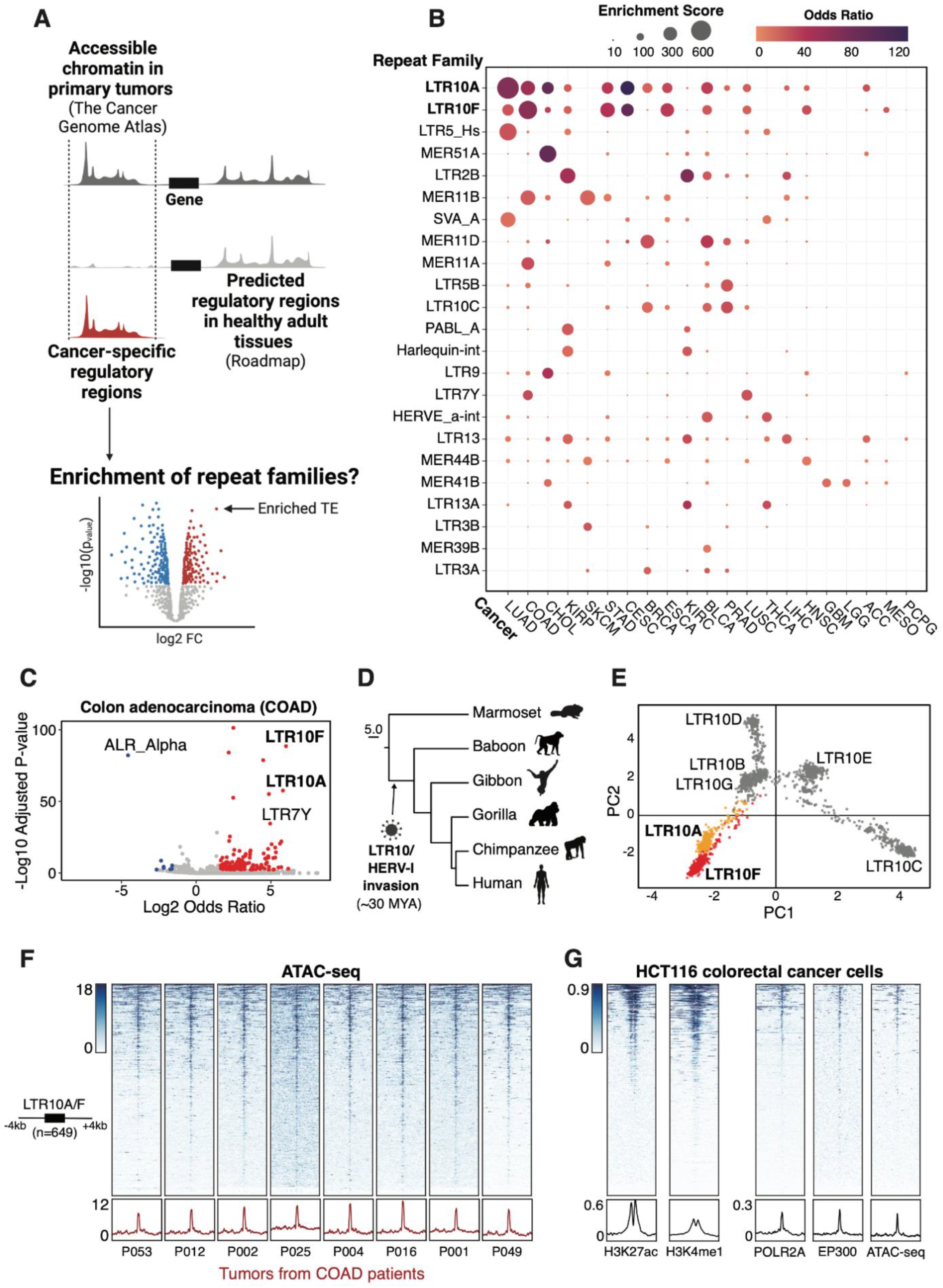
Pan-cancer epigenomic analysis of TE activity. **(A)** Pipeline to estimate TE subfamily enrichment within cancer-specific regulatory regions. Aggregate ATAC-seq maps associated with each TCGA tumor type were filtered to remove regulatory regions predicted in any normal adult tissues by Roadmap. Cancer-specific accessible chromatin regions were tested for enrichment of 1315 repeat subfamilies using GIGGLE. **(B)** Bubble chart summarizing TE subfamily enrichment within cancer-specific ATAC-seq regions across 21 cancer types profiled by TCGA (acronyms shown on the x-axis; full names provided at https://gdc.cancer.gov/resources-tcga-users/tcga-code-tables/tcga-study-abbreviations). TE subfamilies and cancer types are sorted based on maximum enrichment score. **(C)** Enrichment of TE subfamilies within cancer-specific ATAC-seq associated with colon adenocarcinomas (COAD) from TCGA. Every point represents a TE subfamily. Significantly enriched TEs are shown in red; depleted TEs are shown in blue. **(D)** Estimated origin of HERV-I elements on the primate phylogeny based on genomic presence or absence. **(E)** Principal component analysis based on multiple sequence alignment of all LTR10 sequences over 200 bp in length in the human genome (n=1806). Every point represents an individual LTR10 sequence. LTR10A and LTR10F sequences are colored orange and red, respectively. **(F)** Heatmap of representative patient tumor ATAC-seq signals (TCGA patients COAD P053, P012, P002, P025, P004, P016, P001, P049) over the merged set of 649 LTR10A/F elements. Bottom metaprofiles represent average normalized ATAC signal across elements. **(G)** Heatmap of enhancer-associated chromatin marks from HCT116 cells over the merged set of 649 LTR10A/F elements. From left to right: H3K27ac ChIP-seq (GSE97527), H3K4me1 ChIP-seq (GSE101646), POLR2A ChIP-seq (GSE32465), EP300 ChIP-seq (GSE51176), and HCT116 ATAC-seq (GSE126215). Bottom metaprofiles represent the normalized signal across elements.

### LTR10 elements exhibit cancer-specific regulatory activity

To further investigate the cancer-specific regulatory activity of ERVs, we focused on LTR10 elements, which were enriched within cancer-specific accessible chromatin for several types of epithelial tumors including colorectal, stomach, prostate, and lung tumors (Fig 1C, Supp Fig S1A). LTR10 elements (including LTR10A-G, n=2331) are derived from the LTR of the gammaretrovirus HERV-I, which integrated into the anthropoid genome 30 million years ago (Fig 1D, 1E) ^22^. As our initial TCGA analysis was conducted using aggregate data for each tumor type, we first confirmed that LTR10 elements showed recurrent chromatin accessibility across colorectal tumors from multiple individual patients (Fig 1F, Supp Fig S1B). We then analyzed epigenomic datasets from the HCT116 colorectal cancer cell line ^3,29–31^ and found that LTR10A and LTR10F elements exhibit canonical chromatin hallmarks of enhancer activity, including enrichment of histone modifications H3K27ac and H3K4me1, the transcriptional coactivator p300, and RNA Polymerase II occupancy (Fig 1G).

We did not observe enhancer-like profiles at LTR10C elements, which have previously been identified as a source of p53 binding sites ^22,32^ (Supp Fig S1C). While most LTR10A and LTR10F elements are not transcribed, some show evidence of transcription as promoters for full-length non-coding HERV-I insertions or cellular transcripts (Supp Fig S1D). Therefore, elements derived from the LTR10A and LTR10F subfamilies (hereafter referred to as LTR10 elements) show robust epigenomic signatures associated with enhancer activity in colorectal cancer cells.

We expanded our analysis to include epigenomic states from all adult tissues ^28^. We found no evidence for LTR10 enhancer activity in normal tissues, but instead observed general enrichment of H3K9me3-associated heterochromatin marks (Supp Fig S2A, Supp Table 2). To identify factors that directly bind to and potentially repress LTR10 elements, we analyzed the Cistrome database ^31^ of published human ChIP-Seq datasets to identify transcriptional repressors with evidence for enriched binding within LTR10 elements. Considering all cell types, we found that LTR10 elements are significantly enriched for binding by ZNF562, TRIM28, and SETDB1 (Fig 2A, 2B, Supp Table 3, Supp Table 4), which are components of the KRAB-ZNF transposon silencing pathway ^33^. In additional datasets generated from healthy colorectal tissue samples ^3,34–36^, LTR10 elements do not show any evidence of enhancer activity (Supp Fig S2B). Our analysis suggests that, as expected for most primate-specific TEs ^37^, LTR10 elements are normally subject to H3K9me3-mediated epigenetic silencing in somatic tissues.

**Figure 2:**
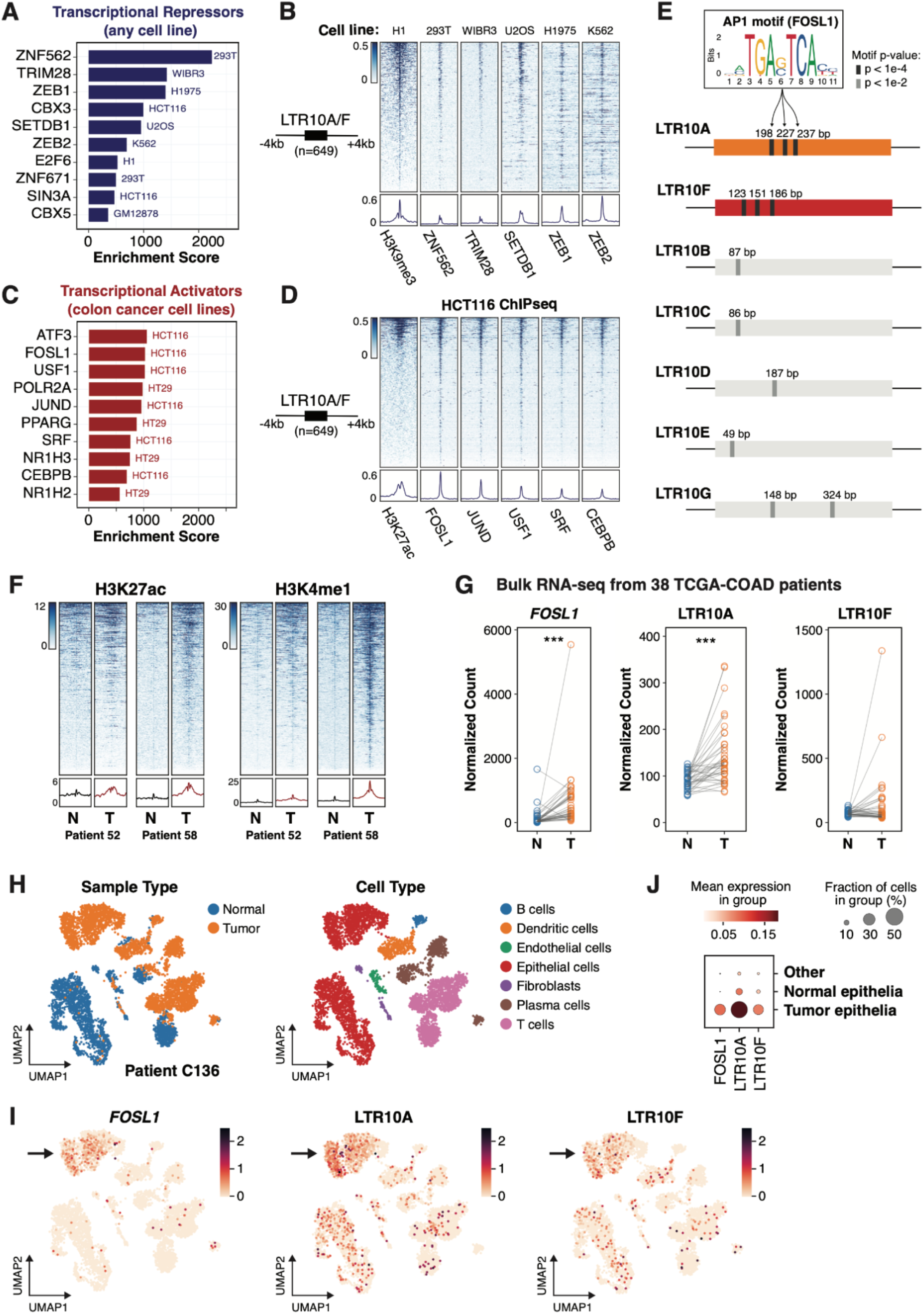
Regulatory activity of LTR10 in tumor and normal cells. **(A)** Transcriptional repressors associated with LTR10A/F elements, ranked by enrichment score. **(B)** Heatmap of ChIP-seq signal from H3K9me3 and repressive factors, over LTR10A/F elements. From left to right: H3K9me3 ChIP-seq (GSE16256), ZNF562 ChIP-seq (GSE78099), TRIM28 ChIP-seq (GSE84382), SETDB1 ChIP-seq (GSE31477), ZEB1 (GSE106896), and ZEB2 ChIP-seq (GSE91406). **(C)** Transcriptional activators associated with LTR10A/F elements, ranked by enrichment score. **(D)** Heatmap of ChIP-seq signal from H3K27ac and activating transcription factors in HCT116 cells, over LTR10A/F elements. From left to right: H3K27ac ChIP-seq (GSE96299), as well as ChIP-seq for FOSL1, JUND, USF1, SRF and CEBPB (all from GSE32465). **(E)** Schematic of AP1 motif locations for LTR10 consensus sequences from each subfamily. Sequence logo for AP1 motif FOSL1 (MA0477.1 from JASPAR) is shown, and predicted motif locations are marked on each consensus. **(F)** Heatmap of H3K27ac and H3K4me1 ChIP-seq signals from tumor (T) and normal (N) samples from colorectal cancer patients (CEMT patients AKCC52 and AKCC58) over LTR10A/F elements. Bottom metaprofiles represent average normalized ChIP signal. **(G)** Dot plots of normalized counts for *FOSL1*, LTR10A and LTR10F from bulk RNA-seq derived from a cohort of 38 TCGA patients with colorectal adenocarcinomas. Each patient has one tumor (T) sample and one normal (N) colon sample. ***: p < 0.001, paired sample Wilcoxon test. **(H)** UMAP projections of the single cell transcriptome of patient C136 from Pelka et al (2021). UMAPs are colored according to tissue type or cell type. **(I)** UMAP projections of the same patient, colored according to the expression of *FOSL1*, LTR10A, or LTR10F. In each case, the arrow points to the tumor-specific cluster of epithelial cells. **(J)** Bubble plot of the same patient, showing the mean expression of *FOSL1*, LTR10A and LTR10F in tumor epithelia vs normal epithelia.

### LTR10 elements are bound by the AP1 transcription factor complex

To identify which pathways are responsible for cancer-specific reactivation of LTR10 elements, we focused our Cistrome enrichment analysis on activating transcription factors in colorectal cancer cell lines. LTR10 elements were significantly enriched for binding by AP1 complex members (Fig 2C, 2D, Supp Table 3) including the FOSL1, JUND, and ATF3 transcription factors. The LTR10A and LTR10F consensus sequences harbor multiple predicted AP1 binding motifs, which are enriched within LTR10 elements marked by H3K27ac in HCT116 cells. Moreover, the AP1 motifs are largely absent in other LTR10 subfamilies (Fig 2E). Expanding our motif analysis to tumor-specific accessible chromatin from 21 different cancer types, we found that AP1 motif enrichment generally correlates with LTR10 enrichment, especially for LTR10A (Supp Fig S2C). In contrast, cancers without LTR10 enrichment show little to no enrichment of AP1 motifs in tumor-specific accessible chromatin (Supp Fig S2C). These analyses indicate that the cancer-specific enhancer activity of LTR10 elements is likely driven by sequence-specific recruitment of the AP1 complex.

### LTR10 epigenetic and transcriptional activity is elevated in patient tumor cells

We next compared the epigenetic status of LTR10 elements between patient-derived colorectal cancer cells and normal cells. In multiple patient-matched epigenomic datasets ^38,39^, LTR10 elements show globally increased levels of enhancer-associated histone modifications H3K27ac and H3K4me1 in tumor samples compared to adjacent normal colorectal tissues (Fig 2F, Supp Fig S2D). In contrast, LTR10 elements did not show global changes in H3K9me3 or H3K27me3 ChIP-seq signal in tumors compared to normal cells (Supp Fig S2E). These observations suggest that removal of repressive histone marks may not be required for LTR10 enhancer activity, however, single-cell epigenomic profiling would be necessary to determine whether LTR10 elements are marked by both active and repressive marks in the same cells.

We further assessed the transcriptional activity of LTR10 elements using matched tumor/normal RNA-seq from 38 patients with colorectal adenocarcinomas from TCGA controlled access data ^27^ (Supp Fig S2F). Our RNA-seq analysis of the patient cohort suggests that LTR10 transcripts are generally increased in tumor versus normal samples, particularly at LTR10A elements (Fig 2G, Supp Fig S2G, Supp Table 5). Likewise, AP1 factor *FOSL1* showed a robust and significant increase in expression in tumor versus normal samples (Fig 2G, Supp Fig S2H, Supp Table 5), consistent with our hypothesis that the AP1 complex drives LTR10 transcriptional activity. Altogether, 15 of the 38 patients show a consistent increase in *FOSL1*, LTR10A and LTR10F transcriptional activity in colorectal tumor cells (Supp Table 5).

### LTR10 transcription marks tumor-specific epithelial cells

We next investigated LTR10 transcription at the single-cell level. We analyzed an independent cohort of 36 colorectal cancer patients with publicly available single-cell RNA-seq from matched tumor and normal samples for each patient ^40^. We used scTE ^41^ to reprocess the datasets and measure cell population-specific expression of TE subfamilies. In line with our previous results from bulk RNA-seq, we found significant and recurrent transcription of LTR10 elements in tumor-specific epithelial cells for 12 out of 36 patients (Fig 2H-J, Supp Fig S2I & S2J, Supp Table 6). We observed co-expression of LTR10 and *FOSL1* in tumor-specific epithelial cells for 10 of these patients (Supp Table 6), consistent with a role for AP1 signaling in regulating LTR10 elements. Thus, our single-cell analysis indicates that a subset of patients show robust LTR10 transcriptional activity specifically in tumor-specific epithelial cells.

### LTR10 transcription is associated with dysregulated MAPK signaling

Our initial analyses of patient cohorts suggest that LTR10 elements become transcriptionally activated in about 30% of colorectal tumors. To determine which tumor molecular subtypes are most likely to drive LTR10 activation, we performed correlative studies between LTR10 activity and tumor mutations, patient survival rates, and clinical outcomes. For this purpose, we obtained and analyzed RNA-seq from 358 primary tumor samples derived from TCGA patients with colon adenocarcinomas or adenomas ^27^. We first focused on correlating LTR10 transcriptional activity with KRAS mutation status. KRAS is one of the most frequently mutated oncogenes in cancer: approximately 30-40% of colorectal cancer patients harbor missense mutations in KRAS, and KRAS mutations have long been associated with increased tumor aggressiveness, resistance to treatment, and poor patient outcomes ^42^. We found that LTR10A transcripts, in particular, are significantly elevated in tumors that harbor a KRAS mutation (Supp Fig S2K, Supp Table 7), although we did not observe a noticeable difference in *FOSL1* expression (Supp Fig S2L, Supp Table 7). Survival analyses based on the expression of LTR10 transcripts or AP1 factors did not reveal any significant correlations (Supp Fig S2M, S2N). However, we note that our analyses might have been limited by the fact that subfamily-level LTR10 transcription by RNA-seq is an imperfect proxy for enhancer activity, and only a small subset of colorectal cancer samples from TCGA had associated survival data, limiting our statistical power.

### AP1 signaling is required for LTR10 enhancer activity

Dysregulation of AP1 signaling occurs in many cancers, driven by mutations that cause oncogenic activation of the MAPK signaling pathway ^43^. Based on our findings that LTR10 elements are bound by AP1, and LTR10 transcriptional activity is correlated with the expression of AP1 component *FOSL1*, we tested whether LTR10 regulatory activity is affected by modulation of the AP1/MAPK signaling pathway using luciferase reporter assays. We synthesized the LTR10A and LTR10F consensus sequences as well as variants where the AP1 motifs were disrupted, and cloned the sequences into an enhancer reporter construct. We measured reporter activity in HCT116 cells that were treated for 24 hrs with either TNFa to stimulate signaling or cobimetinib (a MEK1 inhibitor) to inhibit signaling. Consistent with regulation by AP1, cobimetinib treatment caused a decrease in LTR10-driven reporter activity, and TNFa caused an increase (Fig 3A). Overall regulatory activity was greatly reduced in sequences where the AP1 motif was disrupted (Fig 3A). These results show that LTR10 enhancer activity can be directly regulated by modulation of the AP1/MAPK signaling pathway in cancer cells.

**Figure 3:**
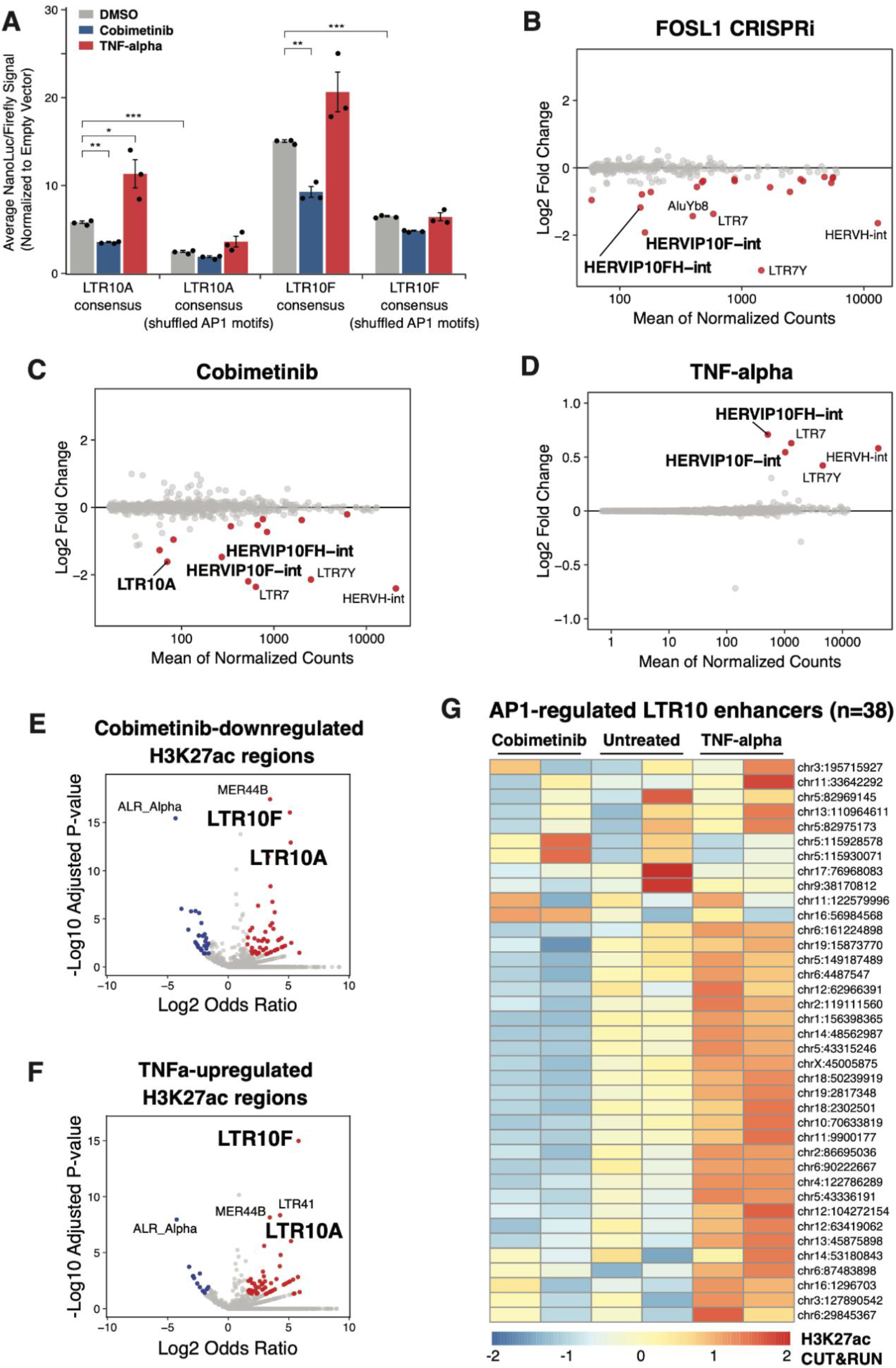
Control of LTR10 activity by AP1/MAPK signaling. **(A)** Luciferase reporter assays of LTR10A/F consensus sequences, including sequence variants containing shuffled AP1 motifs. Reporter activity was measured in HCT116 cells treated with DMSO (n=3), cobimetinib (n=3), or TNF-alpha (n=3) for 24 hrs. Values are normalized to firefly co-transfection controls, and presented as fold-change against the mean values from cells transfected with a empty minimal promoter pNL3.3 vector. *: p < 0.05, **: p < 0.01, ***: p < 0.001, two-tailed student’s t-test. Error bars denote SEM. **(B-D)** MA plots of TE subfamilies showing significant differential expression in HCT116 cells subject to FOSL1 silencing **(B)**, 24hr cobimetinib treatment **(C)**, or 24hr TNF-alpha treatment **(D)**, based on RNA-seq. Dots are colored in red if they are significant (adjusted p<0.05, log2FC<0 for FOSL1/cobimetinib and log2FC>0 for TNFa). **(E)** Volcano plot showing TE subfamily enrichment in the set of H3K27ac regions significantly downregulated by cobimetinib. **(F)** Volcano plot showing TE subfamily enrichment in the set of H3K27ac regions significantly upregulated by TNF-alpha. **(G)** Heatmap of normalized H3K27ac CUT&RUN signal for 38 LTR10 elements predicted to function as enhancers regulating AP1 target genes for each treatment replicate.

To test the role of the AP1 complex in endogenous LTR10 regulation, we silenced the AP1 component *FOSL1* using CRISPRi to determine the impact on LTR10 transcriptional activity. Using HCT116 cells expressing dCas9-KRAB-MeCP2 ^44^, we transfected a guide RNA (gRNA) targeting the *FOSL1* transcription start site (TSS) and used RNA-seq to compare gene and TE expression to control cells transfected with a non-targeting gRNA. We first confirmed silencing of *FOSL1* (Supp Fig S3A, S3B), then analyzed TE transcript expression summarized at the subfamily level to account for reads mapping to multiple insertions of the same TE ^45^. This analysis revealed that full-length LTR10/HERV-I elements were significantly downregulated upon silencing *FOSL1* (Fig 3B), supporting a direct role for the AP1 complex in regulating LTR10 activity.

Next, we investigated how endogenous LTR10 elements respond to modulation of MAPK/AP1 signaling at the chromatin level. We treated HCT116 cells with either cobimetinib or TNFa for 24 hrs and profiled each response using RNA-seq and H3K27ac CUT&RUN (Supp Fig S3C, S3D, S3E, S3F). Consistent with our reporter assay results, our RNA-seq analysis showed that full-length LTR10/HERV-I transcripts were significantly downregulated upon cobimetinib treatment (Fig 3C), and upregulated upon TNFa treatment (Fig 3D). LTR10 elements showed similar responses based on H3K27ac CUT&RUN signal, exhibiting significant enrichment within the genome-wide set of predicted enhancers downregulated by cobimetinib or upregulated by TNFa (Fig 3E, 3F). We also observed clear TNFa-induced H3K27ac signal over LTR10 elements in a published dataset of SW480 colorectal cancer cells ^46^ (Supp Fig S3G). These results indicate that LTR10 elements represent a significant subset of genome-wide enhancers and transcripts in HCT116 cells that are directly modulated by MAPK/AP1 signaling.

### LTR10 elements regulate cancer-specific pathological gene expression

To determine whether any LTR10-derived enhancers have a functional effect on AP1/MAPK-dependent gene expression in colorectal cancer cells, we used our RNA-seq and CUT&RUN data from HCT116 cells to identify elements predicted to have gene regulatory activity. While we found that the AP1 component FOSL1 is required for LTR10 regulatory activity, oncogenic MAPK signaling can mediate transcriptional dysregulation through additional pathways beyond FOSL1 and AP1 signaling ^47^. Therefore, we defined potential AP1/MAPK-regulated genes using two approaches, based on our RNA-seq data from our FOSL1 knockdown or TNFa/cobimetinib treatment. We first defined a set of 456 AP1-dependent genes based on being significantly downregulated by our CRISPRi silencing of the AP1 component *FOSL1* (Supp Table 8). We identified LTR10 elements predicted to regulate these genes using the activity by contact model ^48^ to assign enhancer-gene targets based on LTR10 element H3K27ac signal and chromatin interaction data. This identified 38 LTR10-derived enhancers (Fig 3G) predicted to regulate 56 (12.2%) of the 456 AP1-dependent genes (Supp Table 8), including many with established roles in cancer pathophysiology. In a secondary analysis, we defined 620 MAPK-dependent genes as genes that are both upregulated by TNFa and downregulated by cobimetinib, and found 57 LTR10-derived enhancers predicted to regulate 74 (11.9%) of these genes (Supp Fig S3H, Supp Table 9). Collectively, we identified a total of 71 distinct LTR10 enhancers (Supp Table 10) predicted to contribute to the regulation of roughly 12% of genes with AP1 or MAPK-dependent gene expression in HCT116 cells, supporting an important role in mediating global transcriptional rewiring in cancer.

We tested the regulatory activity of six predicted LTR10 enhancers using CRISPR to knock down or knock out individual elements in HCT116 cells. We prioritized the elements based on epigenomic evidence of tumor-specific enhancer activity and having predicted target genes with reported relevance to tumor development or therapy resistance. We separately silenced each LTR10 element using CRISPRi and selected one element (LTR10.KDM6A) to delete using CRISPR/Cas9, due to its intronic location. We used RNA-seq to determine the transcriptional consequences of perturbing each element. For each LTR10 tested, we observed local downregulation of multiple genes within 1.5 MB of the targeted element, confirming their activity as functional enhancers in HCT116 cells. These included *ATG12, XRCC4, TMEM167A, VCAN, NES, FGF2, AGPAT5, MAOB,* and *MIR222HG* (Fig 4, Fig 5, Supp Fig S4A-S4J, Supp Tables 11-16). For three elements (LTR10.MEF2D, LTR10.MCPH1 and LTR10.KDM6A), the predicted target gene did not show significant expression changes, but we observed downregulation of other AP1/MAPK-dependent genes near the element (Supp Fig S4B, S4F, S4H). Collectively, our characterization of six LTR10 elements verified that 21 genes are regulated by LTR10 elements; most (18/21) of which are regulated by AP1/MAPK signaling based on our RNA-seq data. These experiments demonstrate that multiple LTR10-derived enhancers mediate AP1/MAPK-dependent gene expression of nearby genes in HCT116 cells.

**Figure 4:**
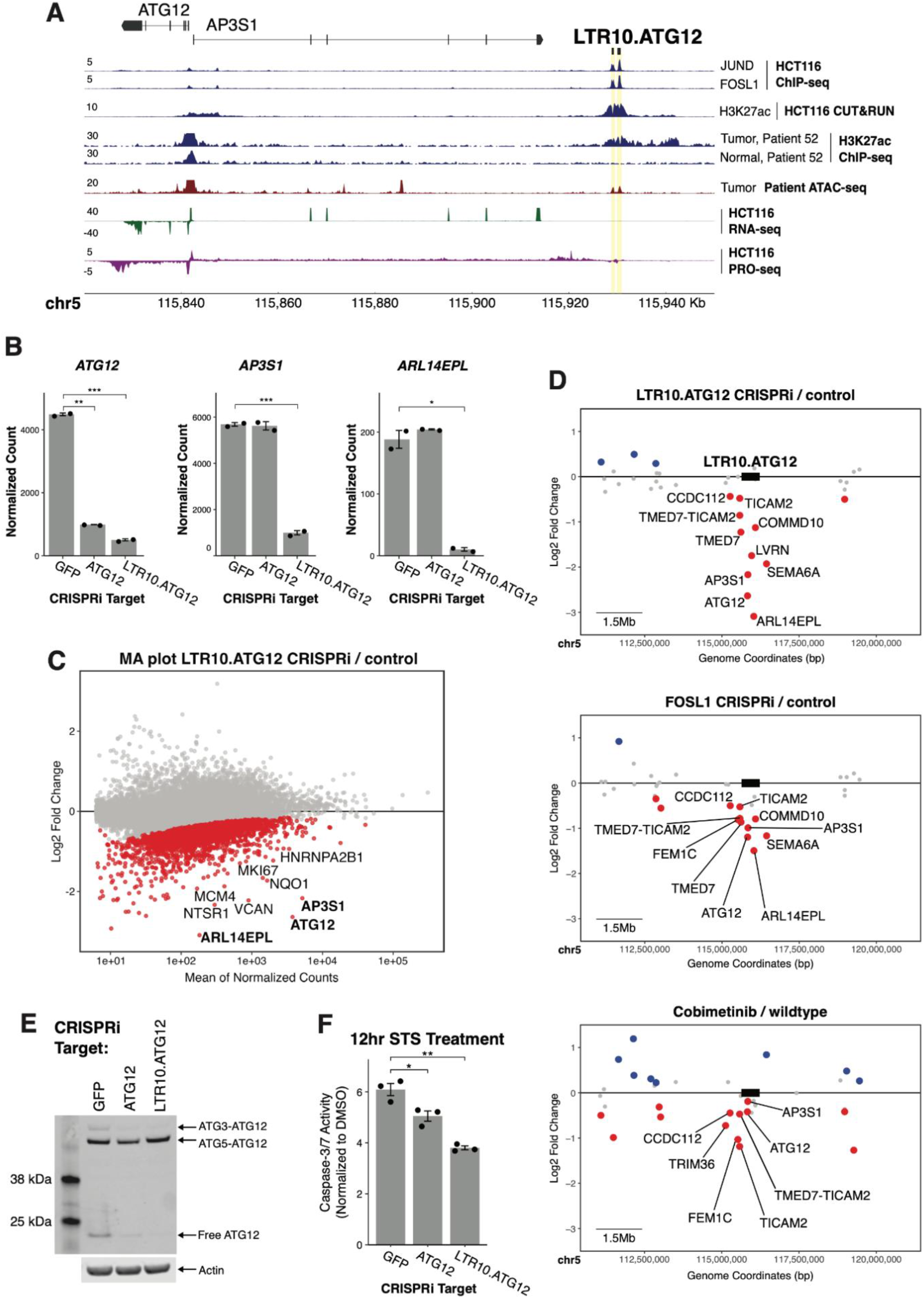
Functional characterization of LTR10.ATG12 in HCT116 cells. **(A)** Genome browser screenshot of the *ATG12/AP3S1* locus with the LTR10.ATG12 enhancer labeled. From top to bottom: JUND and FOSL1 ChIP-seq (GSE32465), H3K27ac CUT&RUN (in-house), H3K27ac ChIP-seq from matched tumor/normal samples from the CEMT Canadian Epigenome Project (patient AKCC52), tumor ATAC-seq from TCGA (patient COAD P022), HCT116 RNA-seq (in-house), and HCT116 PRO-seq (GSE129501). Axis numbers represent the upper limit of the range; the lower limit is always zero. **(B)** Normalized RNA-seq expression values of *ATG12, AP3S1*, and *ARL14EPL* in dCas9-KRAB-MeCP2 HCT116 cells stably transfected with gRNAs targeting the *ATG12* transcription start site (n=2), the LTR10.ATG12 element (n=2), or non-targeting (GFP) control (n=2). *: p < 0.05, **: p < 0.01, ***: p < 0.001, Welch’s t-test. Error bars denote SEM. **(C)** MA plot showing global gene expression changes in cells in response to silencing LTR10.ATG12. Significantly downregulated genes are shown in red. **(D)** Scatterplot of gene expression changes in the locus containing the LTR10.ATG12 element, associated with i) silencing LTR10.ATG12, ii) silencing *FOSL1*, or iii) cobimetinib treatment. Significantly downregulated genes are shown in red; significantly upregulated genes are shown in blue. Significantly downregulated genes located within 1.5 MB of the targeted element are labeled (element box not drawn to scale). **(E)** Immunoblot of endogenous ATG12 in each CRISPRi cell line. Different ATG12 conjugate forms are labeled. **(F)** Caspase-3/7 activity after 12 hrs staurosporine (STS) treatment, measured by the Caspase-Glo 3/7 assay. Treatments were performed in triplicate and signal for each cell line was normalized to signal from DMSO-treatment. *: p < 0.05, **: p < 0.01, Welch’s t-test. Error bars denote SEM.

**Figure 5:**
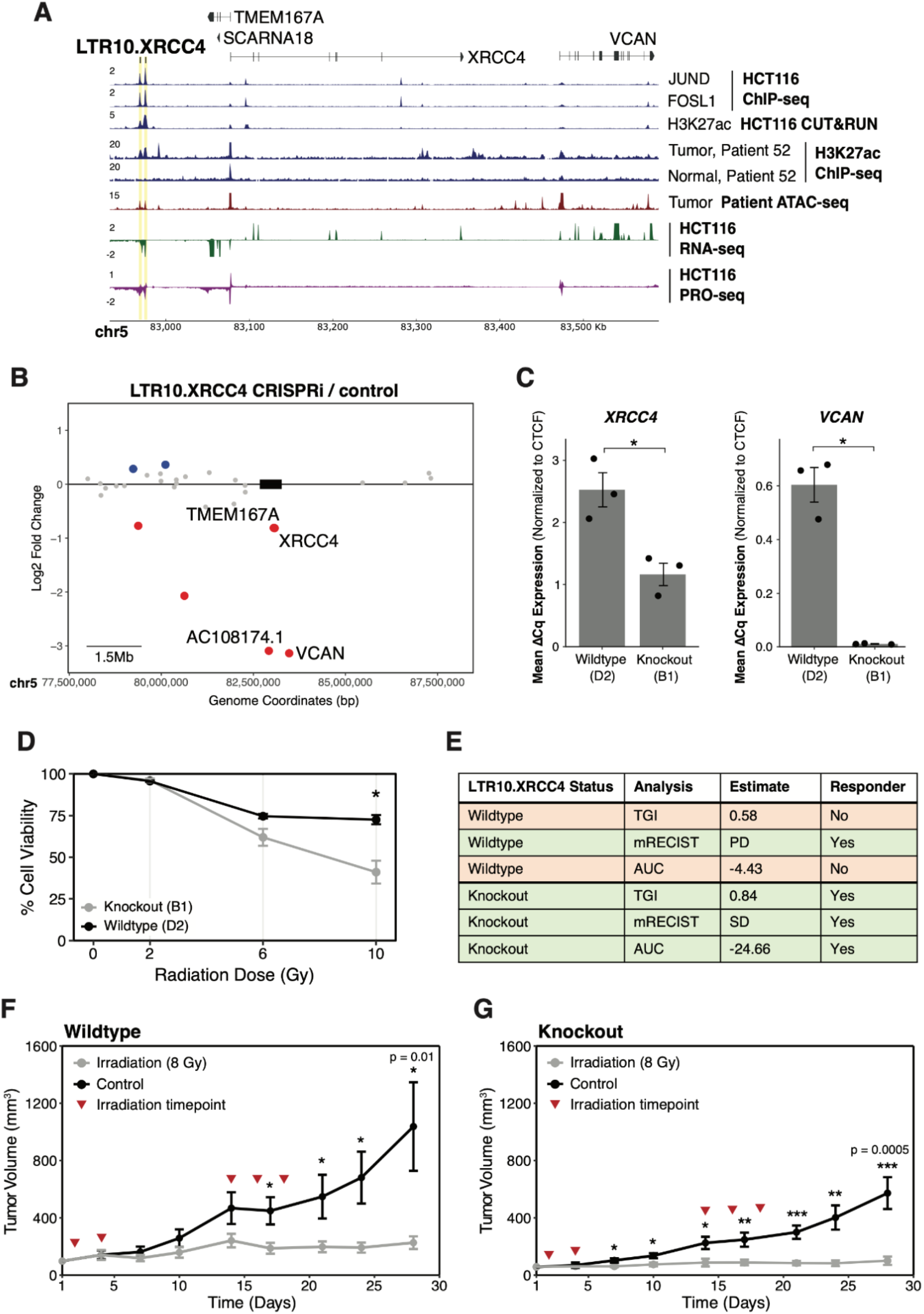
Functional characterization of LTR10.XRCC4 in HCT116 cells and xenograft models. **(A)** Genome browser screenshot of the *XRCC4* locus with the LTR10.XRCC4 enhancer labeled. From top to bottom: JUND and FOSL1 ChIP-seq (GSE32465), H3K27ac CUT&RUN (in-house), H3K27ac ChIP-seq from matched tumor/normal samples from the CEMT Canadian Epigenome Project (patient AKCC52), tumor ATAC-seq from TCGA (patient COAD P022), HCT116 RNA-seq (in-house), and HCT116 PRO-seq (GSE129501). Axis numbers represent the upper limit of the range; the lower limit is always zero. **(B)** Scatterplot of gene expression changes at the *XRCC4* locus after CRISPR silencing of the LTR10.XRCC4 enhancer. Significantly downregulated genes are shown in red; significantly upregulated genes are shown in blue. Significantly downregulated genes located within 1.5 MB of the targeted element are labeled (element box not drawn to scale). **(C)** RT-qPCR expression values of *XRCC4* and *VCAN* in wildtype HCT116 cells (n=3) and LTR10.XRCC4 knockout cells (n=3). *: p < 0.05, Welch’s t-test. Error bars denote SEM. **(D)** Dose-response curve showing cell viability in response to 0-10 Gy irradiation for LTR10.XRCC4 knockout and wildtype cells. Cell viability was measured by CellTiterGlo after 5 days and each replicate (n=2) was normalized to non-irradiated cells (n=2, averaged). *: p < 0.05, paired student’s t-test. Error bars denote SEM. **(E)** Classification of responder versus non-responder for both wildtype and LTR10.XRCC4 knockout cells, based on xenograft growth curves of untreated or irradiated mice. Three measures were calculated using Ortmann et al. (2021) ^69^: tumor growth inhibition (TGI), mRECIST, and area under the curve (AUC). **(F-G)** Average growth curves across replicates (n=9-10) for wildtype **(F)** versus LTR10.XRCC4 knockout **(G)** HCT116 xenograft tumors, with and without irradiation, for 28 days. 8 Gy treatment timepoints (days 2, 4, 14, 16, and 18) are indicated by red triangles. *: p < 0.05, **: p < 0.01, ***: p < 0.001, two sample t-test assuming equal variances. Error bars denote SEM. Individual growth curves are shown in Supp Fig S5E and S5F.

We focused on two of these LTR10-derived enhancers to explore their functional impact on tumor cells. We first investigated an enhancer that regulates *ATG12* (LTR10.ATG12), formed by two LTR10F elements on chromosome 5, located 87 kb from predicted target genes *ATG12* and *AP3S1* (Fig 4A). Silencing the LTR10.ATG12 enhancer resulted in downregulation of *ATG12* as well the neighboring gene *AP3S1* and eight other genes within 1.5 Mb (Fig 4B, 4C, 4D, Supp Table 11). As a separate control, we used CRISPRi to silence the *ATG12* promoter and found highly specific silencing of *ATG12* (Supp Fig S4K, S4L, Supp Table 17). These results indicate that the LTR10.ATG12 element functions as an enhancer that affects multiple genes in the locus. Genome-wide, we observed differential regulation of other genes, possibly due to indirect effects from target gene knockdown or off-target silencing of other LTR10 elements (Supp Fig 4M). Notably, we observed that multiple genes regulated by LTR10.ATG12 showed similar patterns of transcriptional downregulation in response to *FOSL1* silencing and cobimetinib treatment (Fig 4D). These results indicate that LTR10.ATG12 acts as an enhancer that controls AP1-dependent transcriptional activation of multiple genes in the *ATG12/AP3S1* locus in HCT116 cells.

The *ATG12* gene encodes a ubiquitin-like modifier required for macroautophagy as well as mitochondrial homeostasis and apoptosis ^49–52^. Expression of *ATG12* is associated with tumorigenesis and therapy resistance in colorectal and gastric cancer ^53,54^, but the mechanism of cancer-specific regulation of *ATG12* has not been characterized. Therefore, we aimed to determine whether the LTR10.ATG12 enhancer was responsible for regulating *ATG12* expression and activity in HCT116 cells. First, we validated that silencing the enhancer resulted in decreased ATG12 protein levels by immunoblotting (Fig 4E). In cells where either *ATG12* or the enhancer was silenced, there was a clear reduction in protein levels of both free ATG12 and the ATG3-ATG12 conjugate. There was minimal knockdown effect on the levels of the ATG5-ATG12 conjugate, which has previously been observed in *ATG12* silencing experiments and is due to the high stability of the ATG5-ATG12 complex^49^.

We tested whether ATG12-dependent functions require the activity of the LTR10.ATG12 enhancer. We treated each cell line with staurosporine (STS) to trigger mitochondrial apoptosis, which is dependent on free ATG12 binding to Bcl-2 ^50^. In cells where either *ATG12* or the enhancer was silenced, we observed significantly reduced caspase 3/7 activity, indicating defective mitochondrial apoptosis (Fig 4F). We did not detect differences in macroautophagy in cells treated with bafilomycin (Supp Fig S4N), consistent with the lack of knockdown of the ATG5-ATG12 conjugate ^52^. Our experimental results from silencing both *ATG12* and the enhancer are concordant with previous studies directly silencing *ATG12* using siRNAs in other cancer cell lines ^49,50^. Together, these experiments demonstrate that the LTR10.ATG12 enhancer is functionally important for ATG12-dependent activity in HCT116 cells.

We next focused on the LTR10.XRCC4 enhancer, which regulates *XRCC4* and *VCAN* based on our CRISPRi silencing experiment (Fig 5A, 5B, Supp Fig S5A, S5B, Supp Table 12). *XRCC4* is a DNA repair gene required for non-homologous end joining and promotes resistance to chemotherapy and radiation therapy ^55–59^. *VCAN* is an extracellular matrix protein that promotes tumor metastasis, invasion, and growth ^60–62^. Both *VCAN* and *XRCC4* have been reported to be regulated by MAPK/AP1 signaling in tumor cells ^56,63^, but the specific regulatory elements driving tumor-specific expression of these genes are unknown. We validated the enhancer activity of this element by generating cells harboring homozygous deletions using CRISPR (Supp Fig S5C, S5D) and confirmed that *XRCC4* and *VCAN* were significantly downregulated in edited cells (Fig 5C).

Previous studies have demonstrated that silencing or knocking out *XRCC4* directly causes increased sensitivity to DNA-damaging agents such as irradiation ^55,64,65^, including in HCT116 cells ^66^. To test whether the LTR10.XRCC4 enhancer regulates *XRCC4* function in cancer, we subjected control and knockout cells to 10 Gy irradiation, and found that knockout cells showed reduced viability following irradiation (Fig 5D), consistent with a previous study showing the role of XRCC4 in tumor cell survival following irradiation ^67^. We next tested how the deletion of LTR10.XRCC4 affects tumor response to irradiation in a mouse xenograft model. Irradiation inhibits the growth of tumors derived from HCT116 cells ^68^, therefore we tested whether reducing XRCC4 expression by deleting LTR10.XRCC4 affects tumor growth inhibition by irradiation.

We transplanted either control HCT116 cells or cells harboring a homozygous deletion of LTR10.XRCC4 into athymic nude mice, and subjected the mice to 8 Gy irradiation or mock irradiation. Tumors derived from both control and knockout cells showed growth inhibition in response to irradiation (Fig 5E-G, Supp Fig S5E-G, Supp Table 18). However, LTR10.XRCC4 knockout tumors showed more significant overall tumor growth inhibition by irradiation (Fig 5E), including at earlier timepoints (Fig 5F, 5G). No significant toxicities were seen in animal weights. While the difference seen in knockout tumors was modest, these results are consistent with previous studies in other cancers showing increased sensitivity to radiation in tumor xenografts when XRCC4 is silenced ^55^, and implicate a role for the LTR10.XRCC4 enhancer in regulating a clinically relevant tumor phenotype.

### LTR10 elements contain highly mutable VNTRs

Lastly, we investigated variation at LTR10 elements across 15,708 human genomes profiled by the Genome Aggregation database (gnomAD) ^70^. All LTR10 insertions are fixed, but we observed an unexpected enrichment of >10bp indel structural variants affecting the AP1 motif region specific to LTR10A and LTR10F, but not other LTR10 subfamilies such as LTR10C (Fig 6A). Further sequence inspection revealed that LTR10A and LTR10F elements contain an internal variable number of tandem repeats (VNTR) region, composed of a 28-30 bp sequence that includes the AP1 motif (Fig 6B, Supp Fig S6A). Individual LTR10 elements show a wide range of regulatory potential in HCT116 cells, as approximated by peak scores of H3K27ac CUT&RUN and FOSL1 ChIPseq (Supp Fig S6B, Supp Table 19), and demonstrated by the CRISPR-validated LTR10 enhancers. We speculate that the number of AP1 motifs within LTR10 elements may influence their regulatory potential. LTR10 elements annotated in the reference genome show extensive variation in tandem repeat length, with up to 33 copies of the AP1 motif (Supp Fig S6C, Supp Table 20). The number of motifs strongly correlates with H3K27ac and FOSL1 binding activity in HCT116 cells (Supp Fig S6C), suggesting that tandem repeat length affects AP1-dependent regulation of individual elements. Across the human population, LTR10A and LTR10F elements harbor many rare and common indel structural variants of lengths that follow a 28-30 bp periodicity, and this pattern is absent in LTR10C elements which lack the tandem repeat region (Fig 6C, 6D). These elevated levels of polymorphism across copies and individuals are characteristic of unstable tandem repeat regions ^71^, and suggest that LTR10 VNTR regions may be a common source of genomic regulatory variation.

**Figure 6:**
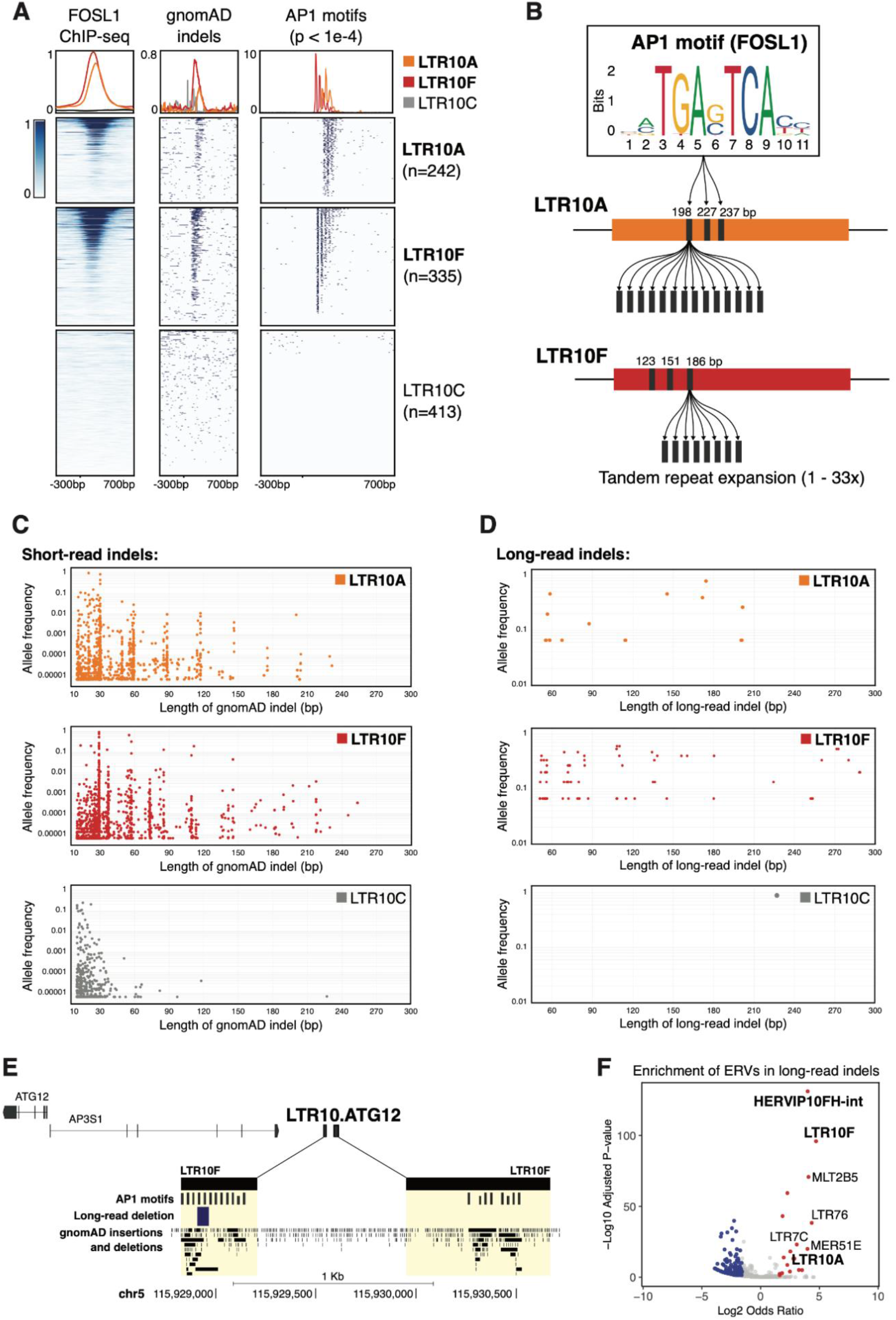
LTR10 repeat instability and polymorphism. **(A)** Heatmap of FOSL1 ChIP-Seq, gnomAD indels between 10-300bp in length, and AP1 motif matches (p<1e-4) across LTR10A, LTR10F, and LTR10C elements. Overlapping elements were removed, retaining 990 LTR10 elements total across the three subfamilies. FOSL1 ChIP-seq was obtained from GSE32465. **(B)** Schematic of variable number tandem repeat (VNTR) region within LTR10A and LTR10F elements. **(C)** Scatterplot of high-confidence gnomAD indels between 10-300 bp in length detected in LTR10A, LTR10F, or LTR10C subfamilies. Each indel is plotted by its length and allele frequency. **(D)** As in (C) but using long-read supported data. **(E)** Genome browser screenshot of LTR10.ATG12 showing AP1 motifs, long-read indels (58bp deletion reported by Quan et al., 2021), and gnomAD indels. **(F)** GIGGLE enrichment of ERVs within long-read indels. Significantly enriched ERVs are shown in red; significantly depleted ERVs are shown in blue.

Accurately genotyping tandem repeat length polymorphisms remains a major challenge using short-read data, therefore we validated the presence of LTR10 VNTR polymorphisms using structural variant calls generated from long-read whole-genome sequences from 15 individuals ^72^. We recovered indel structural variants within 24 distinct LTR10A and LTR10F elements, which also showed 28-30 bp periodicity (Fig 6D, Supp Fig S6D). We confirmed the presence of additional LTR10 VNTR indels using a separate long-read dataset from 25 Asian individuals (Fig 6E, Supp Fig S6E-J) ^73^. At the LTR10.ATG12 locus, we observed multiple indels supported by both short-read and long-read data that are predicted to affect AP1 motif copy number (Fig 6E, Supp Fig S6F). At a genome-wide level, LTR10 elements were a significantly enriched source of long-read indels, despite being fixed in the population (Fig 6F). Therefore, expansions or contractions within LTR10 VNTR regions are an underappreciated source of germline genetic variation that could underlie regulatory variation, consistent with polymorphisms recently reported in the VNTR region of SINE-VNTR-Alu (SVA) elements ^74^.

Finally, we searched for evidence of tumor-specific somatic expansions within LTR10 VNTR regions. We analyzed a long-read whole-genome sequencing dataset generated from matched colorectal tumor and normal tissues from 20 patients ^75^, using Sniffles2 ^76^ to identify tumor-specific repeat expansions within LTR10 VNTR regions. After manually inspecting reads at each locus, we found evidence for tumor-specific VNTR expansions at H3K27ac-marked LTR10 elements in five out of 20 patients (Supp Table 21). Three patients showed independent somatic expansions at the same LTR10A locus on chromosome 1 located near gene *GPR137B* (Supp Fig S6K, S6L, S6O), suggesting that this locus is prone to inter-individual variation at both the germline (Supp Fig S6D) and somatic level. We also found evidence of tumor-specific mosaic VNTR deletions in four patients (Supp Table 21). In fact, one patient with high microsatellite instability showed evidence of multiple tumor-specific LTR10 variants: a predicted LTR10A VNTR expansion over 11,600 bp in length (Supp Fig S6K, Supp Table 21), as well as two tumor-specific deletions at different LTR10F VNTR loci (Supp Fig S6Q, S6R). While a larger cohort would be necessary to determine if these expansions are enriched within tumors with microsatellite instability, these analyses provide evidence that LTR10 VNTRs are subject to tumor-specific somatic expansions and contractions, which can alter tumor-specific gene regulatory activity.

## DISCUSSION

Our study demonstrates that oncogenic MAPK/AP1 signaling drives global epigenetic and transcriptional activation of LTR10 elements in colorectal cancer and other epithelial cancers. A subset of these elements act as enhancers that facilitate pathological AP1-dependent transcriptional rewiring at multiple loci in cancer cells. Collectively, our data have several key implications for understanding how TEs shape cancer-specific regulatory networks.

First, our pan-cancer epigenomic analysis revealed multiple primate-specific ERV families that are enriched within tumor-specific accessible chromatin across all 21 solid tumor types profiled by TCGA ^27^. This implicates ERVs as a pervasive source of regulatory elements that shape gene regulation across most tumor types, expanding on recent studies that characterized TE-derived enhancers in prostate cancer ^26^ and acute myeloid leukemia ^25^ as well as other genomic studies profiling tumor-specific TE-derived enhancer activity in different cancers ^17,18,20,21,77^. We focused on LTR10 elements as a case example, which showed recurrent epigenomic signatures of enhancer activity in epithelial cancers including colorectal cancer. Both bulk and single-cell RNA-seq analysis of patient tumors revealed that LTR10 elements display tumor-specific transcriptional activation in a substantial fraction (∼30%) of cases. While our study found that LTR10 elements are normally repressed in adult somatic tissues and show largely tumor-specific enhancer activity, a recent study reported that many LTR10 elements also show enhancer activity in the developing human placenta ^78^, consistent with the hypothesis that reactivation of placental-specific gene regulatory networks may contribute to cancer pathogenesis ^79–81^.

Using CRISPR to silence or knock out individual elements in HCT116 colorectal cancer cells, we found that LTR10-derived enhancers causally drive AP1-dependent gene expression at multiple loci, including genes with established roles in tumorigenesis and therapy resistance including *ATG12*, *XRCC4*, and *VCAN* ^53,55,59–61,82–84^. While we focused on LTR10 elements predicted to regulate genes with established relevance to cancer, we also uncovered many elements that did not have predicted gene regulatory or functional consequences, indicating that LTR10 enhancer activity is not intrinsically pathological. Moreover, the regulatory activity of different LTR10-derived enhancers across the genome is likely to vary across individual tumors depending on the genetic and epigenetic background of the tumor and individual. Nevertheless, our findings support a model where LTR10-derived enhancers are important contributors to tumor-specific transcriptional dysregulation, which in some cases can significantly influence tumorigenesis and therapy resistance.

Second, our work shows that ERV-derived enhancers link oncogenic AP1/MAPK signaling to pathological transcriptional rewiring in colorectal cancer. Components of the MAPK pathway are frequently mutated in cancers, leading to oncogenic hyperactivation of MAPK signaling which promotes pathological gene expression and tumor cell proliferation ^43,85^. However, this process is poorly defined at the genomic level, and the specific regulatory elements that drive AP1-dependent transcriptional dysregulation have remained uncharted. Furthermore, inhibition of MAPK signaling is a common therapeutic strategy for many cancers ^86,87^ including colorectal cancer ^88,89^, but we have an incomplete understanding of how MAPK inhibition alters cancer epigenomes to achieve a therapeutic effect. Our study shows that oncogenic AP1/MAPK signaling results in activation of LTR10 enhancers, and treatment with a MAPK inhibitor effectively silences LTR10 regulatory activity in cancer cells. Therefore, the silencing of LTR10 ERV regulatory activity is an important but underappreciated mechanism underlying therapeutic MAPK inhibition.

Finally, we discovered that LTR10 elements are frequently affected by tandem repeat expansions that could influence their regulatory activity. Although all LTR10 insertions are fixed in the human population, they contain internal tandem repeats that show high levels of length polymorphism associated with repeat instability, consistent with a recent report of variable-length SVA elements which also contain internal tandem repeats ^74^. Germline or somatic variation in AP1 motif copy number within these elements may alter cancer-specific enhancer landscapes, and we found evidence that LTR10 VNTRs can be subject to somatic expansions or contractions in cancer cells with microsatellite instability ^90^. Our study of LTR10 highlights how TEs that are normally silenced can become reactivated in cancer and cause aberrant gene expression. For elements that promote pathogenesis, their restricted activity in age-associated diseases like cancer may result in reduced or nearly neutral fitness consequences. Therefore, the accumulation of TEs subject to epigenetic silencing may be a fundamental process that shapes cancer-specific gene regulatory networks.

## METHODS

### Cell culture

The HCT116 cell line was purchased from ATCC and cultured in McCoy’s 5A medium supplemented with 10% FBS and 1% penicillin/streptomycin (Gibco). Cells were cultured at 37°C in 5% carbon dioxide. Transfections were performed using FuGENE (Promega). For treatments modulating MAPK signaling, HCT116 cells were untreated or treated for 24 hrs with 1 uM Cobimetinib, 100 ng/mL TNF alpha, or DMSO.

### CRISPR-mediated silencing and knockout of LTR10s

For CRISPR-mediated silencing (e.g., CRISPRi) of select LTR10 elements and gene transcription start sites (TSS), a HCT116 dCas9-KRAB-MeCP2 stable line was first generated using the PiggyBac system (System Bioscience). The PiggyBac Donor plasmid, PB-CAGGS-dCas9-KRAB-MeCP2 was co-transfected with the Super PiggyBac transposase expression vector (SPBT) into HCT116 cells. The pB-CAGGS-dCas9-KRAB-MeCP2 construct was a gift from Alejandro Chavez & George Church (Addgene plasmid # 110824). 24 hours post-transfection, cells were treated with Blasticidin to select for integration of the dCas9 expression cassette, and selection was maintained for 10 days. CRISPR gRNAs specific to the DNA elements of interest (i.e., 0 predicted off target sequences) were selected using pre-computed CRISPR target guides available on the UCSC Genome Browser hg38 assembly, and complementary oligos were synthesized by Integrated DNA Technologies. Complementary oligos were designed to generate BstXI and BlpI overhangs for cloning into PB-CRISPRia, a custom PiggyBac CRISPR gRNA expression plasmid based on the lentiviral construct pCRISPRia (a gift from Jonathan Weissman, Addgene plasmid # 84832). Complementary gRNA-containing oligos were hybridized and phosphorylated in a single reaction, then ligated into a PB-CRISPRia expression plasmid linearized with BstXI and BlpI (New England Biolabs). Chemically competent Stable E. Coli (New England Biolabs) was transformed with 2 uL of each ligation reaction and resulting colonies were selected for plasmid DNA isolation using the ZymoPure Plasmid miniprep kit (Zymo Research). Each cloned gRNA sequence-containing PB-CRISPRia plasmid was verified by Sanger sequencing (Quintara Bio).

To generate CRISPRi stable lines, PB-CRISPRia gRNA plasmids were co-transfected with the PiggyBac transposase vector into the HCT116 dCas9-KRAB-MeCP2 polyclonal stable line. The following number of uniquely-mapping gRNA plasmids were designed per target based on the pre-computed UCSC hg38 CRISPR target track: ATG12 (1), GFP (1), FOSL1 (1), LTR10.ATG12 (4), LTR10.FGF2 (2), LTR10.MCPH1 (3), LTR10.MEF2D (2), LTR10.XRCC4 (2). The same total amount of gRNA plasmid was used for transfections involving one or multiple gRNAs. 24 hours post-transfection, cells were treated with Puromycin to select for integration of the sgRNA expression cassette(s). Selection was maintained for 5 days prior to transcriptional analyses.

For CRISPR-mediated knockout of LTR10.KDM6A, 2 gRNAs (1 specific to each flank of the element) were identified and synthesized as sgRNAs by IDT. For CRISPR-mediated knockout of LTR10.XRCC4, 4 gRNAs (two specific to each flank of the element) were identified and synthesized as sgRNAs by IDT. Using IDT’s AltR technology, RNP complexes were generated in vitro, and electroporated into HCT116 cells using the Neon system (ThermoFisher Scientific). Clonal lines were isolated using the array dilution method in a 96-well plate format, and single clones were identified and screened for homozygous deletions by PCR using both flanking and internal primer pairs at the expected deletion site. gRNAs and PCR primers for each candidate are provided in Supp Table 22.

### Cell autophagy and apoptosis assays

For assaying mitochondrial apoptosis, HCT116 CRISPRi cell lines were treated for 12 hours with Staurosporine (STS) at 0.5 uM or DMSO (vehicle) followed by measurement of Caspase activity via the Caspase-Glo 3/7 assay (Promega). Results are representative of at least 3 independent experiments. For assaying autophagy, HCT116 CRISPRi cell lines were untreated or treated with Bafilomycin A at 10 nM or 100 nM for 6 hrs and 18 hrs, followed by LC3B Western blotting. Results are representative of at least 3 independent experiments.

### Western blots

For ATG12 Western blots, cell lysates were prepared with MPER buffer (ThermoFisher Scientific). For LC3B Western blots, cell lysates were prepared with RIPA buffer. All cell Lysates were resuspended in 4x NuPage LDS Sample buffer containing reducing agent (ThermoFisher Scientific). For ATG12 Western blots, total protein was concentrated and size-selected by passing through an Amicon Ultra 10K column (Millipore), retaining the high molecular weight fraction, and 40 ug of protein was loaded per lane. For LC3B Western blots, 2 ug total protein was loaded per lane. Antibodies used were as follows: ATG12 (cat #4180T, Cell Signaling Technologies), Beta-Actin (cat # 3700T, Cell Signaling Technologies), LC3B (cat # NB100-2220, Novus Biologicals). Results are representative of at least 3 independent experiments.

### Luciferase assay

Reporter assays were conducted in HCT116 cells using the secreted NanoLuc enhancer activity reporter pNL3.3 (Promega) and normalized against a constitutively active firefly luciferase reporter vector, pGL4.50 (Promega). LTR10 consensus sequences for subfamilies LTR10A and LTR10F were downloaded from Dfam. AP1 motifs within LTR10A and LTR10F were shuffled as follows: LTR10A (first two motifs): cctgagtcacc to cagccccgtta; LTR10A (third motif): cttagtcacc to cagtttaccc; LTR10F (all three motifs): cctgactcatt to cgtatccttac. Sequences are provided in Supp Table 22. Due to their high repeat content, consensus sequences were synthesized as multiple fragments (Integrated DNA Technologies, Twist BioScience) and then assembled into pNL3.3 enhancer reporter plasmids using Gibson Assembly (New England Biolabs). Each cloned reporter plasmid was verified by Sanger sequencing (Quintara Bio). To assay reporter activity, HCT116 cells were transfected with a reporter construct as well as the pGL4.50 construct constitutively expressing firefly luciferase. 24 hrs after transfection, media was replaced with media containing 1 uM Cobimetinib, 100 ng/mL TNF alpha, or DMSO (vehicle). 24 hours following treatment, luminescence was measured using the NanoGlo Dual Luciferase Reporter Assay System (Promega). All experiments were performed with 3 treatment replicates per condition in a 96-well plate format. Luminescence readings were first normalized to firefly co-transfection controls, then presented as fold-change against cells transfected with an empty minimal promoter pNL3.3 vector as a negative control. Results are representative of at least 3 independent experiments. Barplots are presented as mean +/- s.d.

### Irradiation experiment

HCT116 control or knockout cells were irradiated using a Faxitron irradiator (Model RX-650) at 0, 2, 6 or 10 Gy then left to recover for up to 5 days. Cell viability was measured by CellTiter-Glo luminescence assay (Promega). Two replicates (each based on the average of three CellTiter-Glo readings) were normalized to unirradiated (0 Gy) as a control.

### Mouse xenograft experiment

All experiments were approved by the Institutional Animal Care and Use Committee of the University of Colorado Anschutz Medical Campus and conducted in accordance with the National Institutes of Health Guidelines for the Care and Use of Laboratory Animals. Female athymic nude mice (aged 15-16 weeks at start of study) were purchased from Envigo (Indianapolis, IN) and implanted subcutaneously on the hind flanks with 2.5 million cells in 100 ul of either HCT116 wildtype or LTR10.XRCC4 CRISPR knockout cells under isoflurane anesthesia with a 23 ga ½ needle. The cell solution injected consisted of 1:1 ratio of RMPI media and cultrex (Cultrex PathClear BME, Type 3 from Bio-Techne). We injected wildtype or knockout cells into 40 mice (20 each, one side per mouse), then mice were randomized into treatment groups (20 irradiated, 20 mock) and treatments were initiated when the average tumor volume reached between 50-100 mm^3^. Tumor volume was calculated by {(width^2^) x length} x 0.52. Irradiation treatment consisted of 8 Gy x 3 fractions on days 2, 4, 14, 16, and 18. Tumor measurements were taken twice weekly using digital calipers, and toxicity was monitored by measuring body weight twice weekly and the study ended at 28 days. Tumor growth inhibition was measured using KuLGaP ^69^.

### RNA-seq

Sequencing libraries were prepared from RNA harvested from treatment or transfection replicates. Total RNA was extracted using the Quick-RNA Miniprep Plus Kit (Zymo Research). PolyA enrichment and library preparation was performed using the KAPA BioSystems mRNA HyperPrep Kit according to the manufacturer’s protocols. Briefly, 500 ng of RNA was used as input, and KAPA BioSystems single-index or unique dual-index adapters were added at a final concentration of 7 nM. Purified, adapter-ligated library was amplified for a total of 11 cycles following the manufacturer’s protocol. The final libraries were pooled and sequenced on an Illumina NovaSeq 6000 (University of Colorado Genomics Core) as 150 bp paired-end reads.

### CUT&RUN

Libraries were prepared from treatment replicates. Approximately 5x10^5^ viable cells were used for each CUT&RUN reaction, and pulldowns were generated following the protocol from ^91^. All buffers were prepared according to the “High Ca^2+^/Low Salt” method using digitonin at a final concentration of 0.05%. The following antibodies were used at the noted dilutions: rabbit anti-mouse IgG (1:100), rabbit anti-H3K27ac (1:100). pA-MNase (gift from Steven Henikoff) was added to each sample following primary antibody incubation at a final concentration of 700 ng/mL. Chromatin digestion, release, and extraction was carried out according to the standard protocol. Sequencing libraries were generated using the KAPA BioSystems HyperPrep Kit according to the manufacturer’s protocol with the following modifications: Freshly diluted KAPA BioSystems single-index adapters were added to each library at a final concentration of 9 nM. Adapter-ligated libraries underwent a double-sided 0.8X/1.0X cleanup using KAPA BioSystems Pure Beads. Purified, adapter-ligated libraries were amplified using the following PCR cycling conditions: 45 s at 98°C, 14x(15 s at 98°C, 10 s at 60°C), 60 s at 72°C. Amplified libraries underwent two 1X cleanups using Pure Beads. The final libraries were quantified using Qubit dsDNA High Sensitivity and TapeStation 4200 HSD5000. Libraries were pooled and sequenced on an Illumina NovaSeq 6000 (University of Colorado Genomics Core) as 150 bp paired-end reads.

### Processing of sequencing data

Reads obtained from our own datasets and from published studies were reprocessed using a uniform analysis pipeline. FASTQ reads were evaluated using FastQC (v0.11.8) and MultiQC (v1.7), then trimmed using BBDuk/BBMap (v38.05). For ATAC-seq, ChIP-seq, and CUT&RUN datasets, reads were aligned to the hg38 human genome using BWA (v0.7.15) and filtered for uniquely mapping reads (MAPQ > 10) with samtools (v1.10). ChIP-Seq and ATAC-seq peak calls and normalized signal coverage bigwig plots were generated using MACS2 (v2.1.1) (with setting --SPMR). CUT&RUN peak calls were generated using MACS2 in paired-end mode using a relaxed p-value threshold without background normalization (-- format BAMPE --pvalue 0.01 --SPMR -B --call-summits). MACS2 was also run in single-end mode with additional parameters --shift -75 and --extsize 150, and Bedtools (v2.28.0) was used to merge peaks across the two modes of peak calling for each sample (bedtools merge with options -c 5 -o max).

RNA-seq and PRO-seq reads were aligned to hg38 using hisat2 (v2.1.0) with option --no-softclip and filtered for uniquely mapping reads with samtools for MAPQ > 10. Bigwig tracks were generated using the bamCoverage function of deepTools (v3.0.1), with CPM normalization (ignoring chrX and chrM) and bin size 1bp. Some datasets from TCGA, ENCODE, Cistrome DB and the CEMT Canadian Epigenomes Project were downloaded as post-processed peaks and bigwig files.

### TE colocalization analysis

To determine TE subfamily enrichment within regulatory regions, we used GIGGLE (v0.6.3) ^92^ to generate a genomic interval index of all TE subfamilies in the hg38 human genome, based on Dfam v2.0 repeat annotation (n=1315 TE subfamilies). Regulatory regions (e.g., ATAC, ChIP-Seq, or CUT&RUN peaks) were queried against the TE interval index using the GIGGLE search function (-g 3209286105 -s). Results were ranked by GIGGLE enrichment score, which is a composite of the product of −log10(P value) and log2(odds ratio). Significantly enriched TE subfamilies were defined as those with at least 25 overlaps between TE copies and a set of peak regions, odds ratio over 10, and GIGGLE score over 100 in at least one cancer type.

### Defining cancer-specific regulatory elements

To define cancer-specific regulatory elements, we first obtained aggregate ATAC-seq regions associated with each tumor type profiled by The Cancer Genome Atlas ^93^, which represent a union of recurrent ATAC-seq regions associated with each tumor type. Next, we identified regulatory regions in healthy adult tissues based on chromHMM regulatory regions defined by the Roadmap project. We used healthy adult tissues from categories 1_TssA, 6_EnhG & 7_Enh. We did not include fetal tissues (e.g. placental tissues, embryonic stem cells, trophoblast stem cells) in our set of Roadmap healthy regulatory regions, due to the high levels of basal ERV regulatory activity in these tissues. Finally, cancer-specific regulatory regions were defined using the subtract function of bedtools (option -A) to subtract Roadmap “healthy adult” regulatory regions from each cancer peak set.

### Transcription factor motif analyses

Motif analysis of LTR10 elements was performed using the MEME suite (v5.1.0) in differential enrichment mode ^94^. Entire LTR10 sequences were used for the motif analysis. HCT116 CUT&RUN H3K27ac-marked LTR10A/F sequences (n=144) were used as input against a background set of unmarked LTR10A/F sequences (n=561), with default settings other than the number of motif repetitions (Any) and the number of motifs to find (5). Each discovered motif was searched for similarity to known motifs using the JASPAR 2018 non-redundant DNA database with TomTom (v5.1.0). FIMO (v5.1.0) was then used to extract motif frequency and hg38 genomic coordinates, with p-value threshold set to 1e-4.

Motif analysis of cancer-specific ATAC-seq peaks from 21 TCGA cancer types was likewise performed using the MEME suite (v5.1.0) ^94^. Cancer-specific peaks for each cancer were defined by subtracting away Roadmap regulatory regions from each cancer peak set, as described in the previous section. The number of cancer-specific peaks for each cancer were as follows: ACC (n=8123), BLCA (n=13737), BRCA (n=30494), CESC (n=2449), CHOL (n=3012), COAD (n=9370), ESCA (n=12538), GBM (n=4114), HNSC (n=9441, KIRC (n=4807), KIRP (n=12315), LGG (n=3673), LIHC (n=8469), LUAD (n=16862), LUSC (n=15143), MESO (n=5275), PCPG (n=7891), PRAD (n=12130), SKCM (n=13710), STAD (n=11222), THCA (n=9991). Bedtools (v2.28.0) getfasta was used to convert the BED format peak files to FASTA format, and all nucleotides were converted to uppercase letters. MEME-ChIP (v5.1.0) was then run on each cancer-specific FASTA file, with settings -ccut 100 (maximum size of a sequence before it is cut down to a centered section), -order 1 (to set the order of the Markov background model that is generated from the sequences), -meme-mod anr (to allow any number of motif repetitions), -meme-minw 6 (minimum motif width), - meme-maxw 20 (maximum motif width), -meme-nmotifs 10 (maximum number of motifs to find), and the JASPAR 2018 non-redundant motif database. The output from CentriMo was used to obtain the AP1 motif p-value for each cancer type (i.e. adjusted p-value for motif id MA0477.1, alt id FOSL1).

### Differential analysis using DESeq2

For RNA-seq samples, gene count tables were generated using featureCounts from the subread (v1.6.2) package with the GENCODE v34 annotation gtf to estimate counts at the gene level, over each exon (including -p to count fragments instead of reads for paired-end reads; -O to assign reads to their overlapping meta-features; -s 2 to specify reverse-strandedness; -t exon to specify the feature type; -g gene_id to specify the attribute type).

To quantify TE expression at the subfamily level, RNA-seq samples were first re-aligned to hg38 using hisat2 with -k 100 to allow multi-mapping reads and --no-softclip to disable soft-clipping of reads. TEtranscripts (v2.1.4) was then used in multi-mapping mode with the GENCODE v34 annotation gtf and hg38 GENCODE TE gtf to assign count values to both genes and TE elements.

For H3K27ac CUT&RUN samples, bedtools multicov was used to generate a count table of the number of aligned reads that overlap MACS2-defined peak regions. The top 20,000 peaks were extracted from each sample and merged (using bedtools merge with -d 100) to produce the peak file used as input to bedtools multicov.

All count tables were processed with DEseq2 (v1.32.0). Normalized count values were calculated using the default DEseq2 transformation. R packages ggplot2 (v3.3.2), ggrepel (v0.8.2) and apeglm (v1.8.0) were used to visualize differentially expressed genes and TEs. The same DEseq2 analyses were used to identify differentially enriched peak regions between H3K27ac CUT&RUN samples (e.g. in response to MAPK treatment). Significantly differentially enriched regions were queried against the GIGGLE index of human repeats to identify over-represented TE subfamilies.

### Reanalysis of patient-derived bulk RNA-seq tumor/normal colon datasets

BAM files of matched tumor/normal RNA-seq datasets from 38 de-identified patients with colon adenocarcinomas were downloaded from TCGA-COAD using the GDC Data Transfer Client with a restricted access token. Each patient had one normal colon sample and one colorectal tumor sample. Gene and TE counts were assigned using TEtranscripts (v2.1.4) in multi-mapping mode, as above, with the GENCODE v34 annotation gtf and hg38 GENCODE TE gtf. Count tables were processed using DEseq2 (v1.32.0) and normalized count values were calculated using the multi-factor DEseq2 design of ∼Patient.ID+Condition, where Condition was either Primary Tumor or Solid Normal Tissue. Potential outliers were identified using principal component analysis based on gene counts (e.g. see Supp Fig S2F), but all samples were retained for downstream analysis. R packages ggplot2 (v3.3.2), ggrepel (v0.8.2) and apeglm (v1.8.0) were used to visualize differentially expressed genes and TEs.

Similarly, in order to perform correlative studies between LTR10 activity and tumor mutations or patient survival rates, RNA-seq BAM files from 358 patient-derived tumor samples were obtained from TCGA-COAD controlled access data. The steps above were repeated for each tumor sample to quantify transcription of LTR10 subfamilies. KRAS mutation status and survival status for each patient were derived from the TCGA-COAD patient metadata.

### Reanalysis of patient-derived single cell RNA-seq tumor/normal colon datasets

Single cell RNA-seq datasets of matched tumor/normal colon from 36 de-identified patients with colon adenocarcinomas from Pelka et al (2021) were downloaded using dbGaP controlled access (phs002407.v1.p1). Only patients with both tumor and adjacent normal tissue were analyzed (n=36). Raw FASTQ files for each sample were renamed according to the required Cell Ranger format, then processed with Cell Ranger (v7.0.0) count function using default parameters and the Cell Ranger transcriptome for the human reference genome (refdata-gex-GRCh38-2020-A). The resulting BAM files were filtered to remove lines without cell barcodes using samtools (v1.10). scTE (v1.0) was used to remap reads to both genes and TEs, using the provided hg38 index and default parameters except for -p 8 (number of threads to use), --hdf5 True (to save the output as a .h5ad formatted file), and - CB CB -UMI UB (to specify that the BAM file was generated by Cell Ranger, with cell barcodes and UMI integrated into the read ’CB:Z’ or ’UB:Z’ tag).

Output h5ad files were processed using Scanpy (v1.9.1) in a customized scRNA-seq workflow. Each patient was processed separately. Cell barcodes were excluded if they satisfied any of the following criteria: (1) fewer than 1200 reads, (2) fewer than 750 genes, (3) more than 25% of UMIs mapping to the mitochondrial genome. Genes and TEs were excluded if their expression level was deemed “undetectable”, i.e. at least two cells had to contain at least 5 reads from the gene/TE. Tumor and normal samples from the same patient were merged after filtering and quality control, retaining the tissue of origin (T vs N) information.

For each patient, the filtered and merged data was normalized to 10,000 reads per cell, log-transformed, and then clustered. Dimensionality reduction was performed using principal component analysis (log=True, n_pcs=40), tSNE (perplexity=30, learning_rate=1000, random_state=0, n_pcs=40), and UMAP (n_neighbors=30, n_pcs=40, min_dist=0.8, spread=1, random_state=0, maxiter=100). Leiden clustering (resolution=0.75) was used to assign cells to clusters, and cell clusters with less than 20 cells were excluded from final UMAP visualizations. Cell types were annotated using the PanglaoDB database ^95^ of gene expression markers, with manual verification.

### Prediction of LTR10 enhancer gene targets

LTR10 elements were initially prioritized for CRISPR silencing or deletion based on enhancer predictions from the Activity-by-Contact (ABC) model ^48^. Publicly available HCT116 ATACseq (GEO accession GSM3593802) and in-house HCT116 H3K27ac CUT&RUN were used as input to the ABC pipeline, as well as the provided averaged human cell line HiC file. Predicted enhancer regions with an ABC interaction score over 0.001 were intersected with H3K27ac-marked LTR10A/F elements. Putative LTR10 enhancers were then checked for proximity (e.g. within 1.5Mb) to FOSL1-regulated genes (i.e. genes that were significantly downregulated by FOSL1 knockdown), or MAPK-regulated genes (i.e. genes that were significantly affected by MAPK treatments Cobimetinib and TNF-alpha, based on inhouse RNAseq).

### Evolutionary analysis of LTR10 sequences

Genomic coordinates of LTR10 elements in the hg38 human genome were obtained from Dfam v2.0, based on RepeatMasker v4.0.6 repeat annotation. The nucleotide sequence of each LTR10 element was extracted using the getfasta function from bedtools (using -name+ to include coordinates in the header and -s for strand specificity). VSEARCH (v2.14.1) was used to set a minimum length threshold of 200bp for LTR10 sequences (-sortbylength - minseqlength 200), prior to alignment. MUSCLE (v3.8.1551) was used to align the remaining sequences. Jalview (v2.11.1.4) was used to perform a principal component analysis on pairwise similarity scores derived from the multiple sequence alignment.

To confirm that LTR10 elements can be uniquely mapped, all individual LTR10A/F sequences were clustered at 99% identity (-qmask none -id 0.99) with VSEARCH (v2.14.1). No clusters contained more than one sequence, indicating that no identical LTR10A/F copies exist within the human genome.

LTR10 consensus sequences representing each LTR10 subfamily (A-G) were downloaded from Dfam v2.0. Sequences were concatenated into one FASTA file and aligned using MUSCLE. FastTree was used to infer a maximum likelihood phylogeny representing the LTR10 subfamily relationships.

The phylogeny of known primate relationships was obtained from TimeTree ^96^ and the HERV-I insertion estimate was confirmed based on the presence or absence of LTR10 sequences across mammals, using BLAST (v2.7.1+) ^97^.

### VNTR identification

gnomAD (v3.1) VCF files for each hg38 chromosome were filtered for high-confidence indels (FILTER=PASS) using the query function of bcftools (v1.8) with format parameter - f’%CHROM\t%POS0\t%END\t%ID\t%REF\t%ALT\t%AF\t%TYPE\tFILTER=%FILTER\n’. The remaining indels were then subsetted by size to retain insertions or deletions between 10 to 300bp in length. Chromosome VCFs were concatenated into one whole genome BED file. Bedtools (v2.28.0) was used to intersect the indel BED file with LTR10 elements from each subfamily, based on Dfam (v2.0) repeat annotation.

Indels from additional short- and long-read datasets were likewise filtered by variant type (INS or DEL) and indel length (10-300bp for short reads; 50-300bp for long reads, since the minimum length reported by long-read SV callers is 50bp). Filtered VCFs were then intersected with LTR10 elements using bedtools (v2.28.0). Deletion length versus allele frequency was plotted for each subfamily, for each separate dataset. VNTR regions within LTR10 elements were also intersected with GTEx v8 fine-mapped CAVIAR and DAP-G cis-eQTL files ^98^, again using bedtools (v2.28.0).

To identify tumor-specific VNTR expansions or contractions, we downloaded a long-read whole-genome nanopore sequencing dataset generated from matched tumor/normal tissues from 20 patients with advanced colorectal adenocarcionomas ^75^. For each sample, we used minimap2 ^99^ (v2.22-r1101) to align reads to the hg38 reference genome, with parameters - a to generate output in SAM format, -x map-ont to specify nanopore input reads, -t 4 to set the number of threads to 4, and -Y to use soft clipping for supplementary alignments. We then samtools (v1.10) to generate sorted BAM files, with commands samtools view -bS to convert from SAM to BAM format, samtools sort (default parameters) to sort reads by coordinate, and samtools index (default parameters) to generate a BAM index file for each BAM. We then used Sniffles2 (v2.0.7) ^76^ to identify tumor-specific SVs within LTR10 VNTR regions. For each tumor/normal pair, we called SVs using both the default parameters (optimized for germline variants), and then again using the --non-germline parameter for the tumor sample only (optimized for detecting low frequency or mosaic variants). The reference genome was set to hg38 and --tandem-repeats were annotated using the Sniffles-provided human_GRCh38_no_alt_analysis_set.trf.bed file. Sniffles was run with the -snf option to save candidate SVs to the SNF binary file, per sample. For each patient, tumor and normal SNF files were then merged using the Sniffles population verge with --vcf to specify VCF output format. All VCF output files were intersected with LTR10 VNTR regions using bedtools (v2.28.0). For each patient, tumor-specific variants were extracted using the SUPP_VEC tag in the INFO field of the output VCFs (i.e. by extracting all SVs with SUPP_VEC=01, which signifies absence in the normal sample and presence in the tumor sample). Finally, for each called insertion or deletion, we manually inspected aligned reads using the UCSC genome browser to confirm differences between the tumor and normal samples.

## DATA AVAILABILITY

High-throughput sequencing data (RNA-seq, CUT&RUN) has been deposited in the Gene Expression Omnibus (GEO) with the accession code GSE186619. GSE IDs of public datasets used in this study are listed in the figure legends and GitHub repository. The following databases were also used: Cistrome DB (downloaded Feb 2019), Roadmap/ENCODE (downloaded Feb 2019), The Cancer Genome Atlas (downloaded Sept 2019), CEMT Canadian Epigenome Project (downloaded July 2020), Dfam 2.0 and gnomAD v3.1.

## CODE AVAILABILITY

Source code and workflows are available on GitHub: https://github.com/atmaivancevic/ERV_cancer_enhancers

## COMPETING INTEREST STATEMENT

The authors declare no competing interests.

## ACKNOWLEDGEMENTS

We thank the University of Colorado Genomics Shared Resource and BioFrontiers Computing core for technical support during this study. We thank Ben Nebenfuehr and Nausica Arnoult for assistance with the cell irradiation experiments, and Marilyn Jackson, Cameron Binns, Stephen Smoots and Adrian Dominguez for assistance with the animal studies. A.I. was supported by the National Cancer Center. E.B.C. was supported by the National Institutes of Health (1R35GM128822), the Alfred P. Sloan Foundation, the David and Lucile Packard Foundation, and the Boettcher foundation.

## Author contributions

A.I. designed and conducted the bioinformatics analyses. D.M.S. and O.J. designed and conducted the CRISPR and cell irradiation experiments. S.M.B and T.M.P designed and conducted the mouse xenograft experiments. L.L.N. performed RNA extractions from colorectal cancer cell lines. A.I. and E.B.C. conceived the study, analyzed the data, and wrote the paper. B.G.B supervised the study and helped write the paper.

## Notes

### Competing Interest Statement

The authors have declared no competing interest.

### Summary of Updates

Manuscript and supplementary updated to include new in vitro and in vivo experiments on the LTR10.XRCC4 enhancer that regulates DNA repair gene XRCC4; new analysis of LTR10 transcriptional activity using patient-derived bulk RNA-seq and single cell RNA-seq tumor/normal datasets; and new analysis of tumor-specific somatic expansions of LTR10 VNTR regions in colorectal cancer patients.

