## Supplementary Figures for "Endogenous retroviruses mediate transcriptional rewiring in response to oncogenic signaling in colorectal cancer"

### LIST OF SUPPLEMENTARY MATERIAL

**Supplementary Figure S1:** Additional figures for the pan-cancer epigenomic analysis of TE activity.

**Supplementary Figure S2:** Additional figures for the regulatory activity of LTR10 regulatory activity in tumor and normal cells.

**Supplementary Figure S3:** Additional figures for the control of LTR10 regulatory activity by AP1/MAPK signaling.

**Supplementary Figure S4:** Additional figures for the CRISPR silencing and deletion of predicted LTR10 enhancers.

**Supplementary Figure S5:** Additional figures for the functional characterization of enhancer LTR10.XRCC4 in HCT116 cells and xenograft models.

**Supplementary Figure S6:** Additional figures showing LTR10 repeat instability and polymorphism.

**Supplementary Table 1:** Enrichment scores for TEs in tumor accessible chromatin from TCGA.

**Supplementary Table 2:** Enrichment scores for LTR10A/F elements in regulatory regions defined by Roadmap.

**Supplementary Table 3:** Enrichment scores for LTR10A/F elements in human transcription factor datasets generated by Cistrome DB.

**Supplementary Table 4:** Enrichment scores for KRAB zinc finger protein ZNF561/562 ChIP-seq peaks in repetitive regions from different repeat subfamilies.

**Supplementary Table 5:** DEseq2 results for matched tumor/normal bulk RNA-seq samples from 38 colorectal cancer patients from TCGA.

**Supplementary Table 6:** Wilcoxon rank-sum test results for FOSL1, LTR10A and LTR10F in tumor vs normal epithelia. Single cell RNA-seq samples from 36 colorectal cancer patients obtained from Pelka et al (2021).

**Supplementary Table 7:** DEseq2 results for 358 tumor bulk RNA-seq samples from 358 colorectal cancer patients with KRAS mutation status from TCGA.

**Supplementary Table 8:** DEseq2 results for CRISPR silencing of AP1 component FOSL1.

**Supplementary Table 9:** DEseq2 results for HCT116 cells treated with 24 hr cobimetinib, 24 hr TNF-alpha, or untreated.

**Supplementary Table 10:** Predicted LTR10 enhancers that regulate AP1/MAPK target genes.

**Supplementary Table 11:** DEseq2 results for CRISPR silencing of LTR10.ATG12 enhancer.

**Supplementary Table 12:** DEseq2 results for CRISPR silencing of LTR10.XRCC4 enhancer.

**Supplementary Table 13:** DEseq2 results for CRISPR silencing of LTR10.MEF2D enhancer.

**Supplementary Table 14:** DEseq2 results for CRISPR silencing of LTR10.FGF2 enhancer.

**Supplementary Table 15:** DEseq2 results for CRISPR silencing of LTR10.MCPH1 enhancer.

**Supplementary Table 16:** DEseq2 results for CRISPR deletion of LTR10.KDM6A enhancer.

**Supplementary Table 17:** DEseq2 results for CRISPR silencing of ATG12 TSS.

**Supplementary Table 18:** Mouse xenograft results showing how the deletion of LTR10.XRCC4 affects tumor response to irradiation.

**Supplementary Table 19:** Fraction of H3K72ac and AP1 peaks in HCT116 cells that are derived from LTR10A/F elements.

**Supplementary Table 20:** GTEx cis-eQTLs associated with LTR10 VNTR regions.

**Supplementary Table 21:** Tumor-specific LTR10 VNTR expansions and contractions from long-read matched tumor/normal whole-genome sequencing from Xu et al (2023).

**Supplementary Table 22:** Primers, gRNAs and LTR10 sequences used for CRISPR experiments and functional validation.

#### Figure S1A

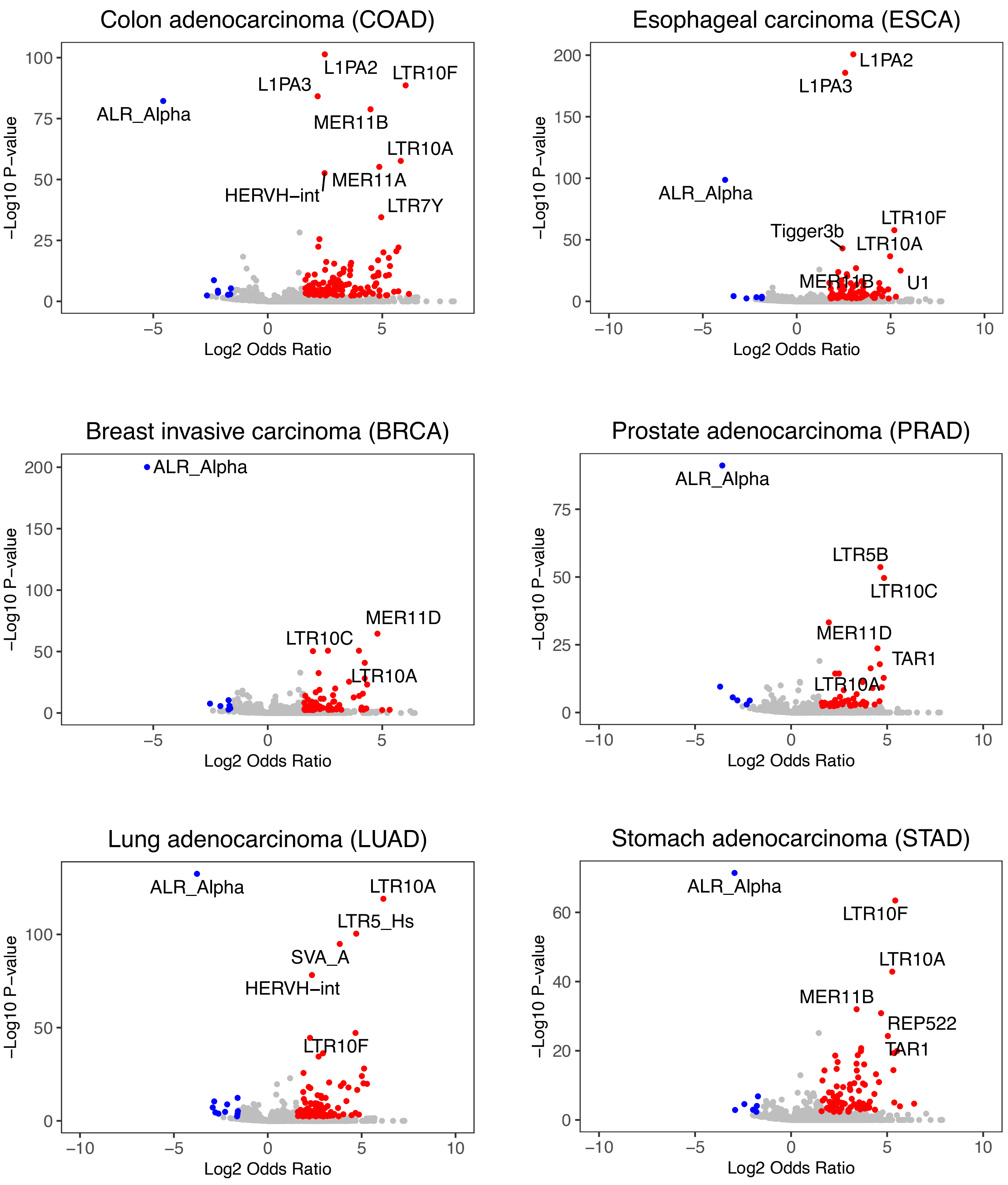


**Figure S1A:** Enrichment of TE families within cancer-specific ATAC-seq associated with different cancer subtypes from TCGA. Significantly enriched TEs are shown in red; depleted TEs are shown in blue.

#### Figure S1B

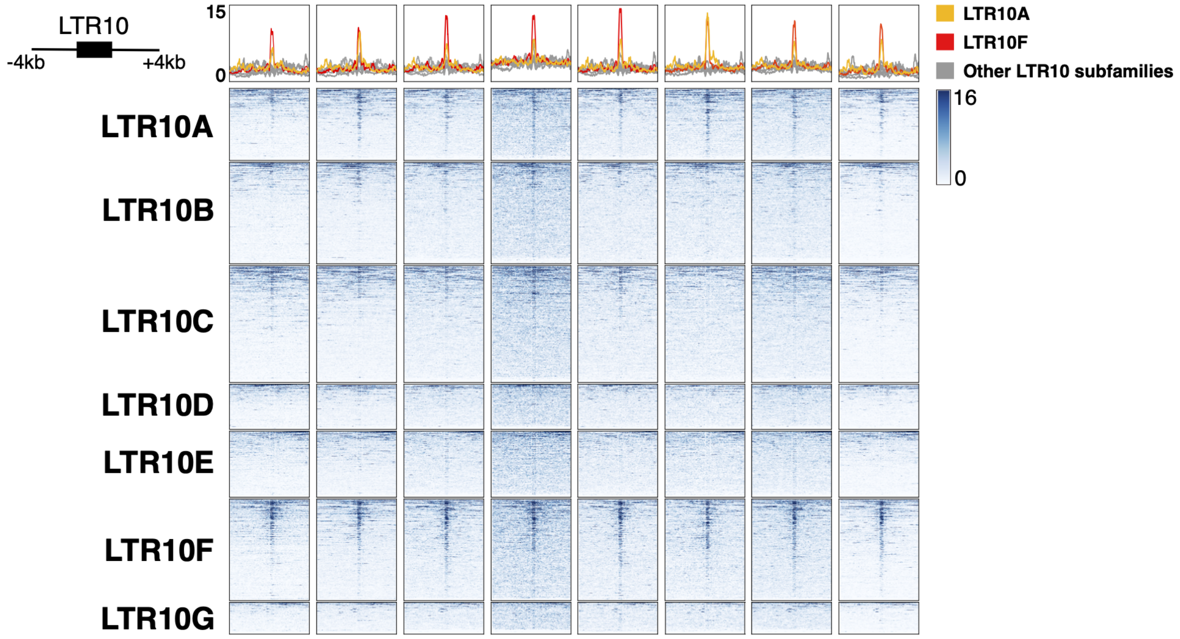


**Figure S1B:** Heatmap of representative patient tumor ATAC-seq signals (TCGA patients P053, P012, P002, P025, P004, P016, P001, P049) over all LTR10 elements, separated by subfamily. Metaprofiles represent average normalized ATAC signal across elements.

#### Figure S1C

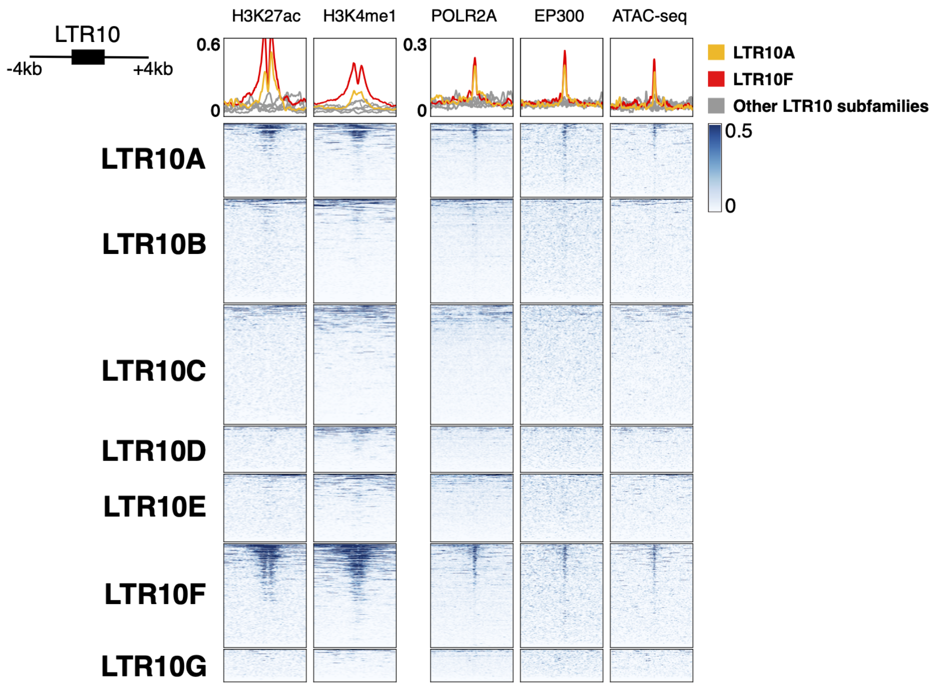


**Figure S1C:** Heatmap of enhancer-associated chromatin marks from HCT116 cells over all LTR10 elements, separated by subfamily. From left to right: H3K27ac ChIP-seq (GSE97527), H3K4me1 ChIP-seq (GSE101646), POLR2A ChIP-seq (GSE32465), EP300 ChIP-seq (GSE51176), and HCT116 ATAC-seq (GSE126215). Metaprofiles represent the normalized signal across elements.

### Figure S1D

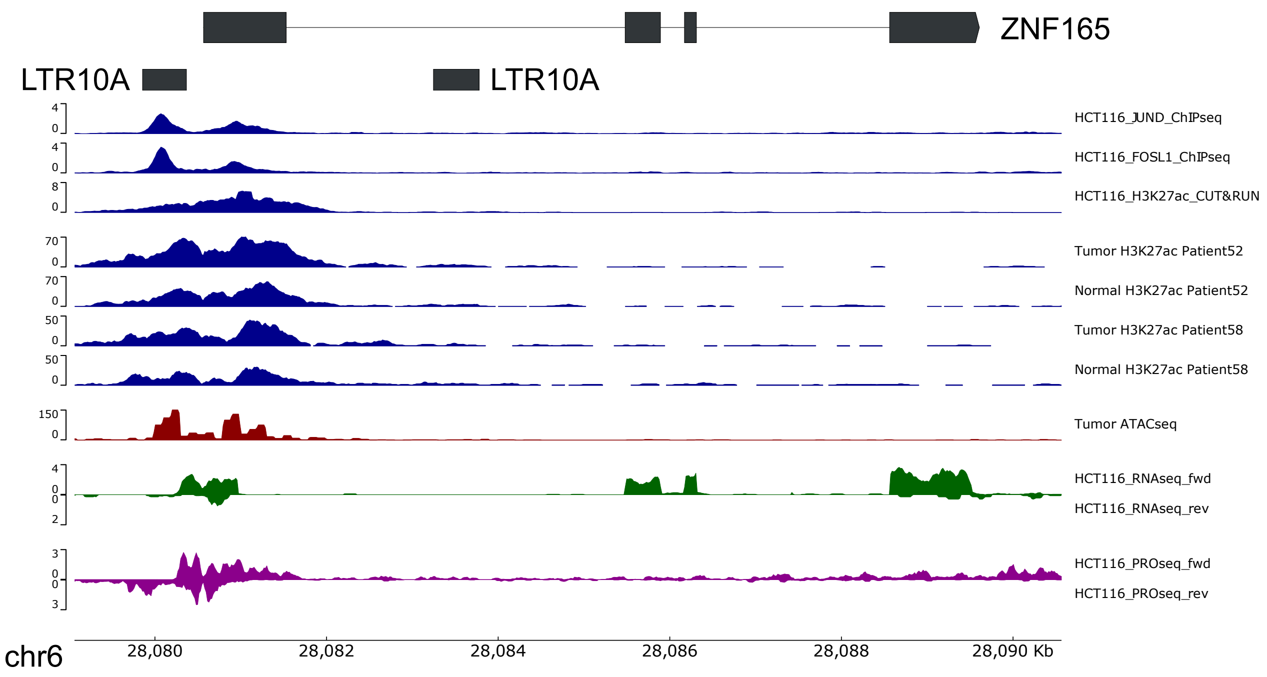


**Figure S1D:** Genome browser view of an LTR10A element co-opted as a promoter for ZNF165. From top to bottom: JUND and FOSL1 ChIP-seq (GSE32465), H3K27ac CUT&RUN (in-house), H3K27ac ChIP-seq from matched tumor/normal samples from the CEMT Canadian Epigenome Project (patients AKCC52 and AKCC58), tumor ATAC-seq from TCGA (patient COAD P022), HCT116 RNA-seq (in-house), and HCT116 PRO-seq (GSE129501).

#### Figure S2A

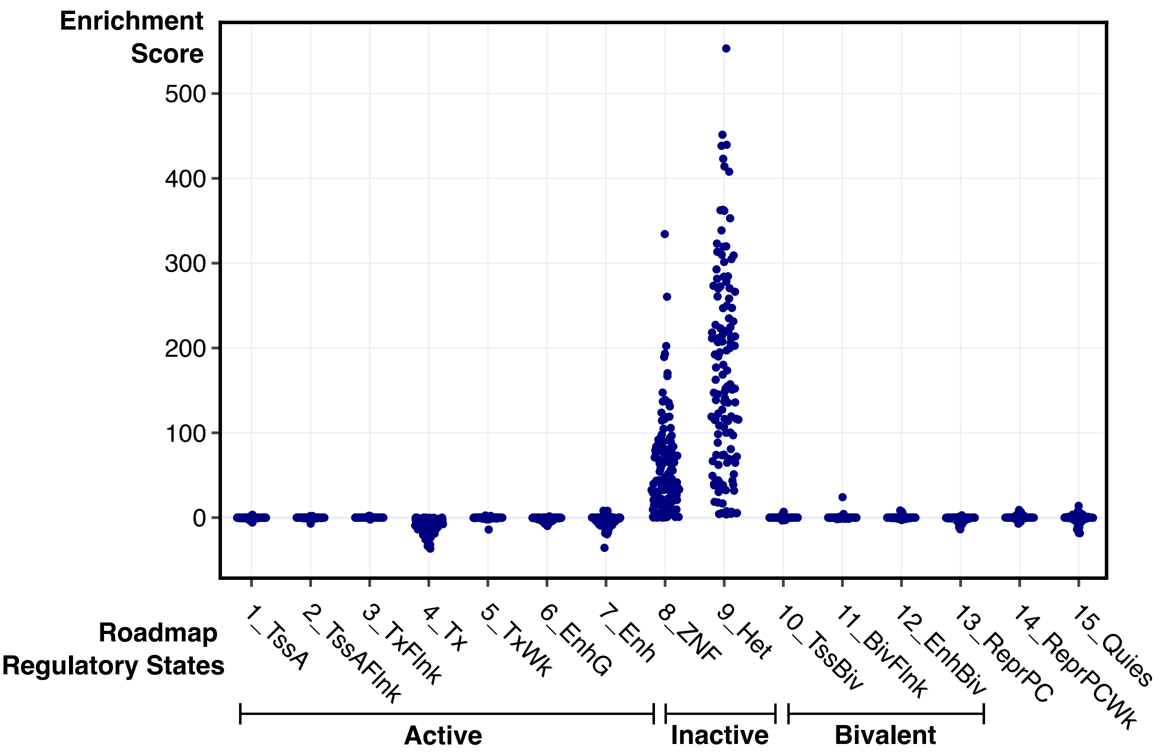


**Figure S2A:** Dotplot showing enrichment scores of LTR10A/F elements for all Roadmap tissues (N=127), across the fifteen regulatory states defined by Roadmap.

#### Figure S2B

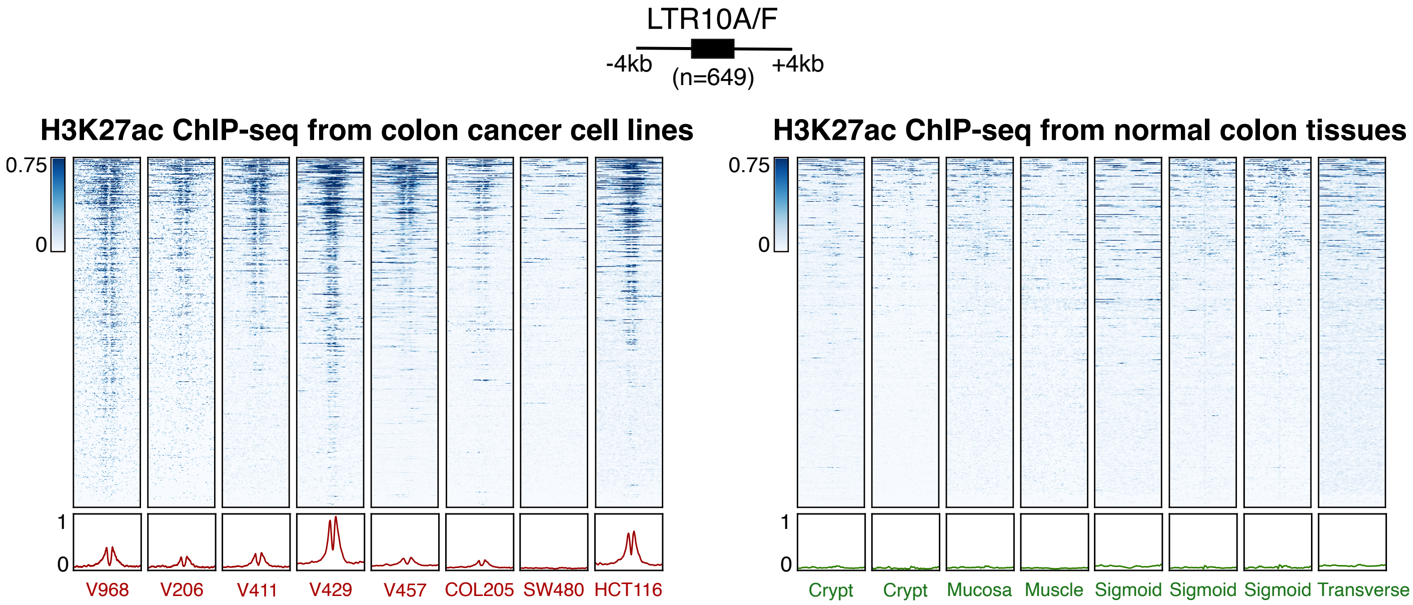


**Figure S2B:** Heatmap of H3K27ac ChIP-seq from different colorectal cancer cell lines (GSE77737) versus H3K27ac ChIP-seq from normal colon tissues (GSE77737, GSE17312, GSE101136, GSE101031, GSE16256), over the merged set of 649 LTR10A/F elements. Bottom metaprofiles represent the normalized signal across elements.

#### Figure S2C

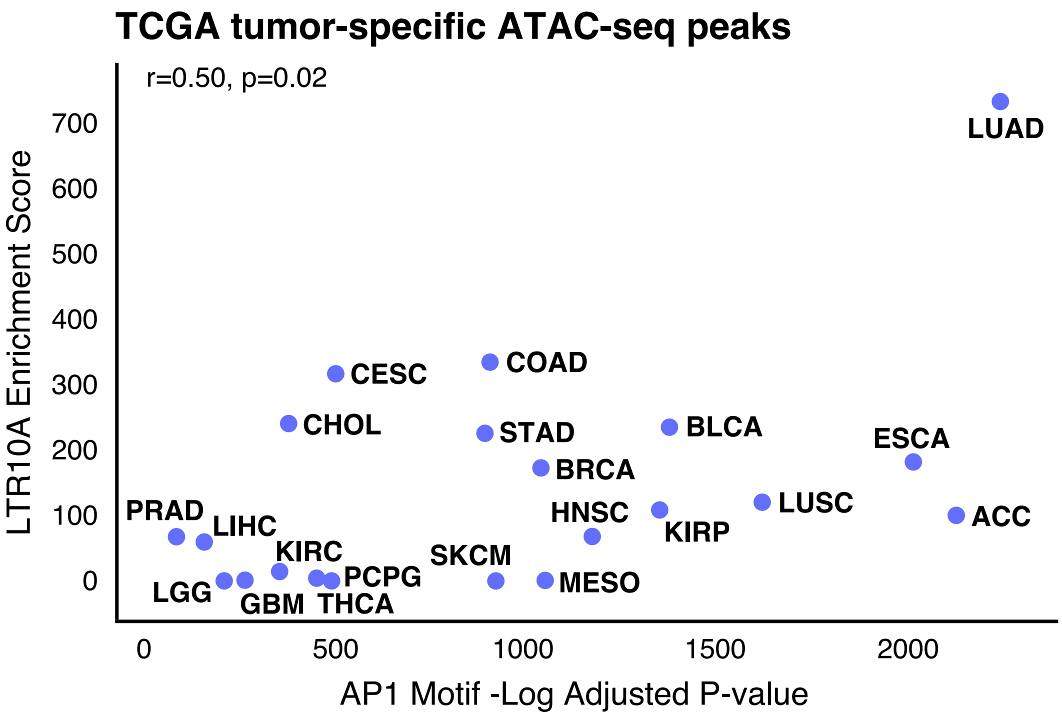


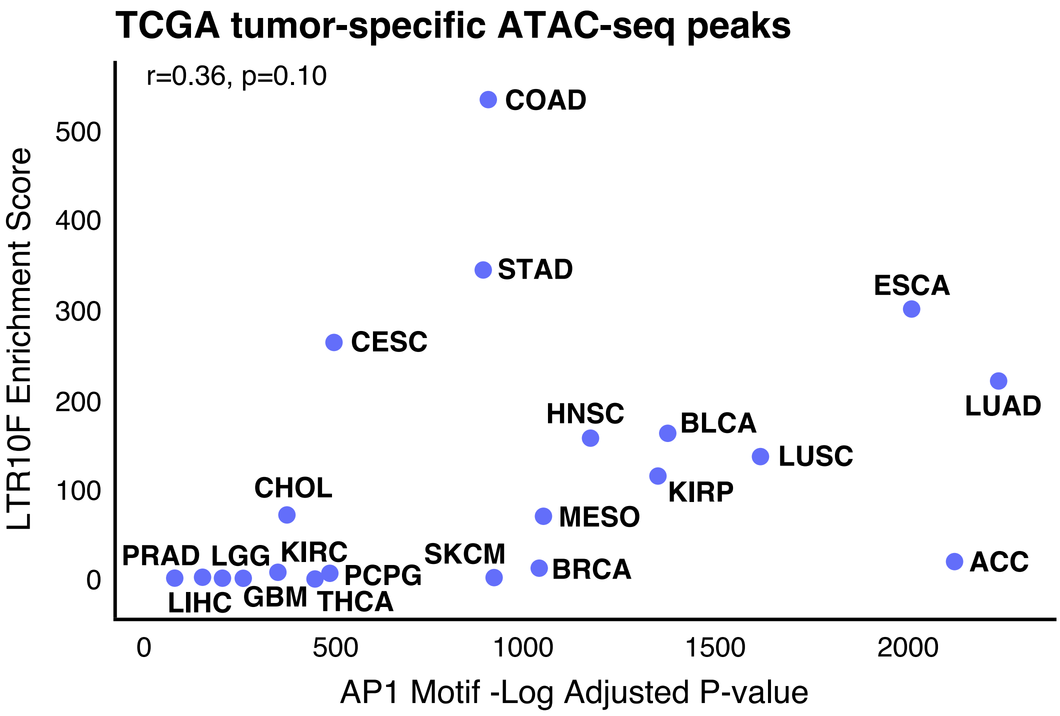


**Figure S2C:** Enrichment of LTR10A or LTR10F versus enrichment of AP1 binding motifs in tumor-specific ATAC-seq peaks for 21 cancer types. Cancer types are labeled by their TCGA abbreviations (e.g. COAD = colon adenocarcinoma). The Pearson correlation coefficient (r) and p-value are shown for each plot.

#### Figure S2D

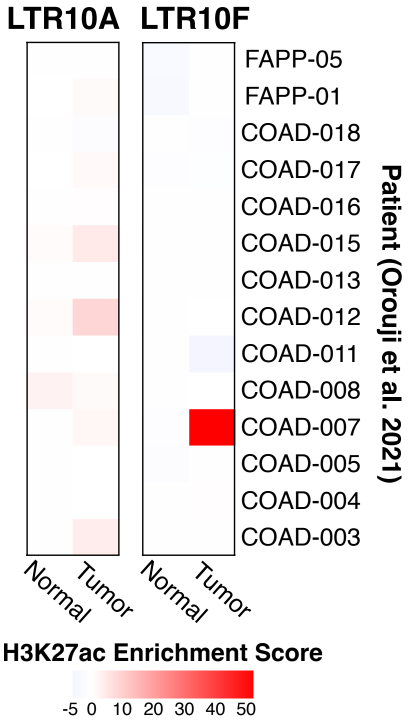


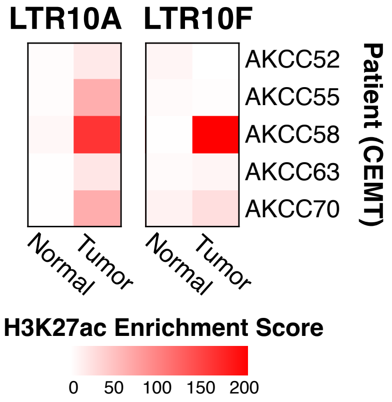


**Figure S2D:** Enrichment of LTR10A/F elements in H3K27ac-defined regulatory regions from patient matched tumor/normal samples. ChIP-Seq data obtained from Orouji et al (2021), and the CEMT Canadian Epigenome Project.

#### Figure S2E

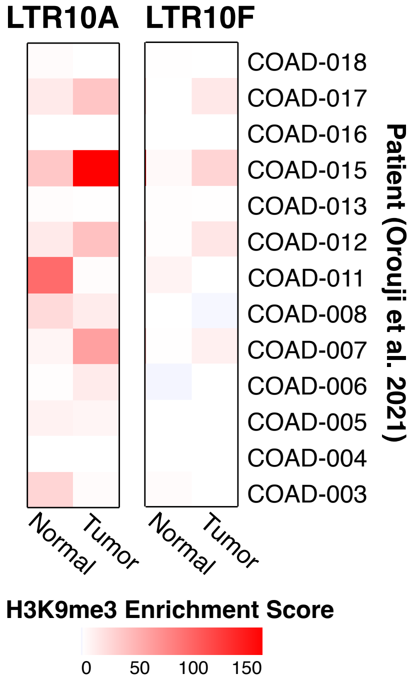


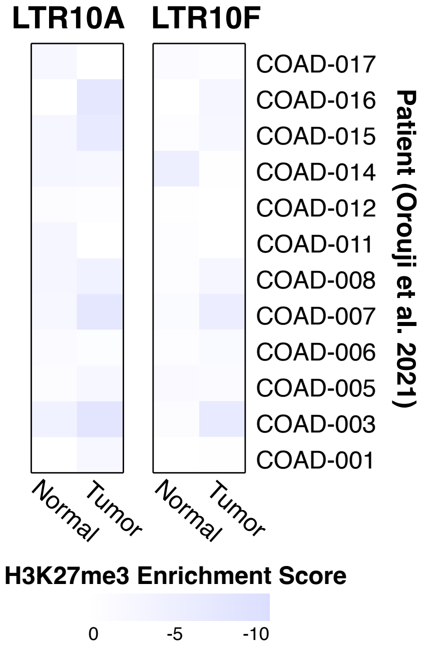


**Figure S2E:** Enrichment of LTR10A/F elements in H3K9me3 and H3K27me3 regulatory regions from patient matched tumor/normal samples. ChIP-Seq data obtained from Orouji et al (2021).

#### Figure S2F

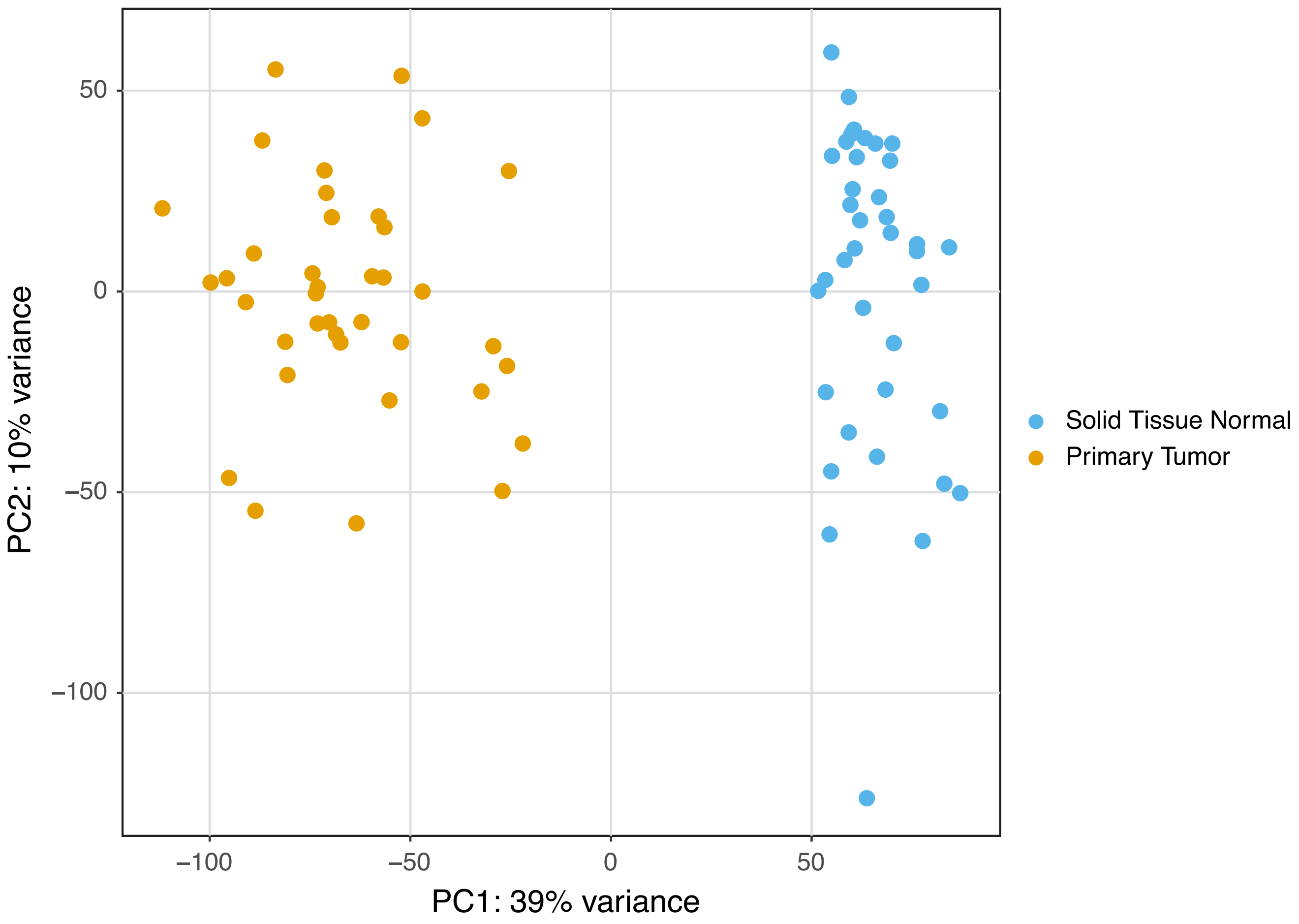


**Figure S2F:** Principal component analysis (PCA) of bulk RNA-seq data from TCGA-COAD controlled access data, showing matched tumor/normal samples from 38 patients with colorectal adenocarcinomas. Each patient has one tumor and one normal colon sample. PCA is based on gene expression only (TEs not included).

#### Figure S2G

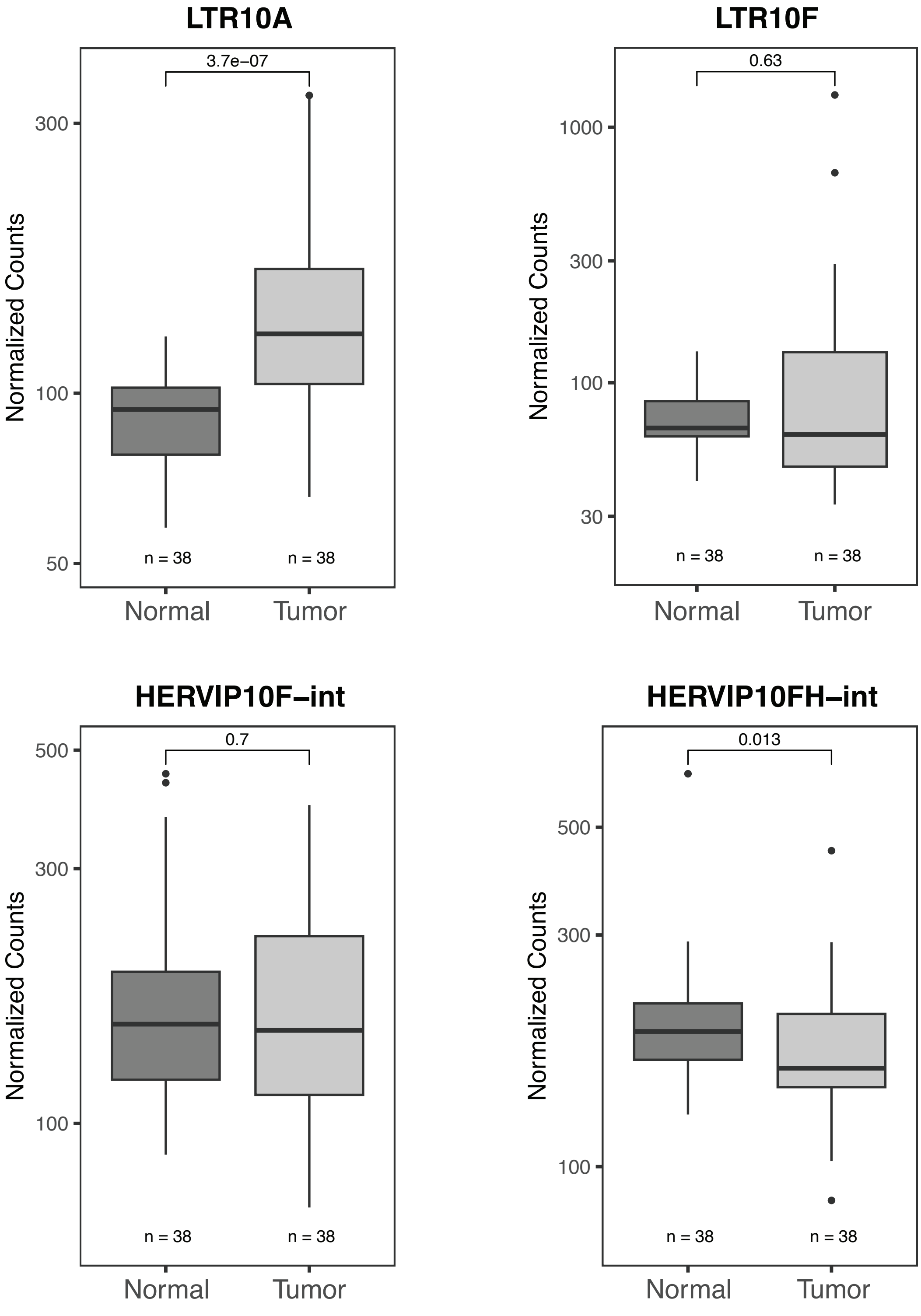


**Figure S2G:** Boxplots of paired sample Wilcoxon test p-values for normalized counts of LTR10 transcripts from bulk RNA-seq of 38 patient-derived matched tumor/normal samples. Normalized counts are depicted on a log scale. RNA-seq was downloaded from TCGA-COAD controlled access data.

#### Figure S2H

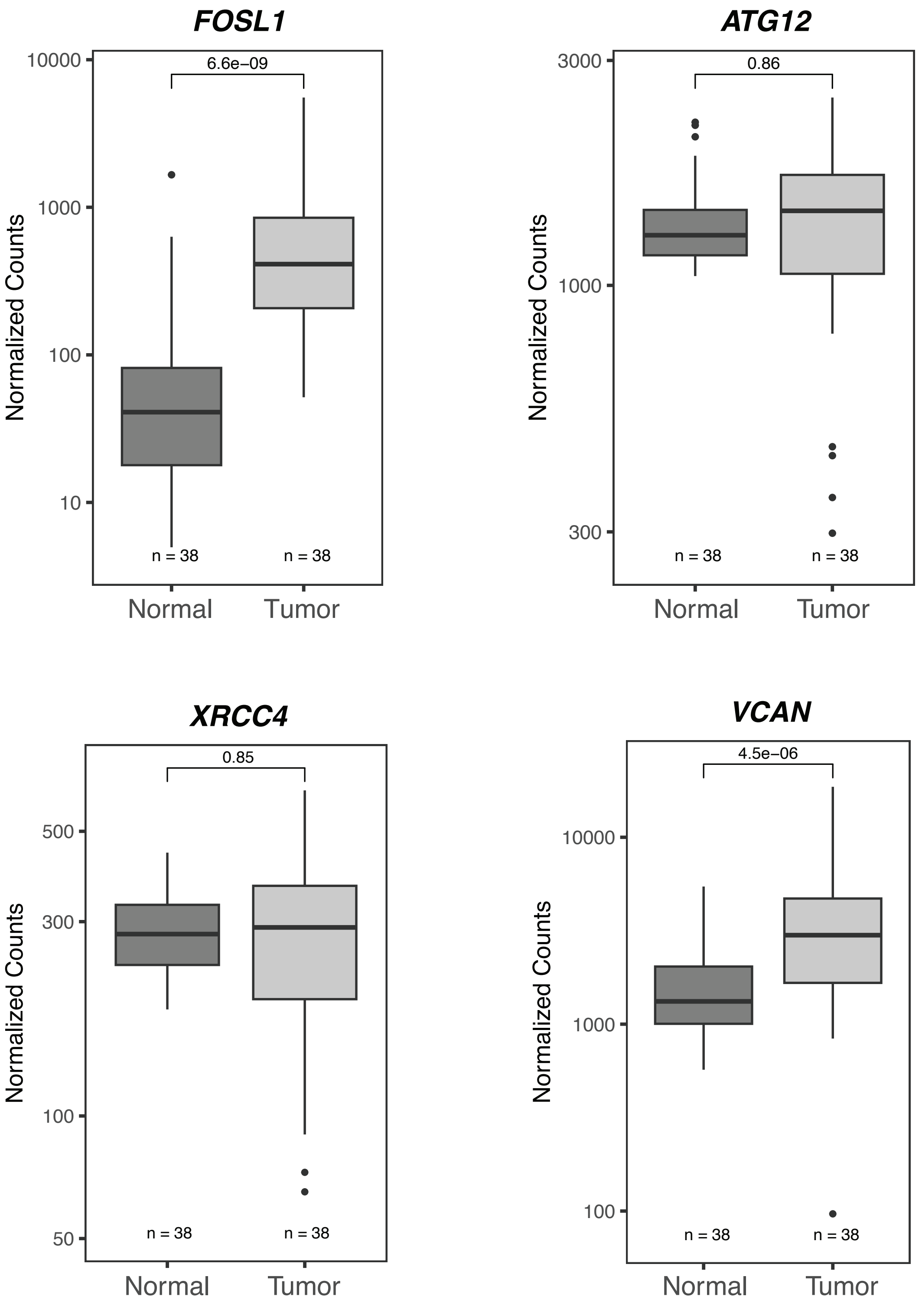


**Figure S2H:** Boxplots of paired sample Wilcoxon test p-values for normalized counts of LTR10-associated genes from bulk RNA-seq of 38 patient-derived matched tumor/normal samples. Normalized counts are depicted on a log scale. RNA-seq was downloaded from TCGA-COAD controlled access data.

### Figure S2I

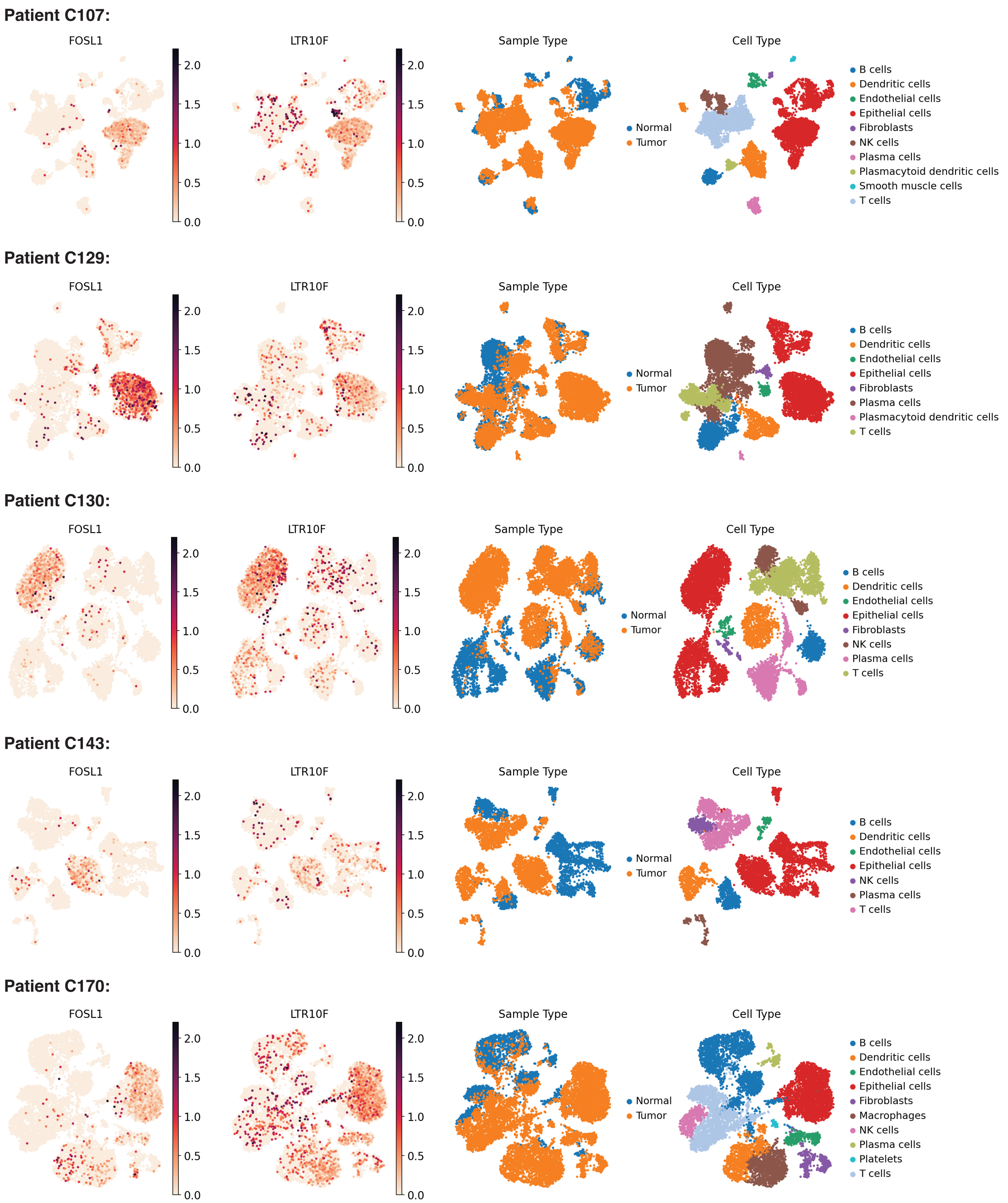


**Figure S2I:** UMAP visualization of single cell RNA-seq from Pelka et al (2021), showing five patients with significant LTR10/FOSL1 expression in tumor epithelial cells.

### Figure S2J

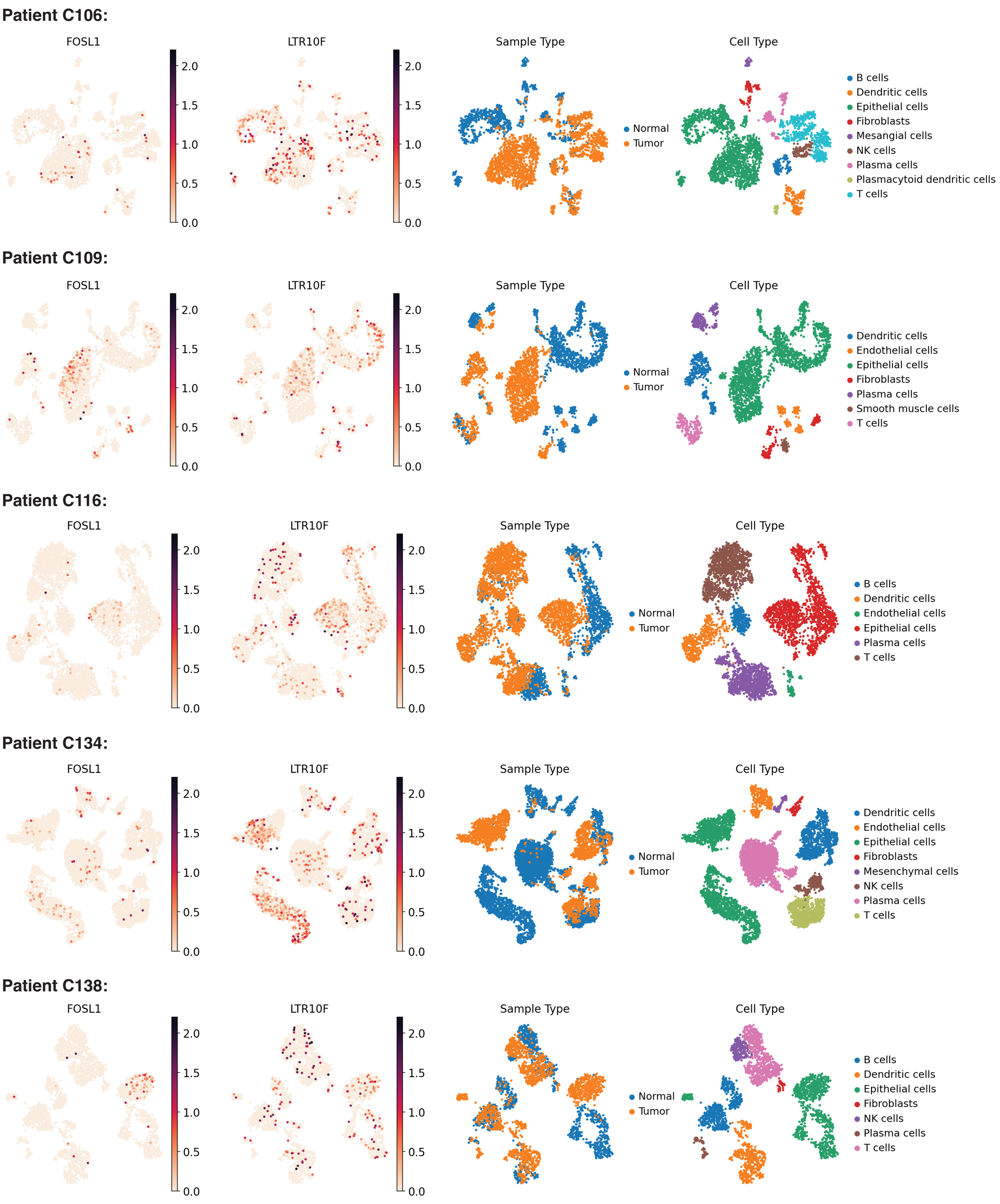


**Figure S2J:** UMAP visualization of single cell RNA-seq from Pelka et al (2021), showing five patients with no significant LTR10/FOSL1 expression in tumor cells.

#### Figure S2K

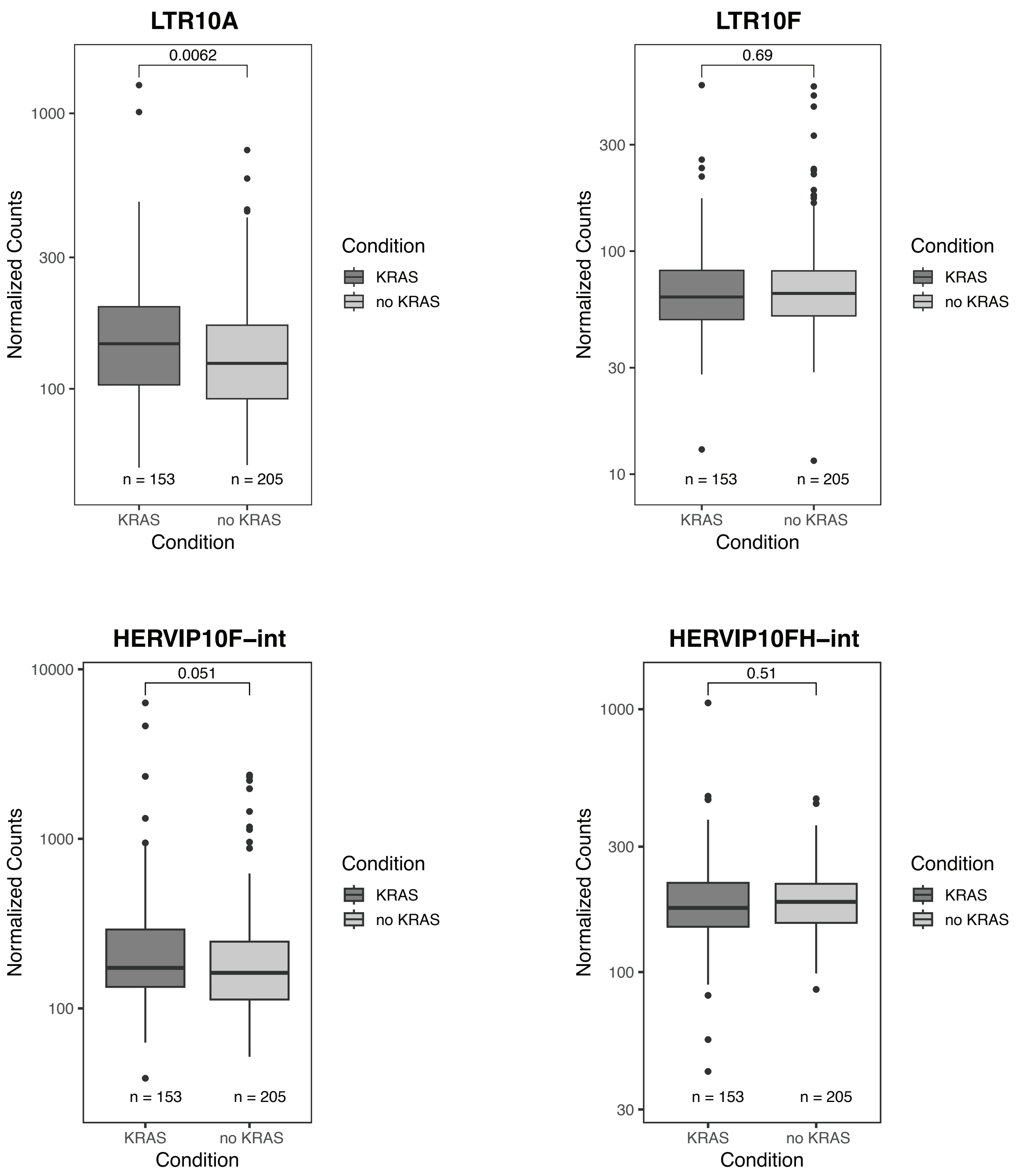


**Figure S2K:** Boxplots of Wilcoxon test p-values (unpaired) for normalized counts of LTR10 transcripts from bulk RNA-seq of 358 colorectal tumors with KRAS mutation status. KRAS = tumor with KRAS mutation; no KRAS = tumor without KRAS mutation. Normalized counts are depicted on a log scale. Tumor RNA-seq was downloaded from TCGA-COAD controlled access data.

**Figure S2L**

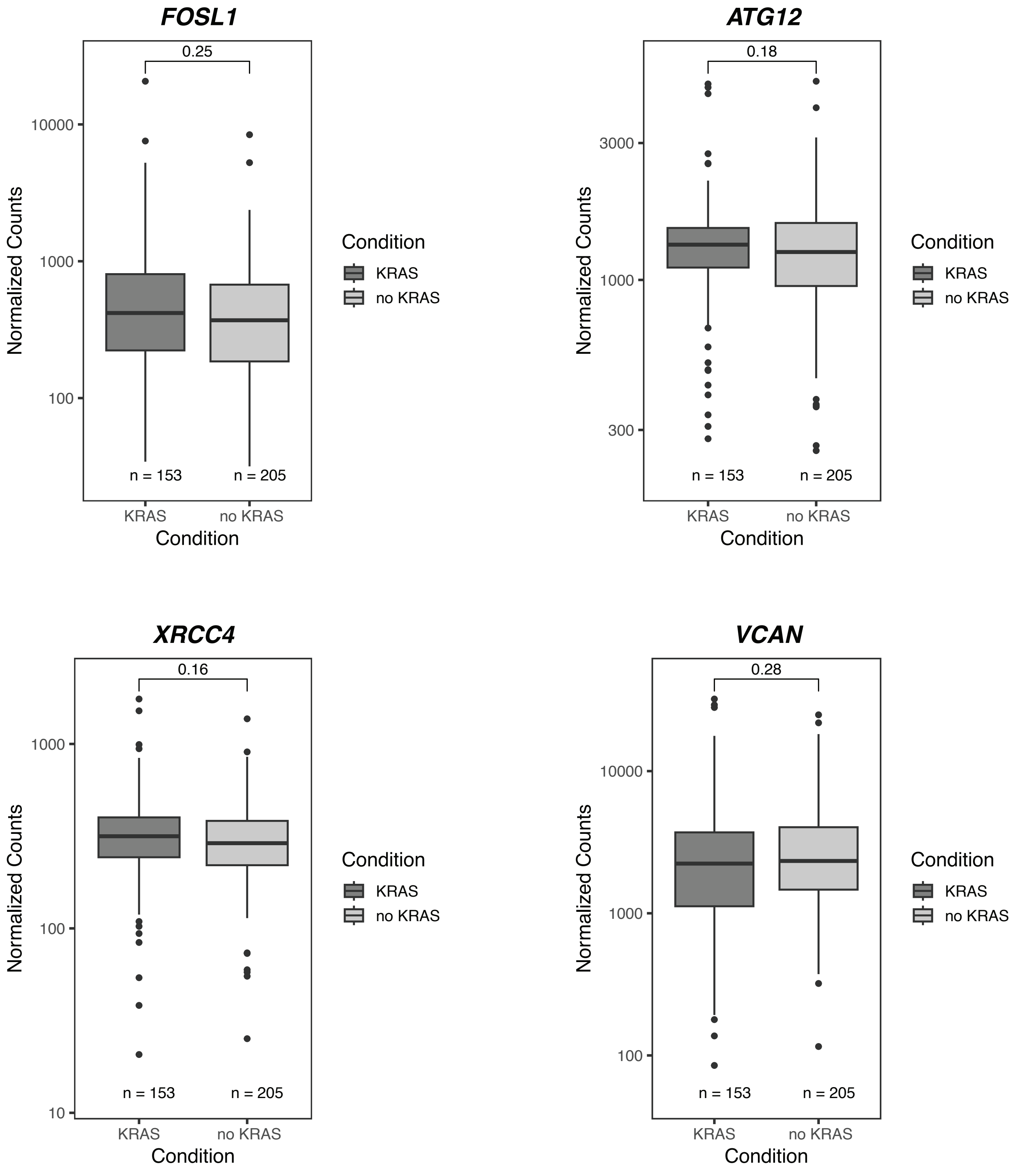


**Figure S2L:** Boxplots of Wilcoxon test p-values (unpaired) for normalized counts of LTR10-associated genes from bulk RNA-seq of 358 colorectal tumors with KRAS mutation status. KRAS = tumor with KRAS mutation; no KRAS = tumor without KRAS mutation. Normalized counts are depicted on a log scale. Tumor RNA-seq was downloaded from TCGA-COAD controlled access data.

#### Figure S2M

LTR10A

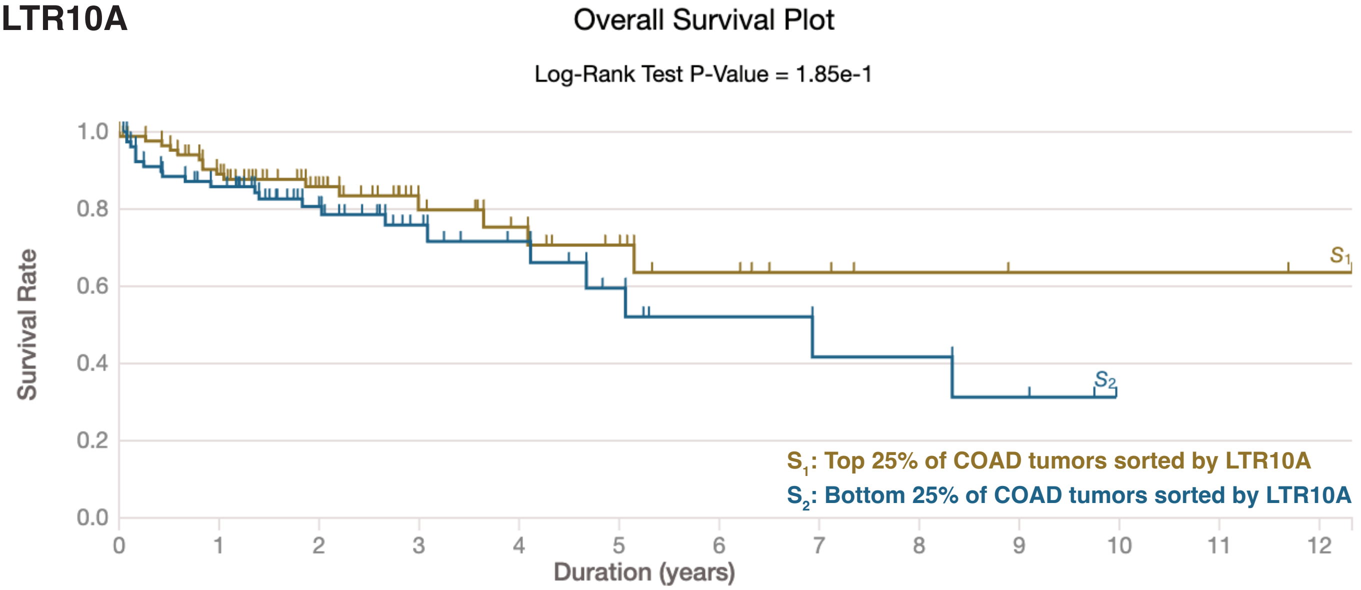


LTR10F

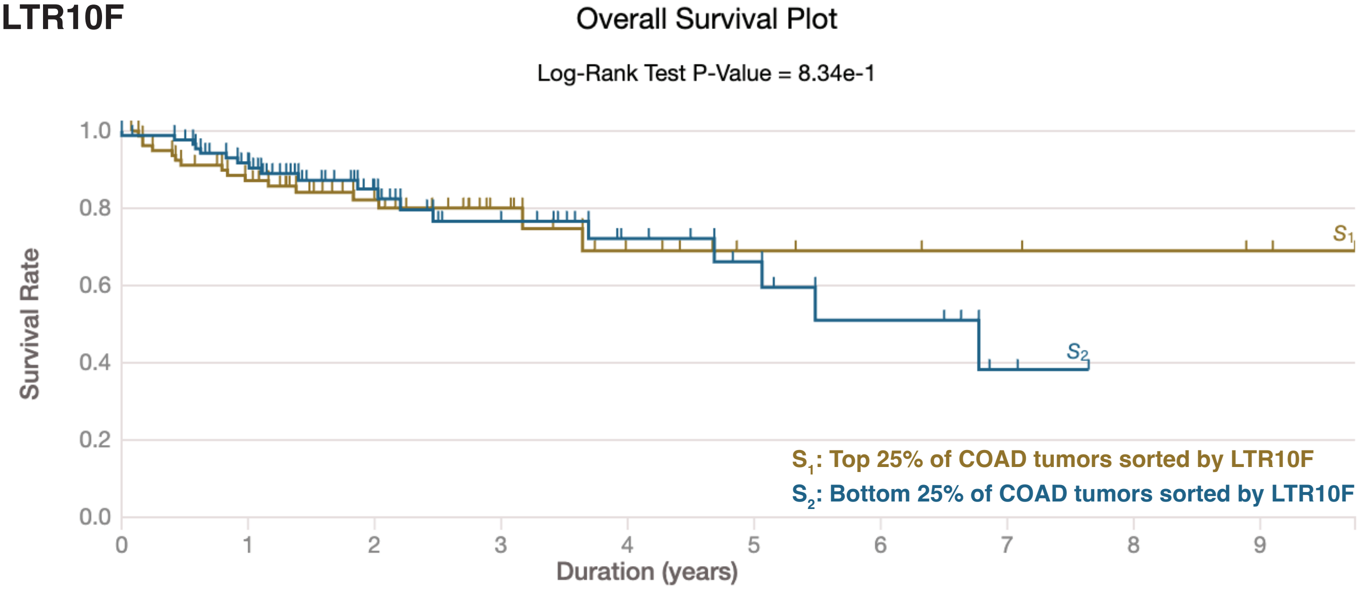


**Figure S2M:** Kaplan Meier survival analyses based on expression of LTR10 transcripts. 358 colorectal tumors from TCGA-COAD were sorted based on expression of the LTR10 element of interest. For each LTR10 element, the upper quartile (i.e. top 90 tumors) was compared against the lower quartile (i.e. bottom 90 tumors) with respect to survival rate.

#### Figure S2N

*FOSL1*

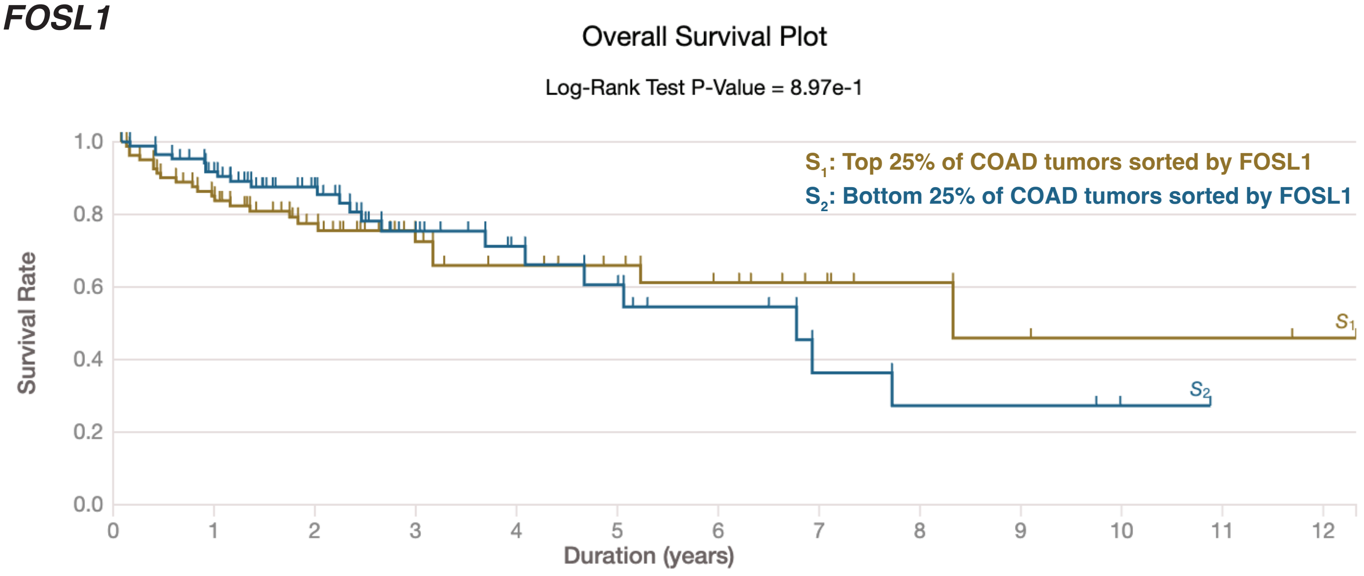


*ATG12*

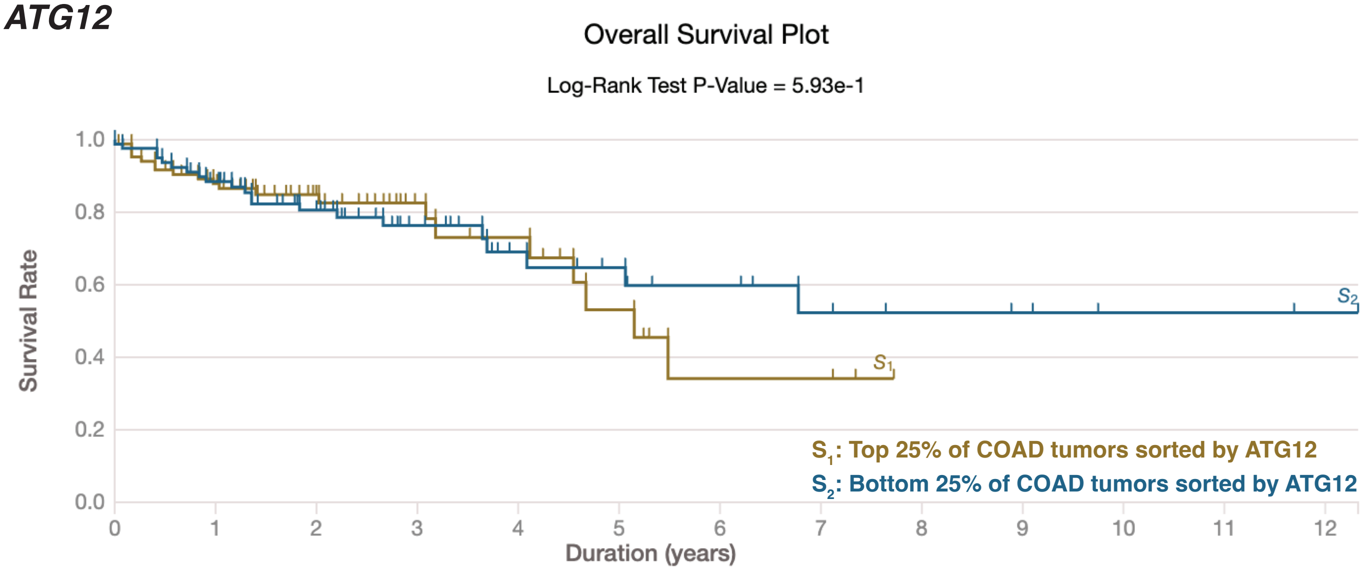


**Figure S2N:** Kaplan Meier survival analyses based on expression of LTR10-associated genes. 358 colorectal tumors from TCGA-COAD were sorted based on expression of the gene of interest. For each gene, the upper quartile (i.e. top 90 tumors) was compared against the lower quartile (i.e. bottom 90 tumors) with respect to survival rate.

#### Figure S3A

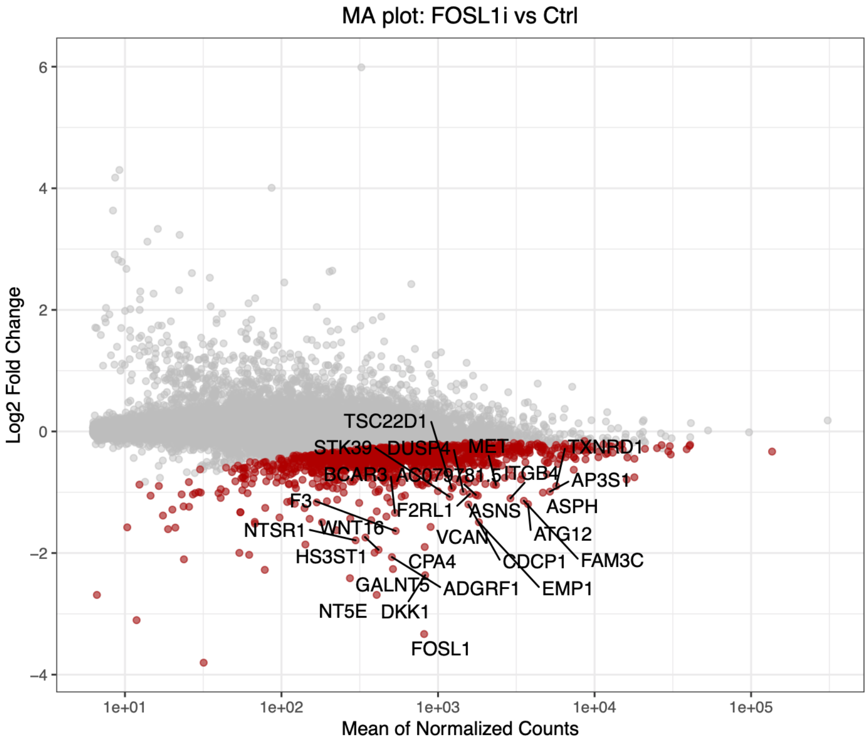


**Figure S3A:** MA plot showing global gene expression changes in cells in response to silencing FOSL1. Significantly downregulated genes are shown in red.

#### Figure S3B

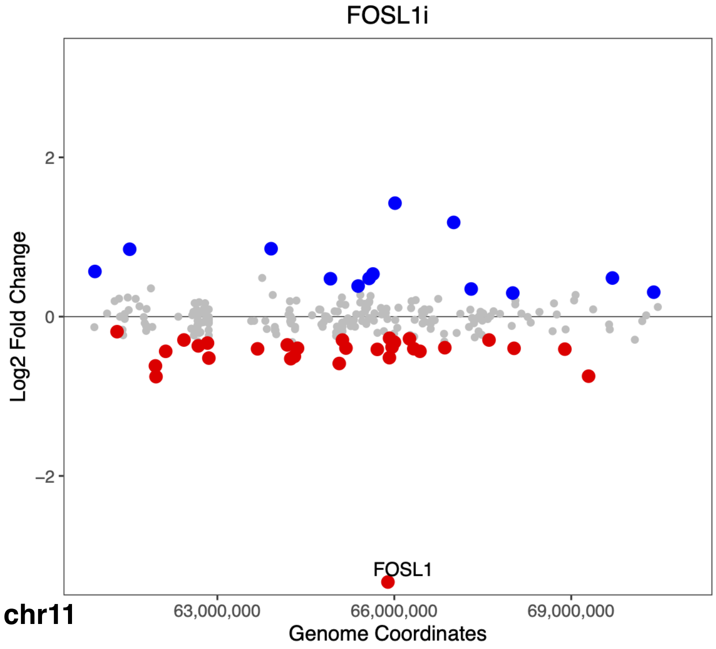


**Figure S3B:** Scatterplot of gene expression changes in response to silencing FOSL1 in a 10Mb window around the target. Significantly down-regulated genes are shown in red, significantly up-regulated genes are shown in blue. The most significantly downregulated gene (FOSL1) is labelled.

#### Figure S3C

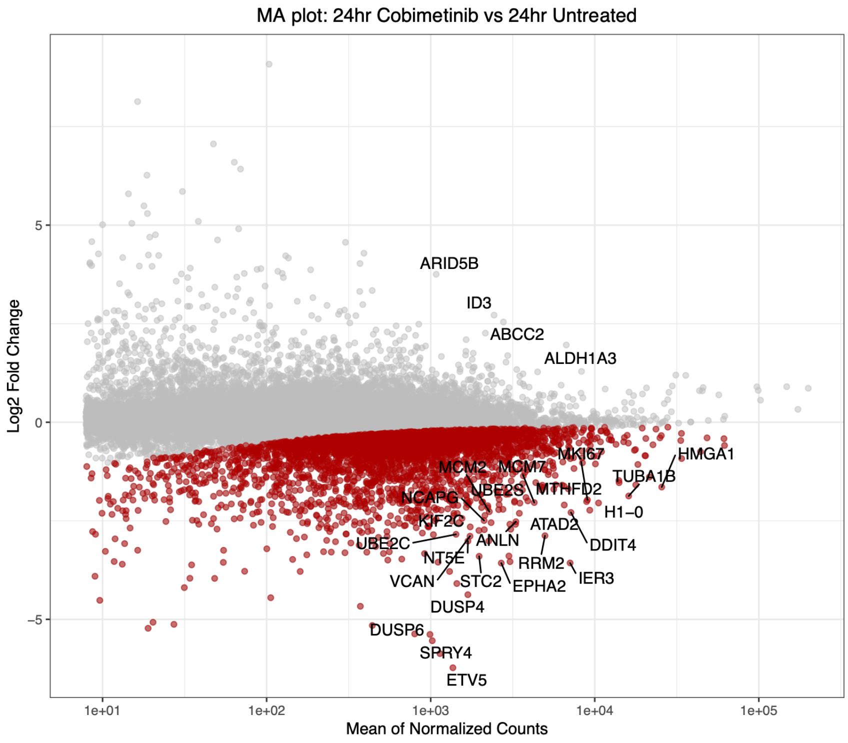


**Figure S3C:** MA plot showing global gene expression changes in cells in response to 24hr cobimetinib treatment. Significantly downregulated genes are shown in red.

#### Figure S3D

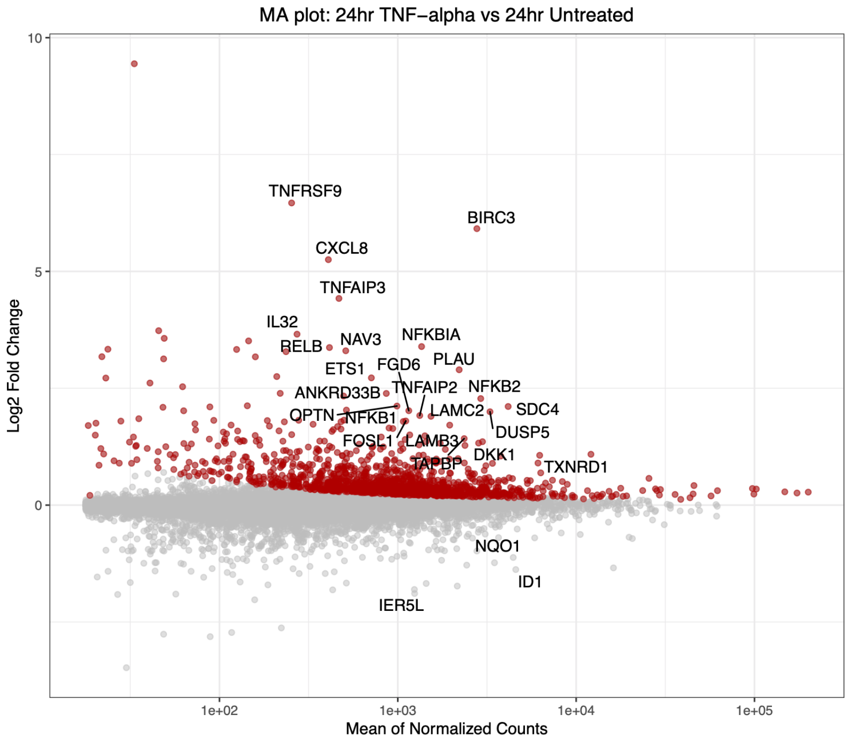


**Figure S3D:** MA plot showing global gene expression changes in cells in response to 24hr TNF-alpha treatment. Significantly upregulated genes are shown in red.

#### Figure S3E

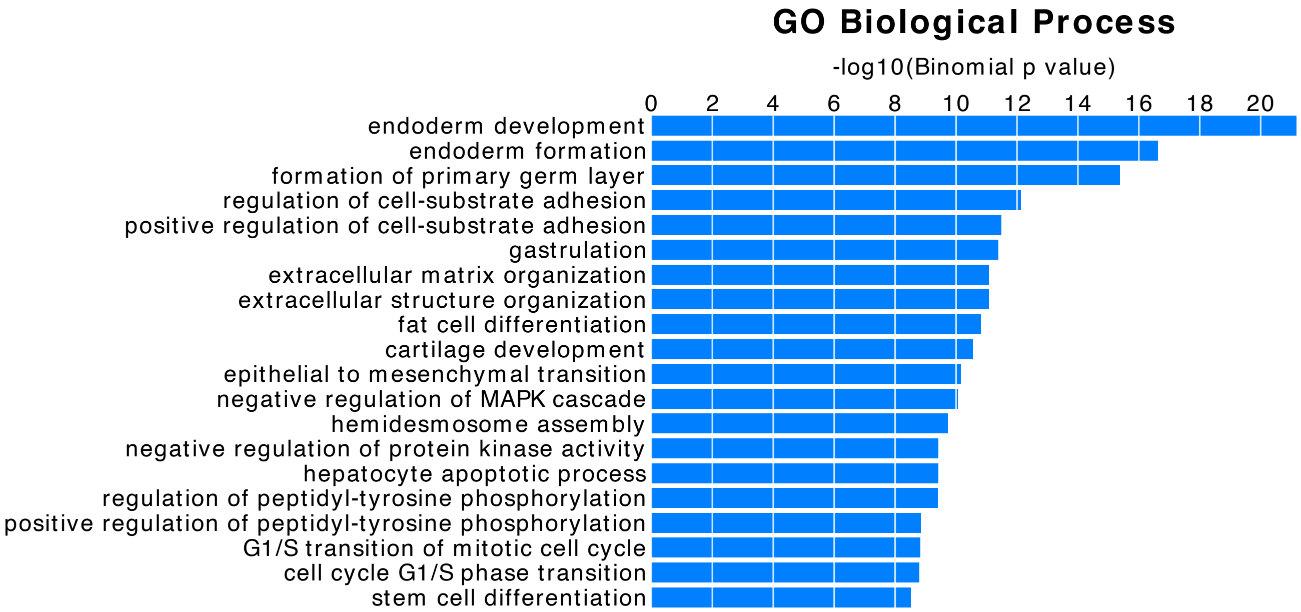


**Figure S3E:** Bargraph of Gene Ontology (GO) biological processes associated with the set of H3K27ac regions significantly downregulated by cobimetinib (N=1634), as predicted by GREAT.

#### Figure S3F

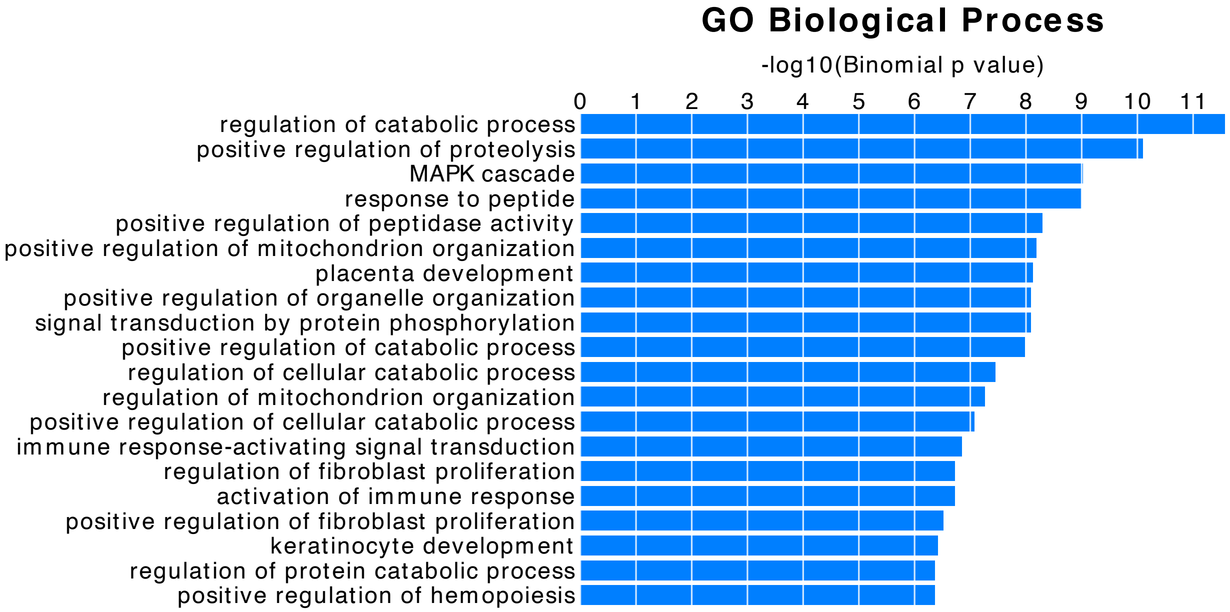


**Figure S3F:** Bargraph of Gene Ontology (GO) biological processes associated with the set of H3K27ac regions significantly upregulated by TNF-alpha (N=775), as predicted by GREAT.

### Figure S3G

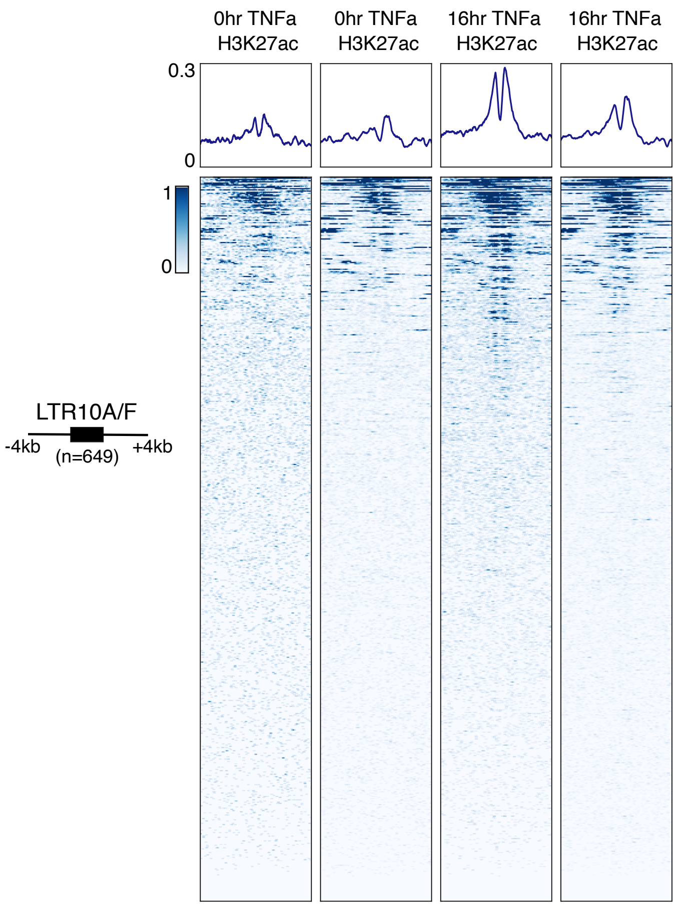


**Figure S3G:** Heatmap of H3K37ac ChIP-seq from SW480 colon cancer cells, either untreated or treated with TNF-alpha for 16hr, over the merged set of 649 LTR10A/F elements (GSE102796). Metaprofiles represent the normalized signal across elements.

#### Figure S3H

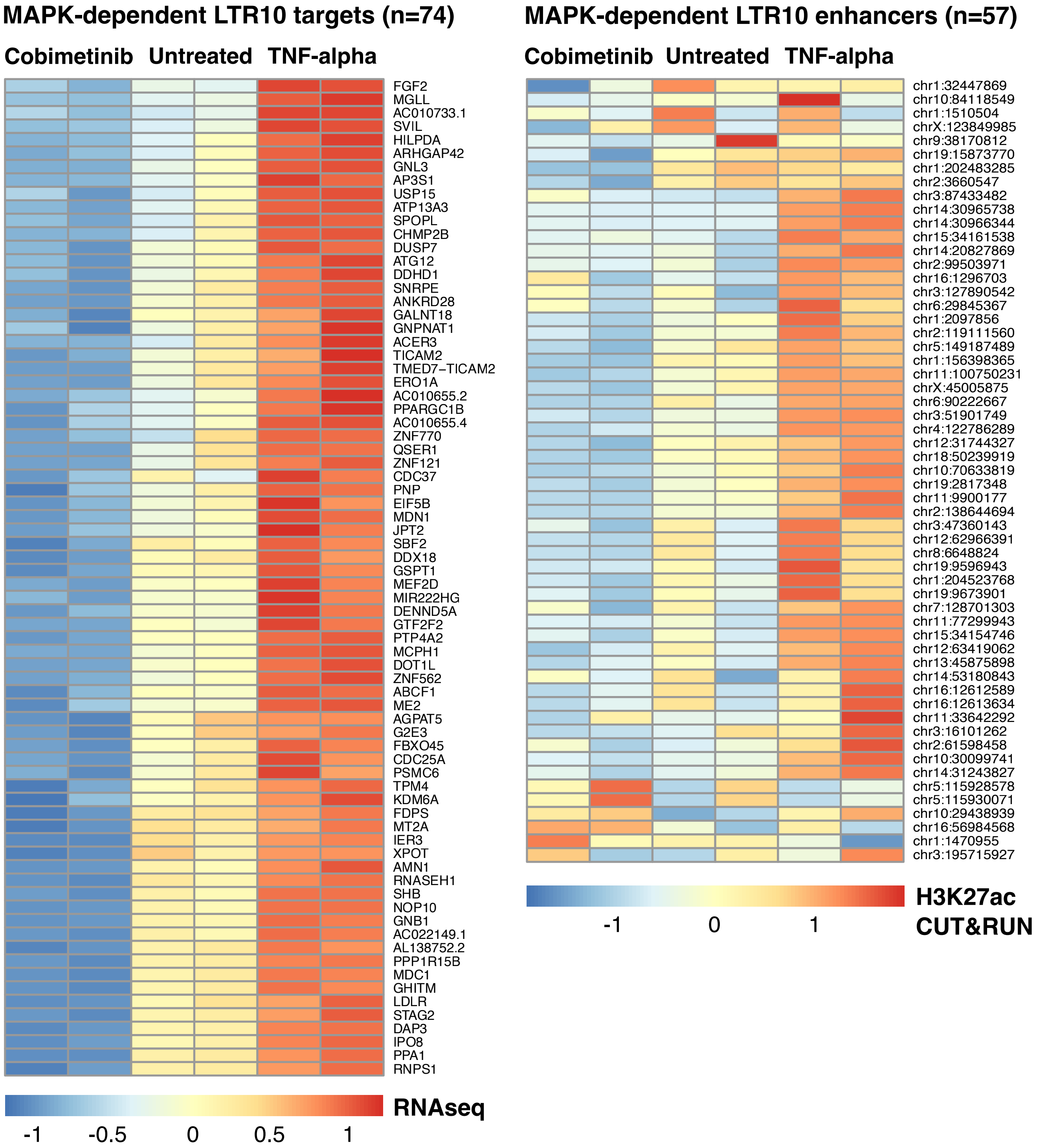


**Figure S3H:** Heatmaps of normalized RNA-seq expression values (left) and normalized H3K27ac CUT&RUN signal (right) for MAPK-dependent predicted LTR10 target genes and enhancers. Normalized values for each treatment replicate are shown.

#### Figure S4A

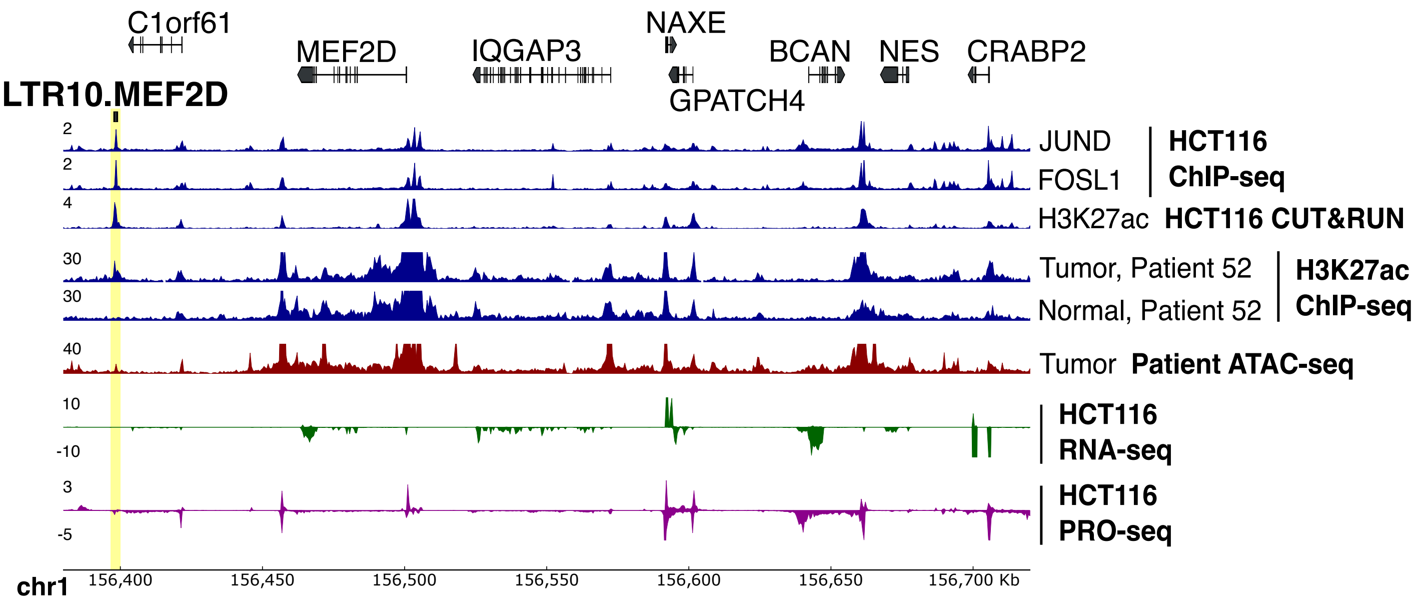


**Figure S4A:** Genome browser screenshot of the *MEF2D* locus with the LTR10.MEF2D enhancer labeled. From top to bottom: JUND and FOSL1 ChIP-seq (GSE32465), H3K27ac CUT&RUN (in-house), H3K27ac ChIP-seq from matched tumor/normal samples from the CEMT Canadian Epigenome Project (patient AKCC52), tumor ATAC-seq from TCGA (patient COAD P022), HCT116 RNA-seq (in-house), and HCT116 PRO-seq (GSE129501). Axis numbers represent the upper limit of the range; the lower limit is always zero.

#### Figure S4B

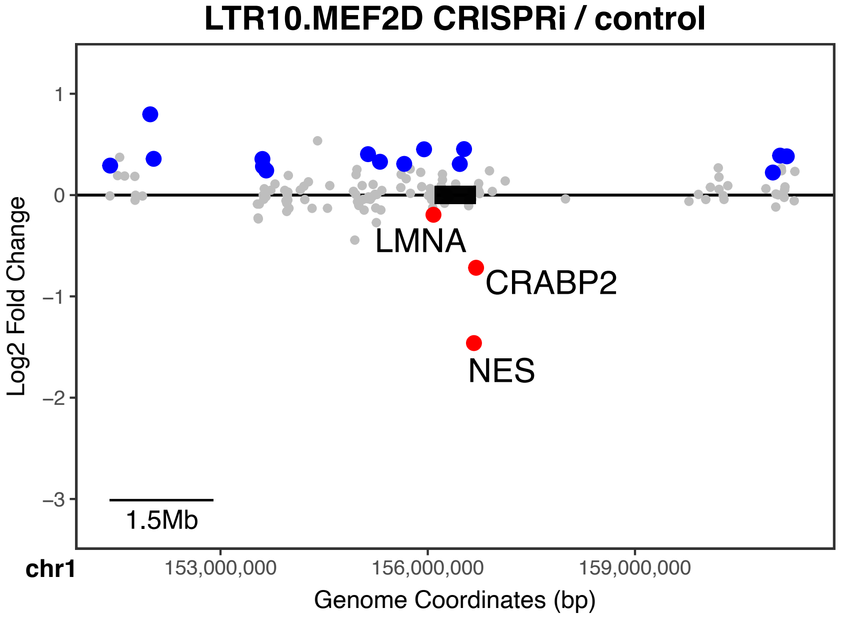


**Figure S4B:** Scatterplot of local gene expression changes in response to CRISPR silencing of the LTR10.MEF2D element. Significantly downregulated genes are shown in red; significantly upregulated genes are shown in blue. Significantly downregulated genes within 1.5 MB of the targeted element are labeled (element box not drawn to scale).

#### Figure S4C

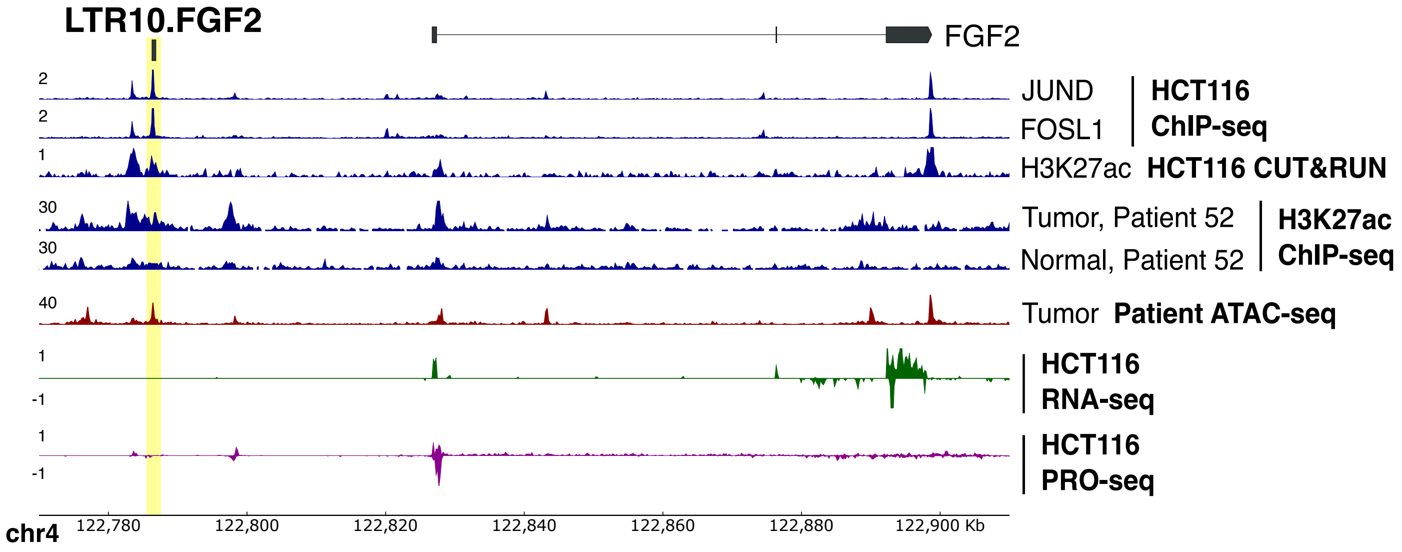


**Figure S4C:** Genome browser screenshot of the *FGF2* locus with the LTR10.FGF2 enhancer labeled. From top to bottom: JUND and FOSL1 ChIP-seq (GSE32465), H3K27ac CUT&RUN (in-house), H3K27ac ChIP-seq from matched tumor/normal samples from the CEMT Canadian Epigenome Project (patient AKCC52), tumor ATAC-seq from TCGA (patient COAD P022), HCT116 RNA-seq (in-house), and HCT116 PRO-seq (GSE129501). Axis numbers represent the upper limit of the range; the lower limit is always zero.

#### Figure S4D

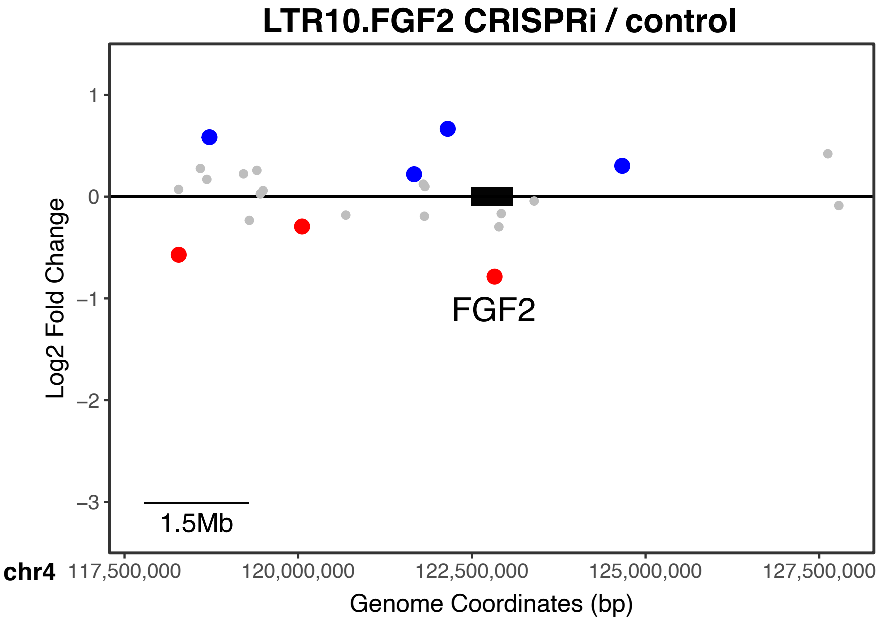


**Figure S4D:** Scatterplot of local gene expression changes in response to CRISPR silencing of the LTR10.FGF2 element. Significantly downregulated genes are shown in red; significantly upregulated genes are shown in blue. Significantly downregulated genes within 1.5 MB of the targeted element are labeled (element box not drawn to scale).

#### Figure S4E

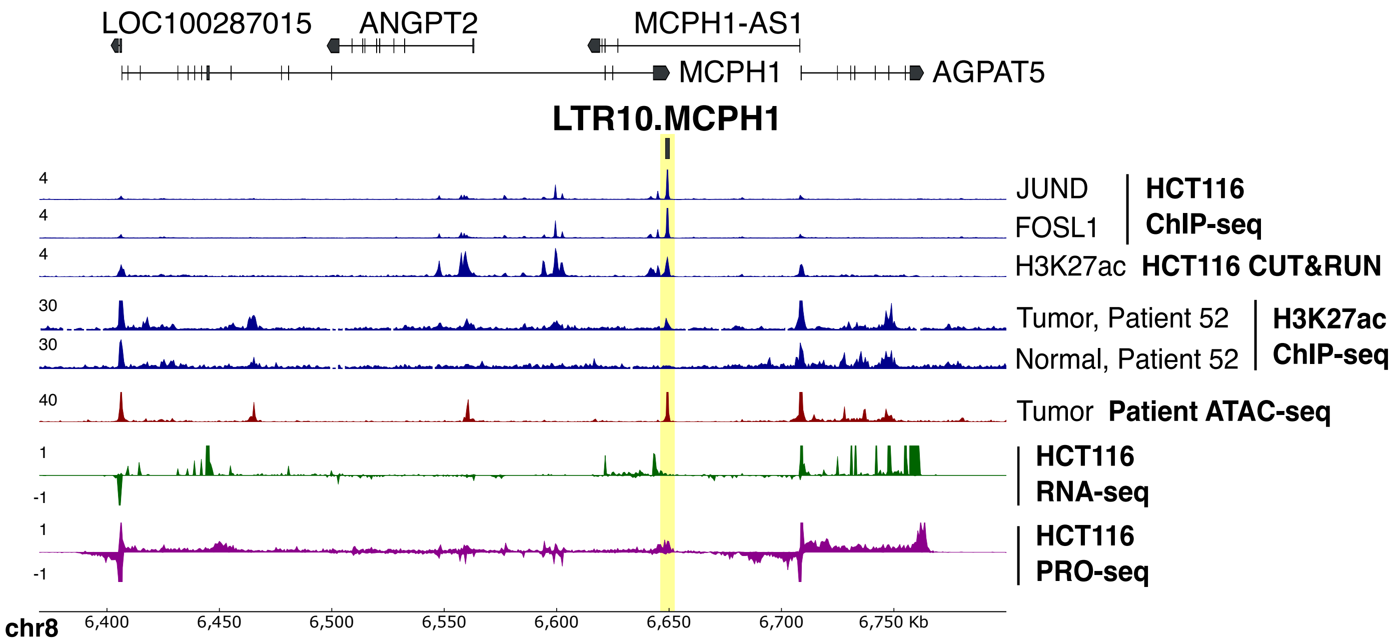


**Figure S4E:** Genome browser screenshot of the *MCPH1* locus with the LTR10.MCPH1 enhancer labeled. From top to bottom: JUND and FOSL1 ChIP-seq (GSE32465), H3K27ac CUT&RUN (in-house), H3K27ac ChIP-seq from matched tumor/normal samples from the CEMT Canadian Epigenome Project (patient AKCC52), tumor ATAC-seq from TCGA (patient COAD P022), HCT116 RNA-seq (in-house), and HCT116 PRO-seq (GSE129501). Axis numbers represent the upper limit of the range; the lower limit is always zero.

#### Figure S4F

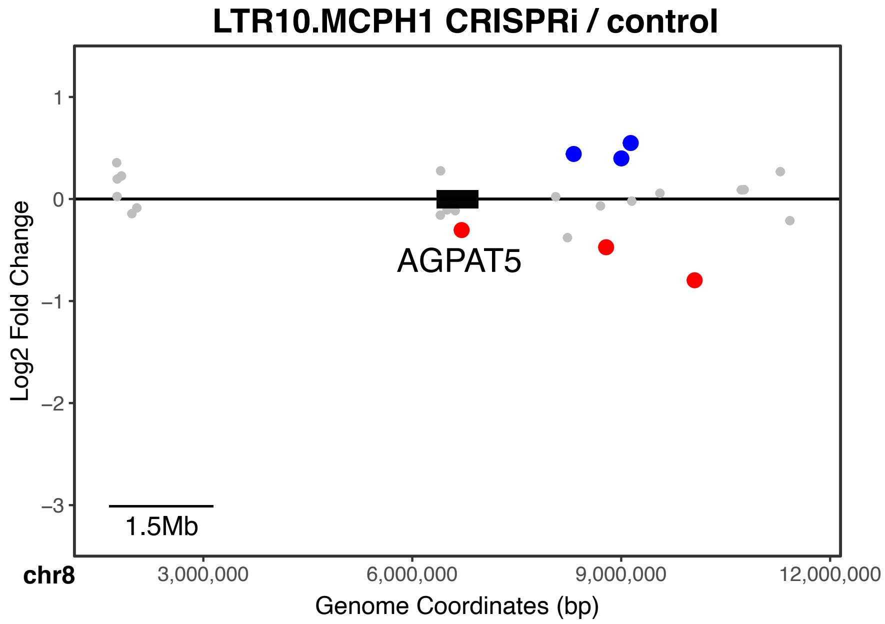


**Figure S4F:** Scatterplot of local gene expression changes in response to CRISPR silencing of the LTR10.MCPH1 element. Significantly downregulated genes are shown in red; significantly upregulated genes are shown in blue. Significantly downregulated genes within 1.5 MB of the targeted element are labeled (element box not drawn to scale).

#### Figure S4G

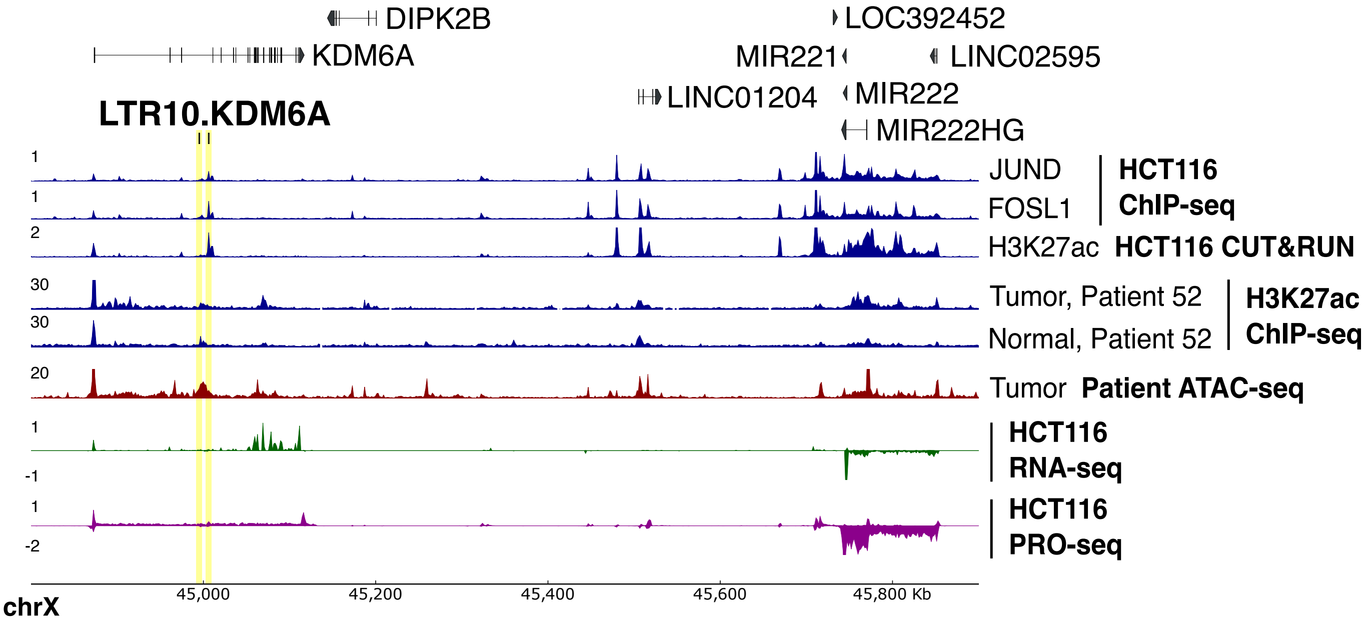


**Figure S4G:** Genome browser screenshot of the *KDM6A* locus with the LTR10.KDM6A enhancer labeled. From top to bottom: JUND and FOSL1 ChIP-seq (GSE32465), H3K27ac CUT&RUN (in-house), H3K27ac ChIP-seq from matched tumor/normal samples from the CEMT Canadian Epigenome Project (patient AKCC52), tumor ATAC-seq from TCGA (patient COAD P022), HCT116 RNA-seq (in-house), and HCT116 PRO-seq (GSE129501). Axis numbers represent the upper limit of the range; the lower limit is always zero.

#### Figure S4H

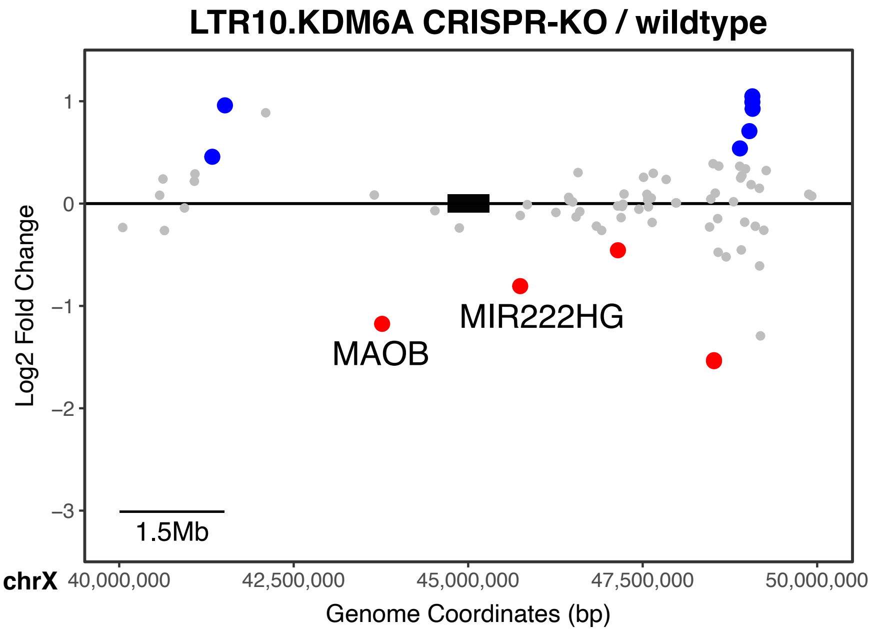


**Figure S4H:** Scatterplot of local gene expression changes in response to CRISPR deletion of the LTR10.KDM6A element. Multiple CRISPR deletion KO clones were generated and compared against wildtype HCT116 cells. Significantly downregulated genes are shown in red; significantly upregulated genes are shown in blue. Significantly downregulated genes within 1.5 MB of the targeted element are labeled (element box not drawn to scale).

#### Figure S4I

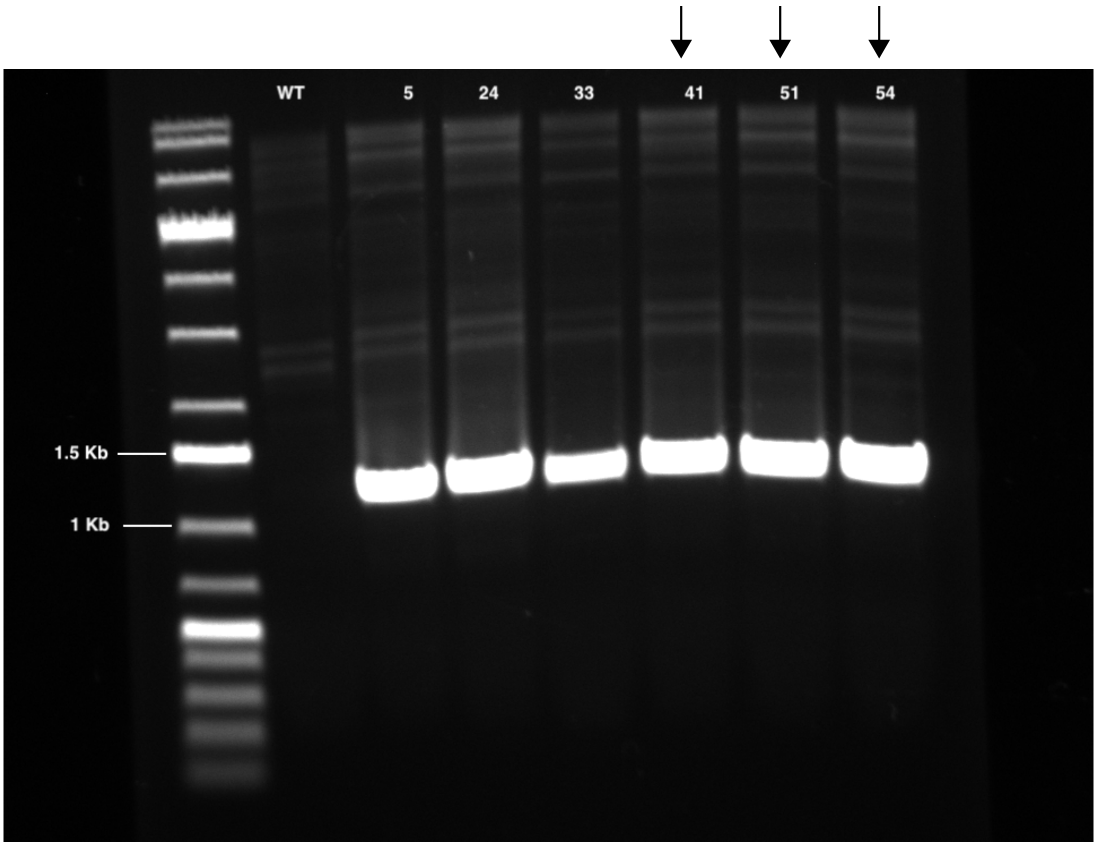


**Figure S4I:** External PCR validation of LTR10.KDM6A CRISPR KO clones. PCR primers used for validation (KDM6A_up_external, KDM6A_down_external) flank LTR10.KDM6A element. Expected KO amplicon size is 1245 bp. Expected wildtype amplicon size (unobserved) is 8854 bp. Arrows indicate clones that were used for DEseq2 analysis against wildtype HCT116 cells, based on PCA results.

#### Figure S4J

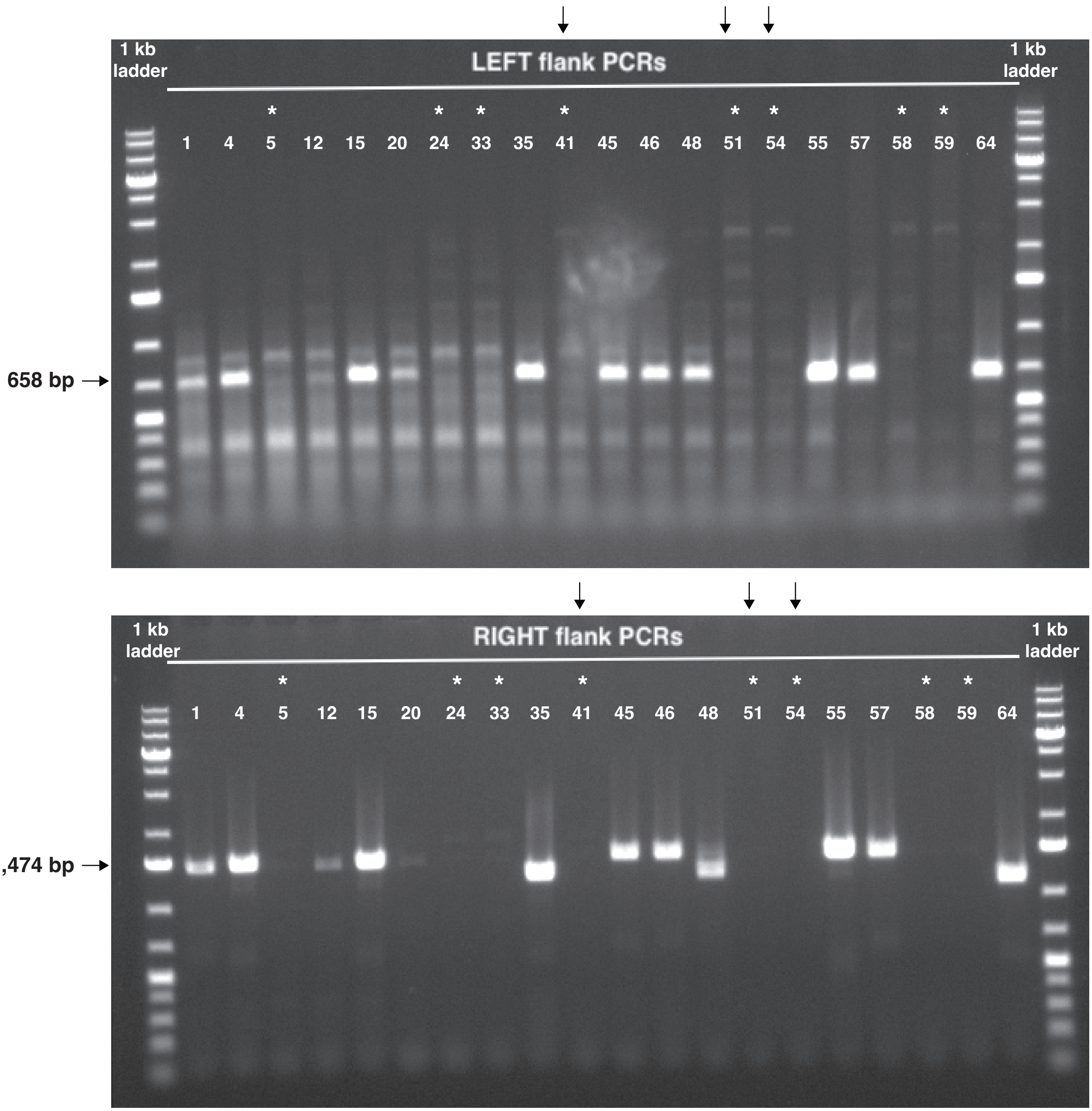


**Figure S4J:** Internal PCR validation of LTR10.KDM6A CRISPR KO clones. Expected wildtype amplicon size is 658 bp for upstream PCR (left flank), 1474 bp for downstream PCR (right flank). Asterisks indicate homozygous KO clones, based on no product from both flanks. Arrows indicate clones that were used for DEseq2 analysis against wildtype HCT116 cells, based on PCA results.

#### Figure S4K

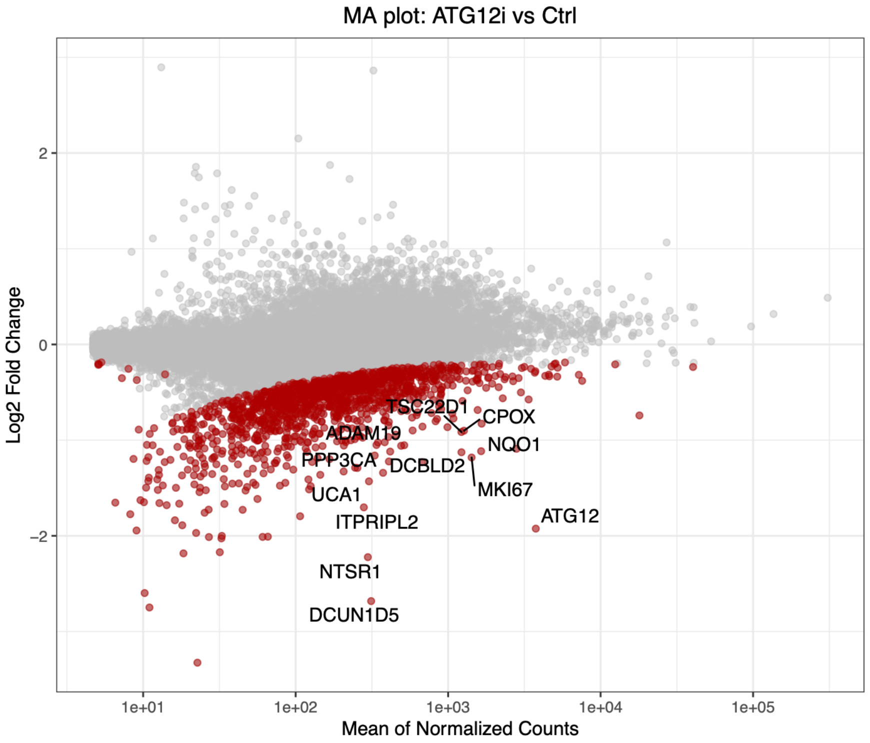


**Figure S4K:** MA plot showing global gene expression changes in cells in response to silencing ATG12 (TSS). Significantly downregulated genes are shown in red.

#### Figure S4L

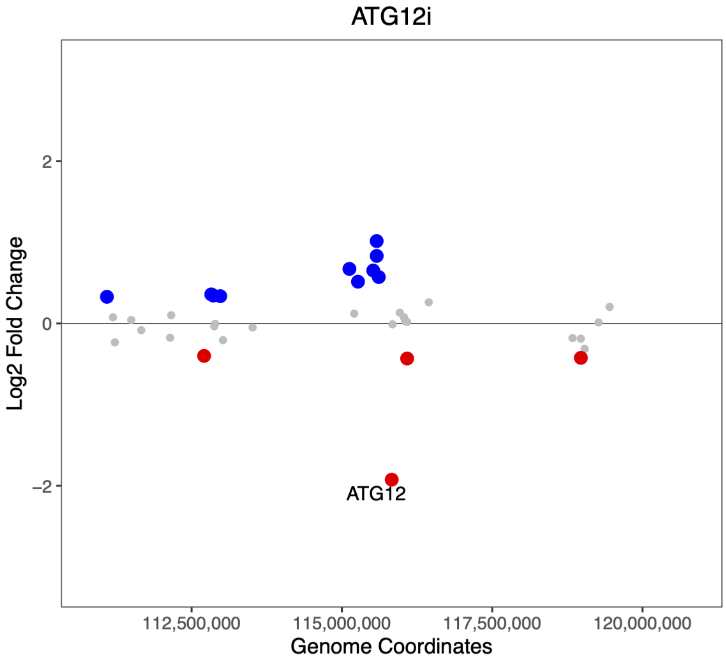


**Figure S4L:** Scatterplot of gene expression changes in response to silencing ATG12 (TSS) in a 10Mb window around the target. Significantly down-regulated genes are shown in red, significantly up-regulated genes are shown in blue. The most significantly downregulated gene (ATG12) is labelled.

#### Figure S4M

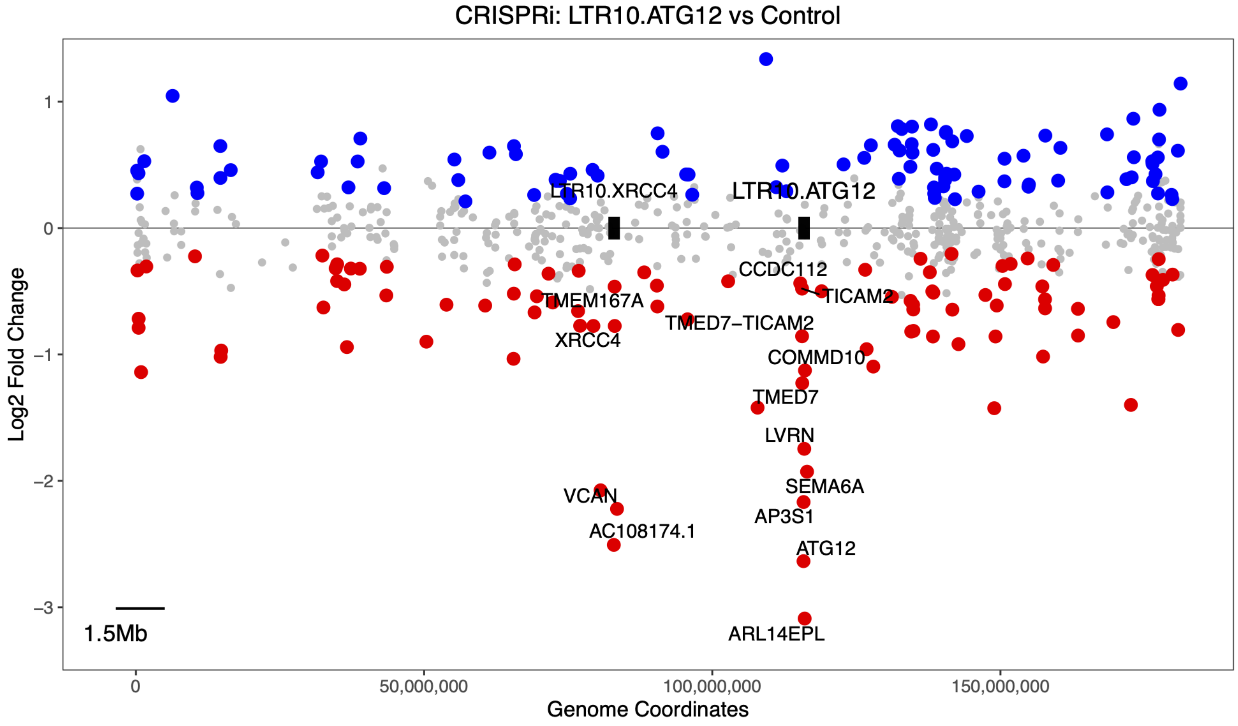


**Figure S4M:** Scatterplot of gene expression changes in response to silencing LTR10.ATG12, showing all of hg38 chromosome 5. The locations of LTR10.XRCC4 and LTR10.ATG12 are marked (element boxes not drawn to scale). Significantly downregulated genes are shown in red; significantly upregulated genes are shown in blue. Significantly downregulated genes within 1.5 Mb of either LTR10 element are labelled.

#### Figure S4N

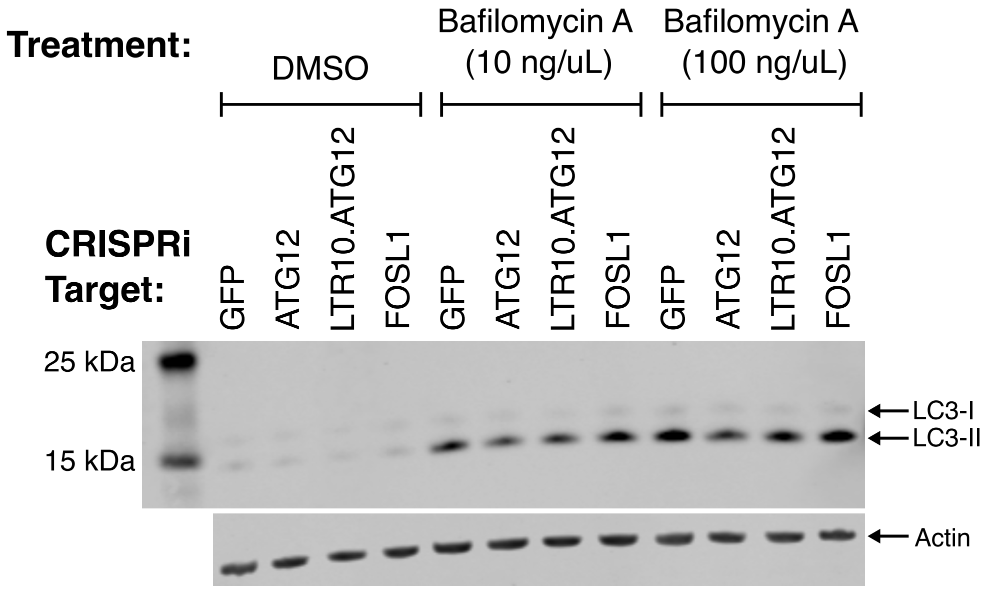


**Figure S4N:** Immunoblot of LC3-I and LC3-II in each CRISPRi cell line, after treating cells for 6 hr with DMSO, bafilomycin A (10 ng/uL), or bafilomycin A (100 ng/uL).

#### Figure S5A

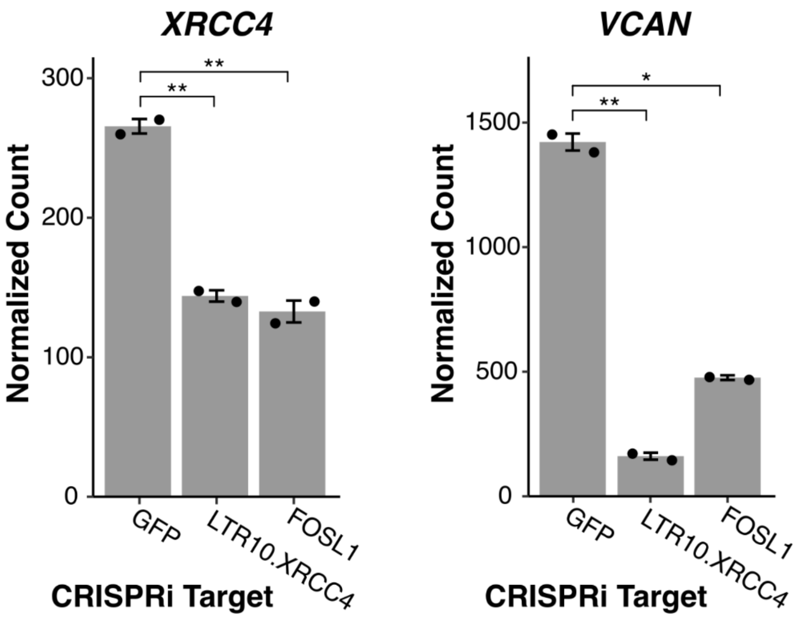


**Figure S5A:** Normalized RNA-seq expression values of *XRCC4* and *VCAN* in dCas9-KRAB-MeCP2 HCT116 cells stably transfected with gRNAs targeting the LTR10.XRCC4 element, the *FOSL1* transcription start site, or non-targeting (GFP) control.

#### Figure S5B

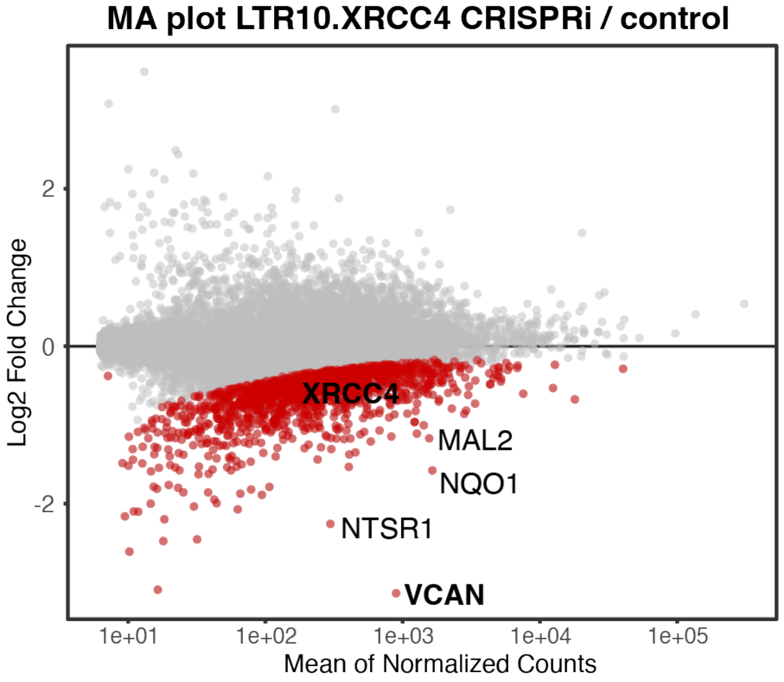


**Figure S5B:** MA plot showing global gene expression changes in cells in response to silencing LTR10.XRCC4. Significantly downregulated genes are shown in red.

### Figure S5C

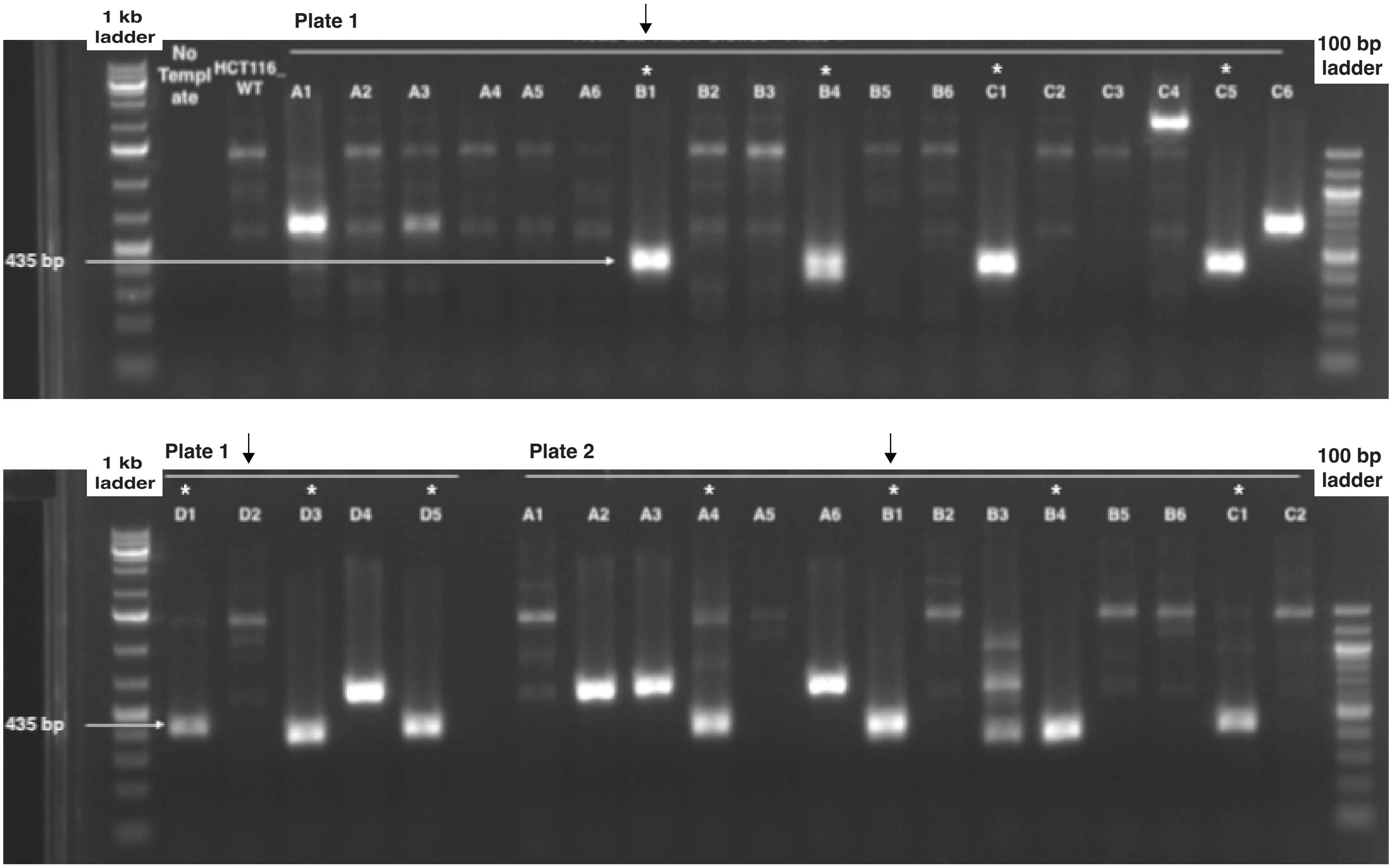


**Figure S5C:** External PCR validation of LTR10.XRCC4 CRISPR KO clones. PCR primers used for validation (XRCC4_up_external, XRCC4_down_external) flank LTR10.XRCC4 enhancer. Expected KO amplicon size is 435 bp. Expected wildtype amplicon size is 7763 bp. Asterisks indicate clones with 435 bp dominant product, indicative of a KO. Several clones showed no product at 435 bp and were propagated to serve as negative controls. Arrows indicate clones that were used for irradiation and mouse xenograft experiments: wildtype D2 and knockout B1.

#### Figure S5D

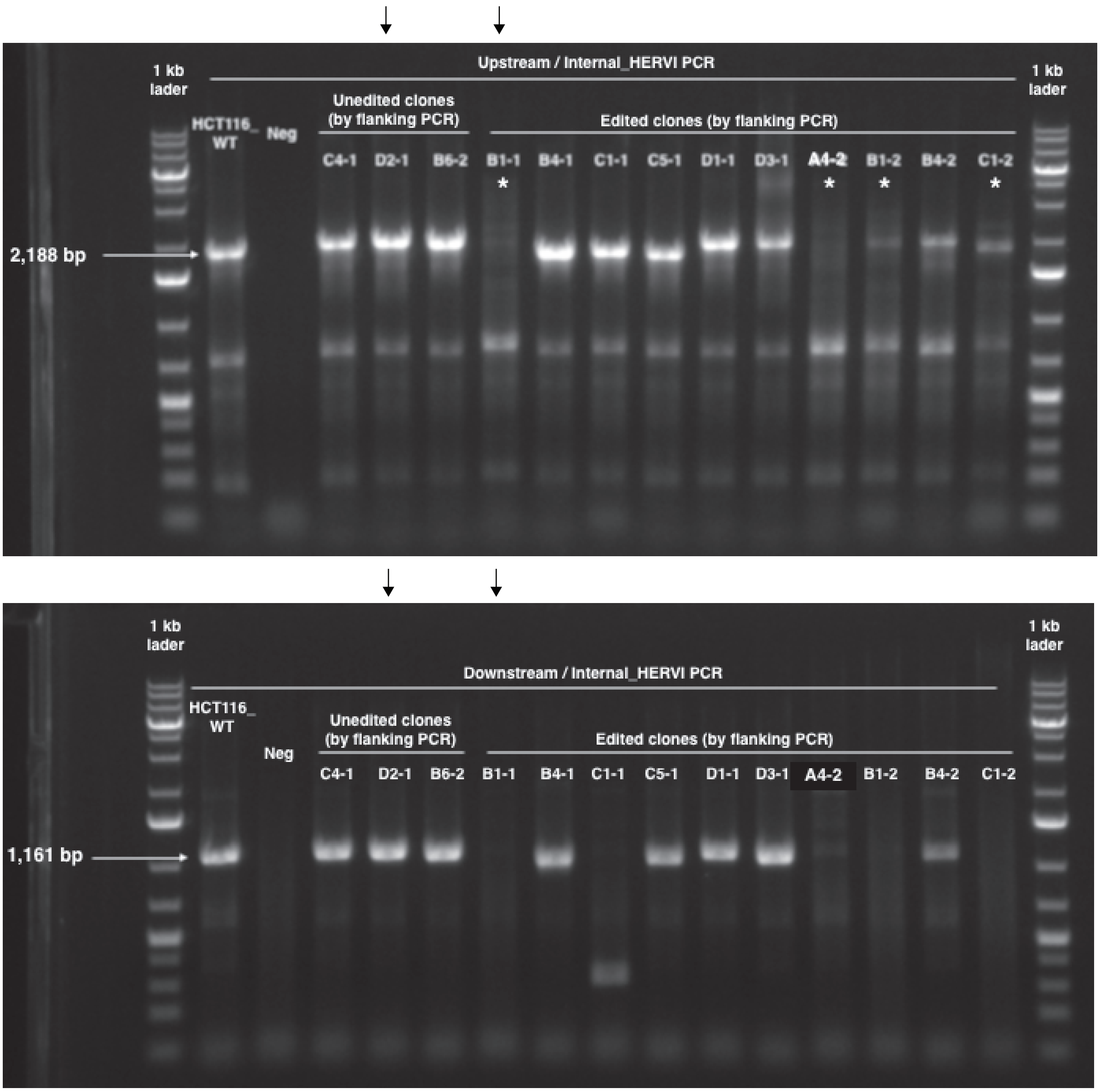


**Figure S5D:** Internal PCR validation of LTR10.XRCC4 CRISPR KO clones. Expected wildtype amplicon size is 2188 bp for upstream PCR, 1161 bp for downstream PCR. Asterisks indicate KO clones. Clones B1-1 and A4-2 showed no product in both reactions, indicative of homozygous KO clones. Clones B1-2 and C1-2 lacked product in the downstream reaction but showed a faint product in the upstream reaction. Arrows indicate clones that were used for irradiation and mouse xenograft experiments: wildtype D2-1 and knockout B1-1.

#### Figure S5E

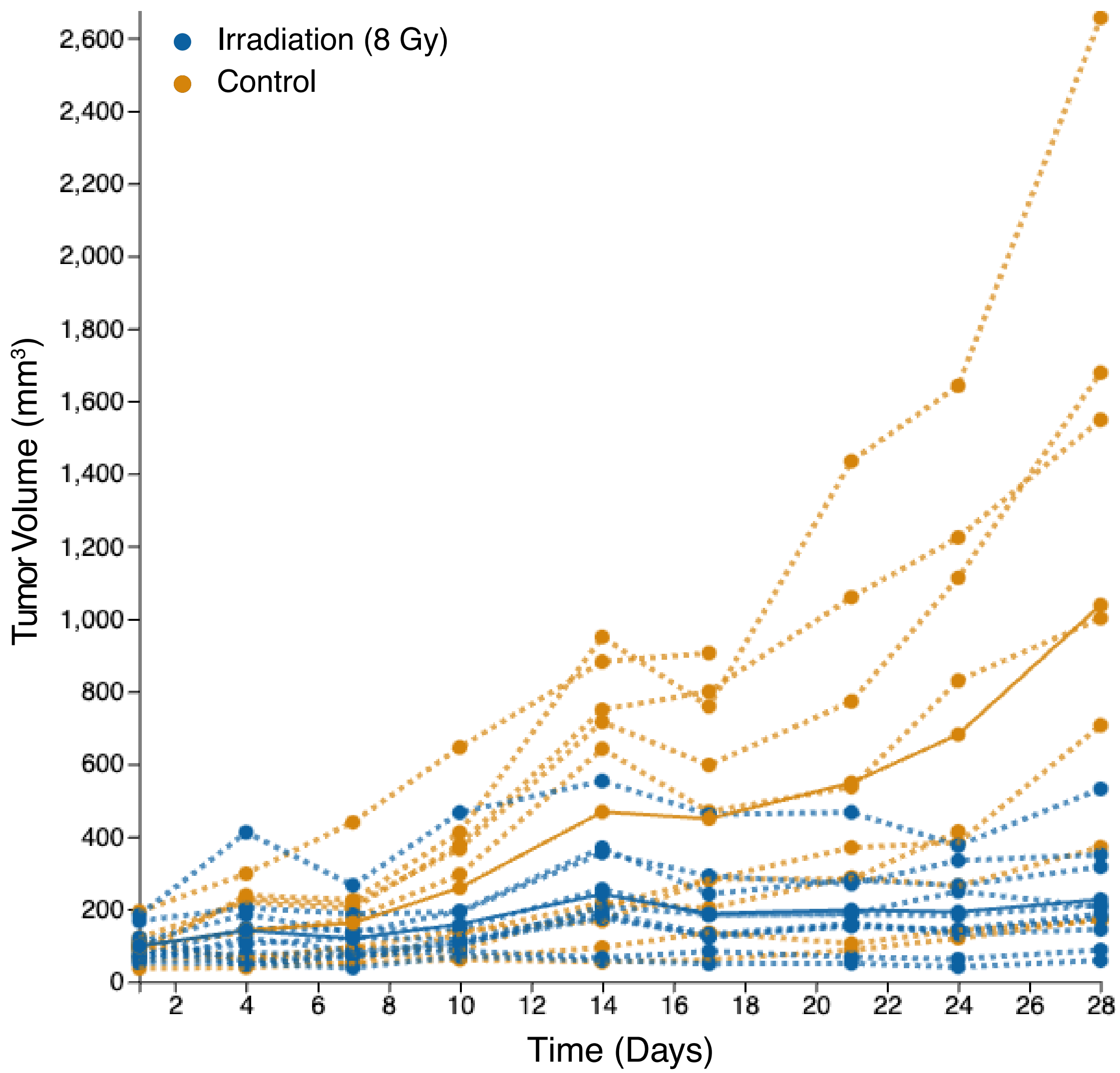


**Figure S5E:** Individual growth curves across replicates for wildtype HCT116 xenograft tumors, with irradiation (n=10) and without irradiation (n=10), for 28 days.

#### Figure S5F

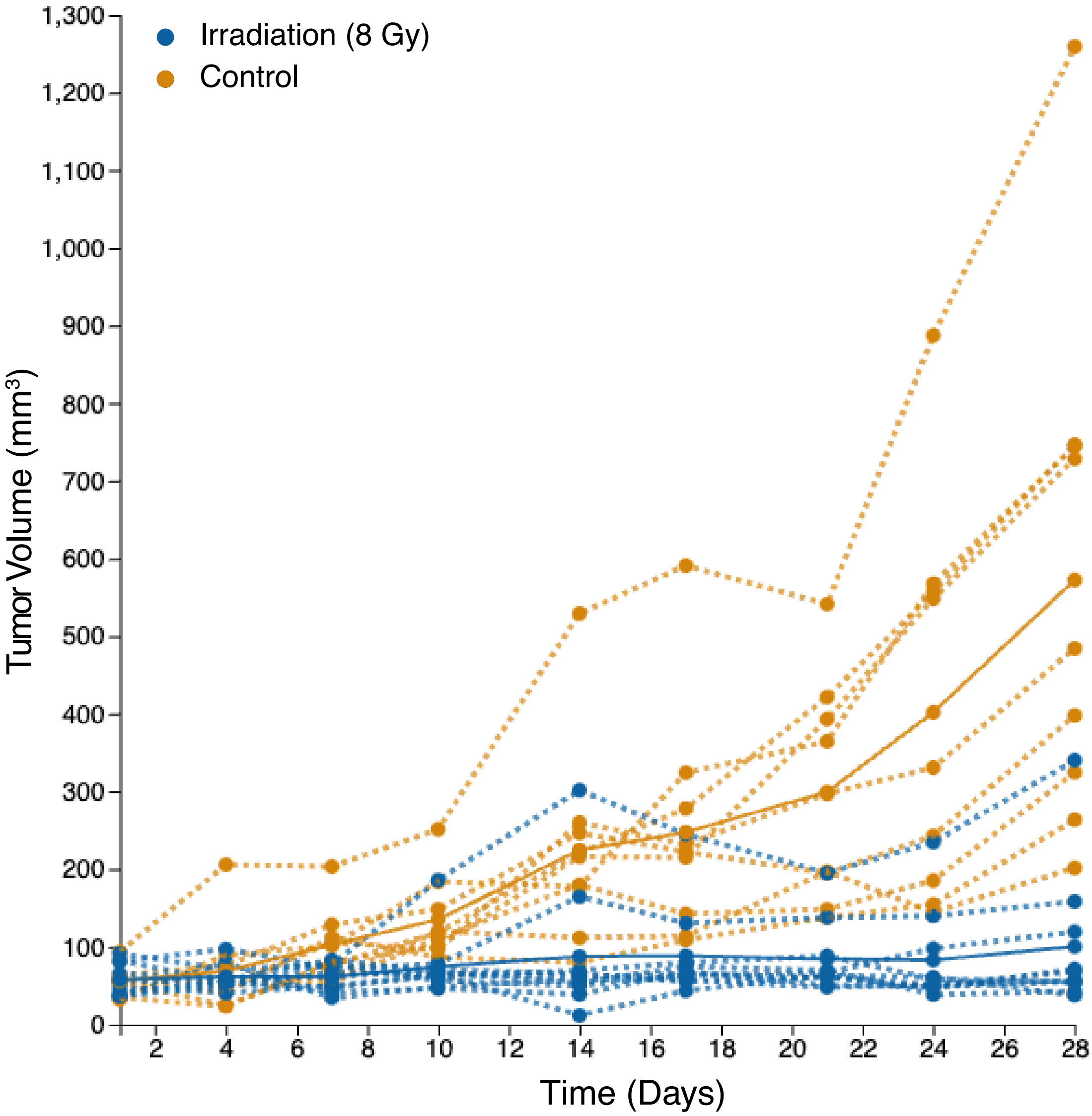


**Figure S5F:** Individual growth curves across replicates for LTR10.XRCC4 knockout HCT116 xenograft tumors, with irradiation (n=10) and without irradiation (n=9), for 28 days.

**Figure S5G**

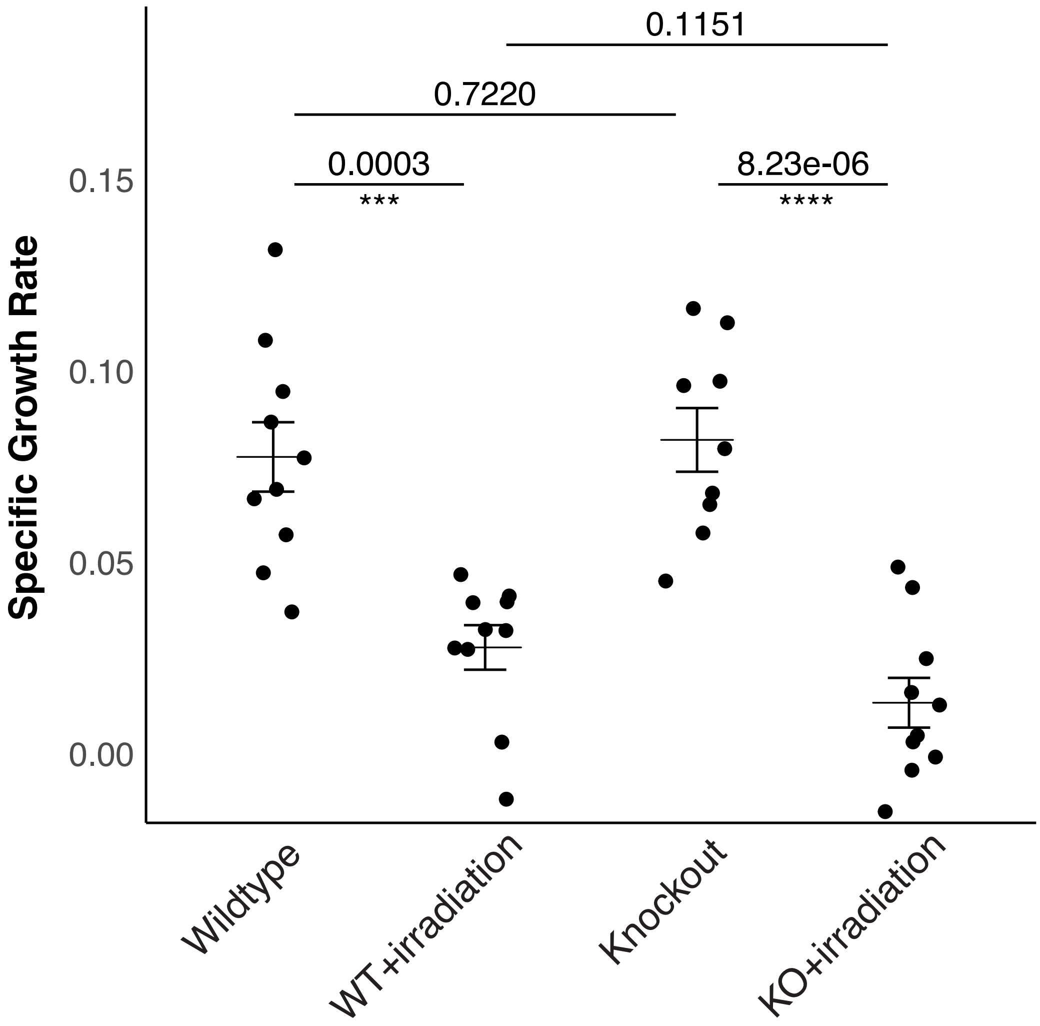


**Figure S5G:** Specific growth rates for each HCT116 xenograft tumor, separated by group: wildtype (n=10), wildtype + irradiation (n=10), LTR10.XRCC4 knockout (n=9), and LTR10.XRCC4 knockout + irradiation (n=10). Welch's t-test p-values are shown for each comparison.

#### Figure S6A

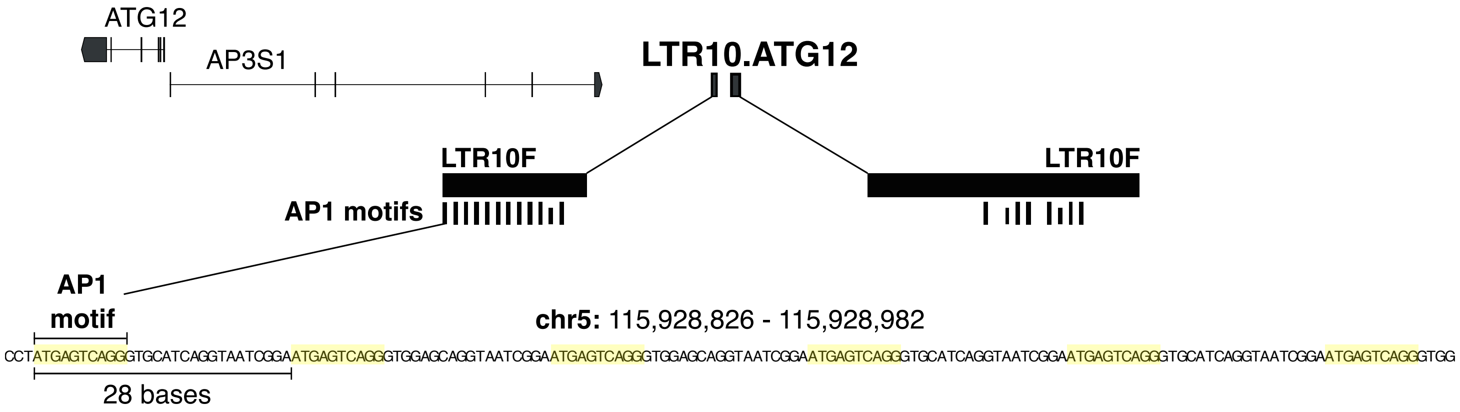


**Figure S6A:** Genome browser screenshot of LTR10.ATG12 showing a close-up of the 28 bp tandem repeat region that includes the AP1 motif.

#### Figure S6B

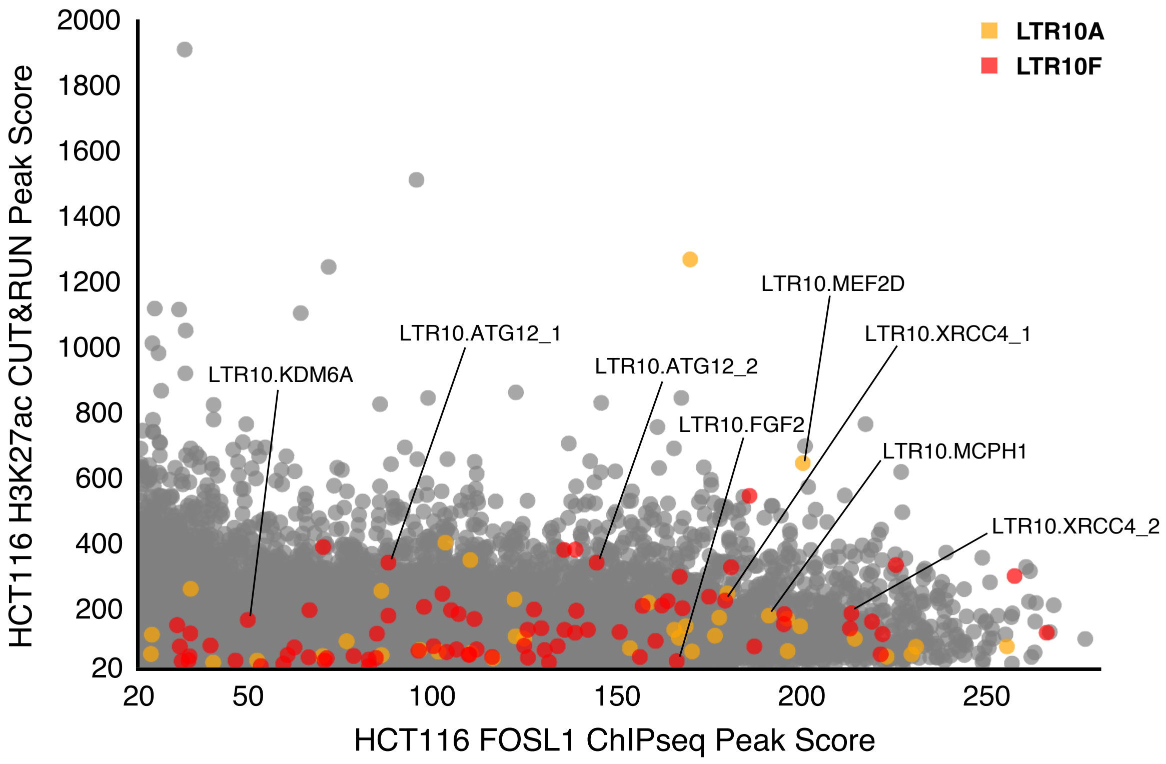


**Figure S6B:** Scatterplot of H3K27ac and FOSL1 peak scores for all HCT116 peak regions that have both marks. HCT116 H3K27ac peaks (in-house CUT&RUN, filtered for peak score over 20) were intersected with HCT116 FOSL1 peaks (public ChIP-seq from GSE32465, filtered for peak score over 20) to define a list of genomic peaks with both marks (n=19,315). LTR10A/F elements were then intersected with these H3K27ac+FOSL1 peak regions. 143 distinct LTR10A/F elements contribute to 115 of these peaks. LTR10A-derived peaks are shown in orange, LTR10F-derived peaks are shown in red, and all other genomic H3K27ac+FOSL1 peaks are shown in grey. CRISPR-validated LTR10 enhancers are labeled by name.

#### Figure S6C

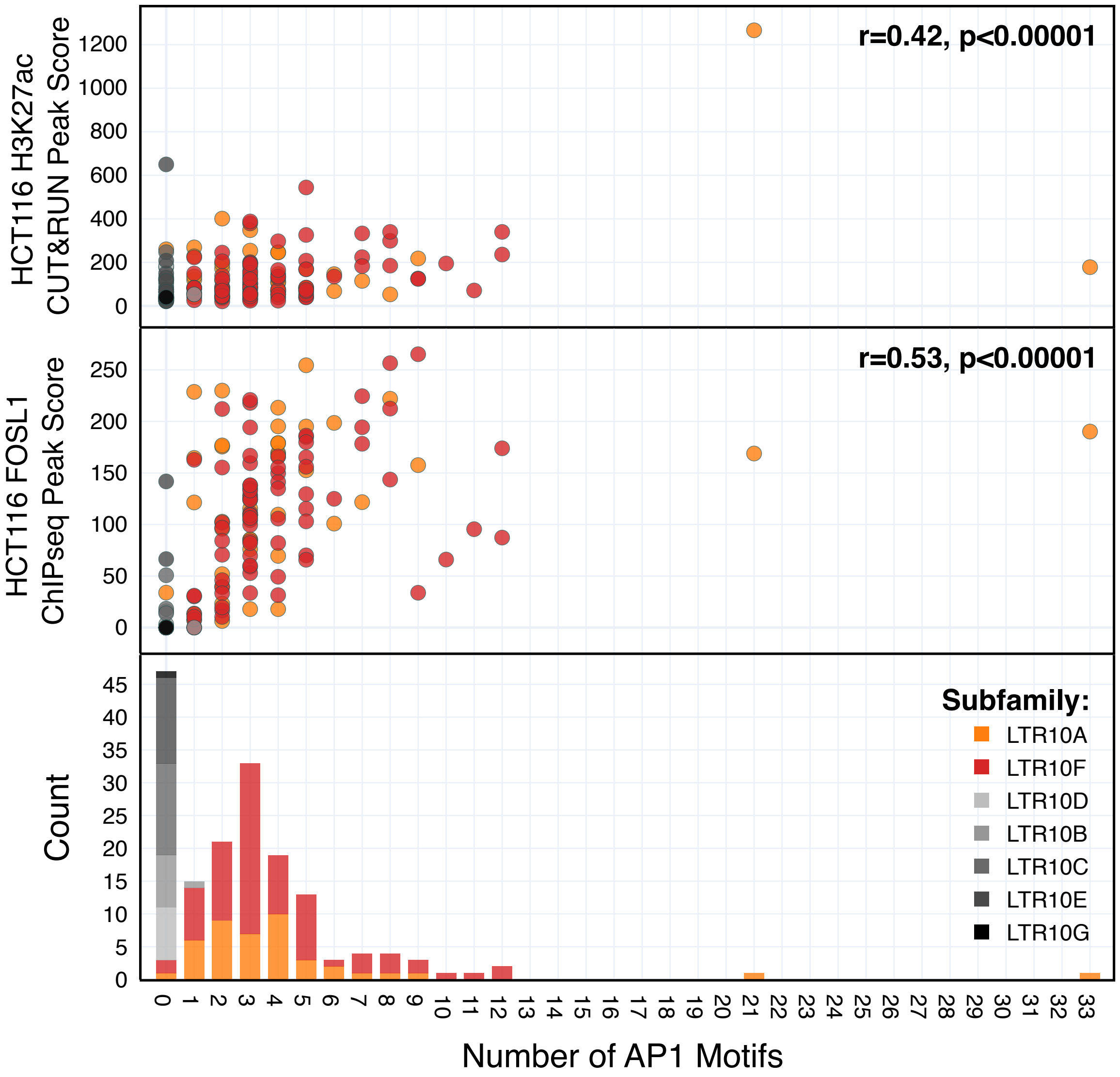


**Figure S6C:** Scatterplots showing correlations between the number of AP1 motifs with HCT116 H3K27ac signal (in-house CUT&RUN) and HCT116 FOSL1 signal (public ChIP-seq from GSE32465). Only LTR10 elements marked by H3K27ac are included. The Pearson correlation coefficient (r) and p-value are shown for each plot.

#### Figure S6D

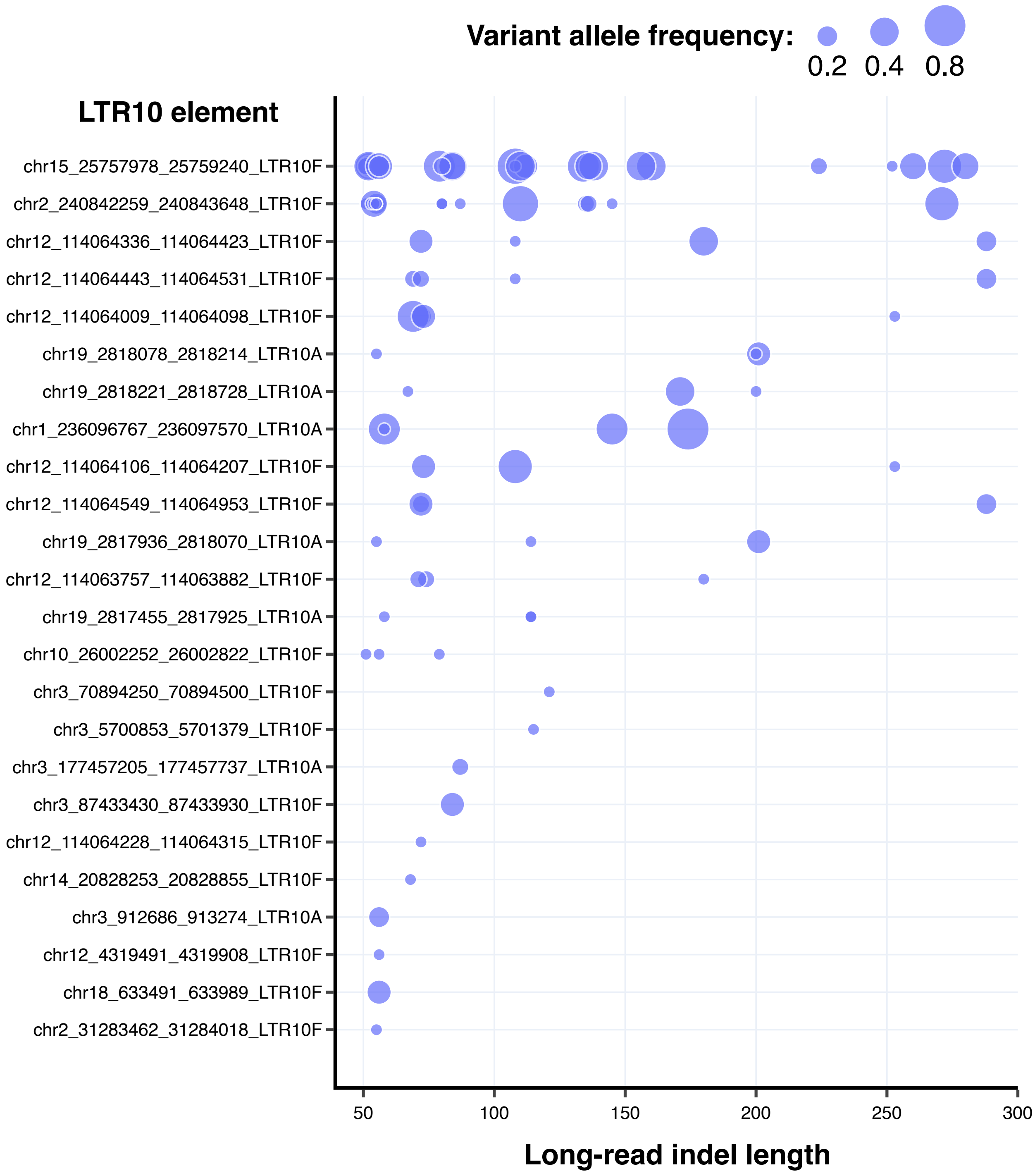


**Figure S6D:** Scatterplot of long-read indels between 50-300 bp in length overlapping LTR10A or LTR10F elements. Each dot represents a distinct indel, plotted by its length and variant allele frequency (bubble size). Indels were derived from long-read structural variant calls generated from 15 individuals (Audano et al. 2019).

#### Figure S6E

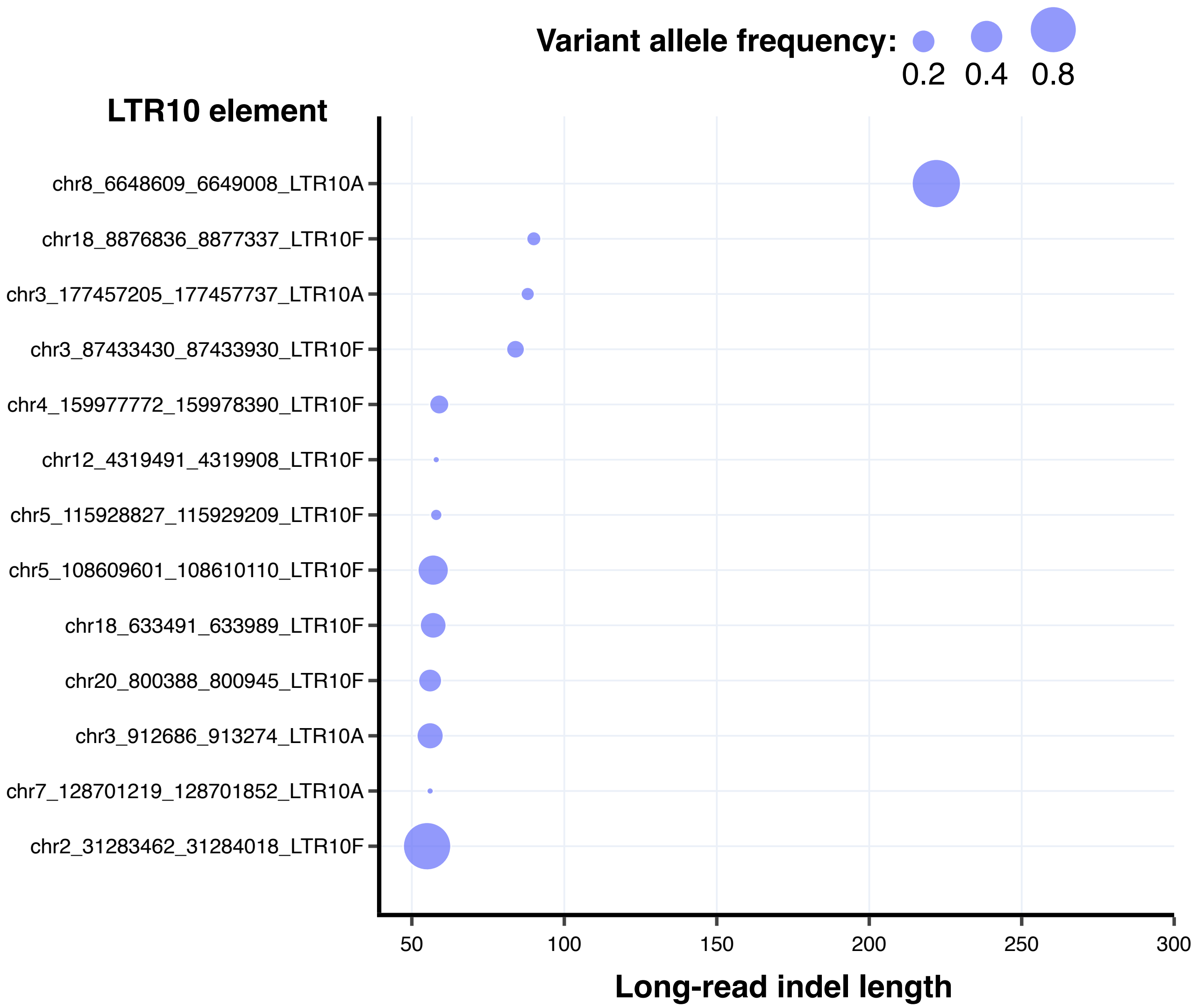


**Figure S6E:** Scatterplot of long-read indels between 50-300 bp in length overlapping LTR10A or LTR10F elements. Each dot represents a distinct indel, plotted by its length and variant allele frequency (bubble size). Indels were derived from long-read structural variant calls generated from 25 individuals (Quan et al. 2021).

### Figure S6F

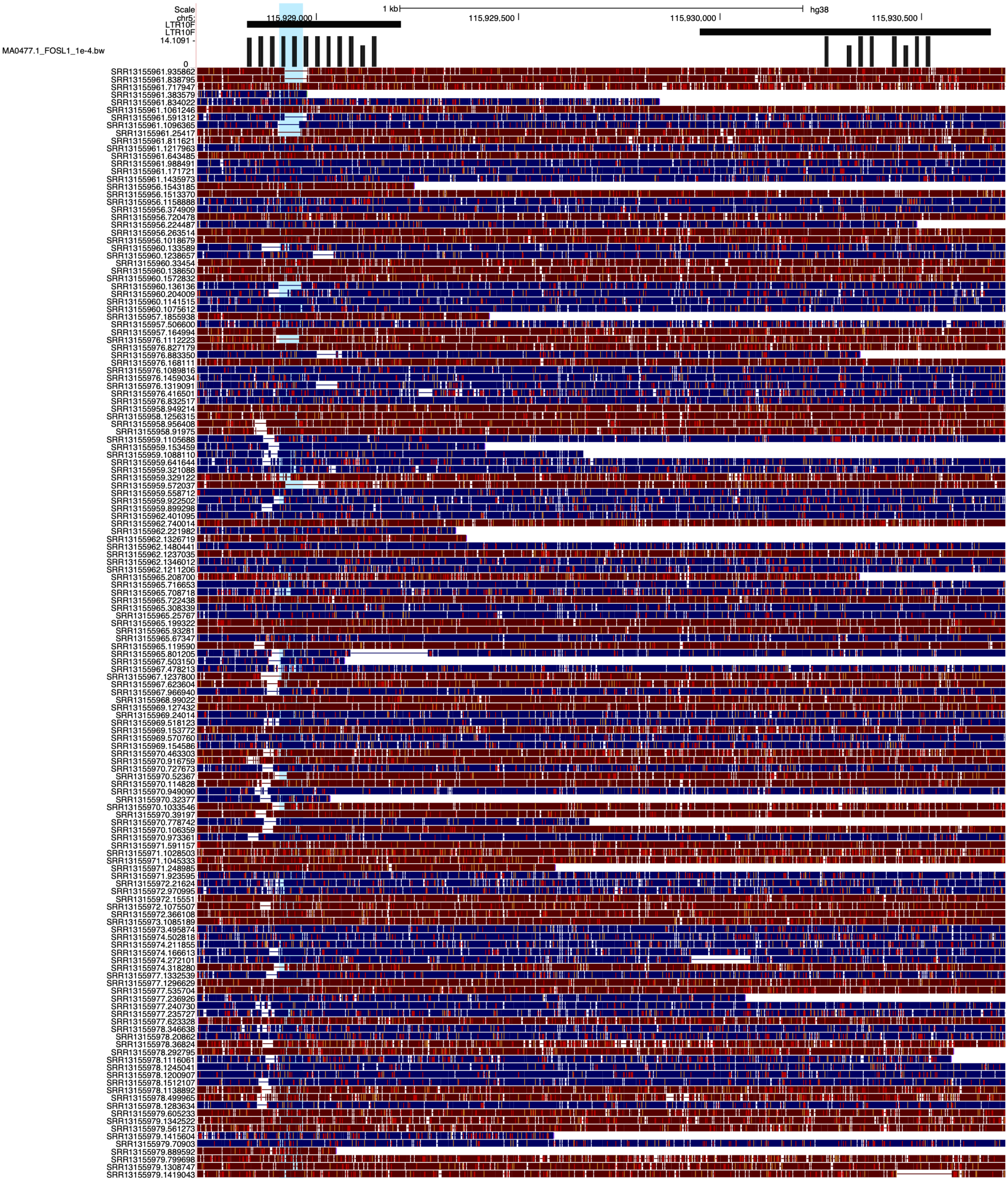


**Figure S6F:** Genome browser screenshot of a deletion within the LTR10.ATG12 element, showing reads from a long-read dataset of 25 Asian individuals (Quan et al. 2021). The blue highlight shows the location of the reported deletion.

### Figure S6G

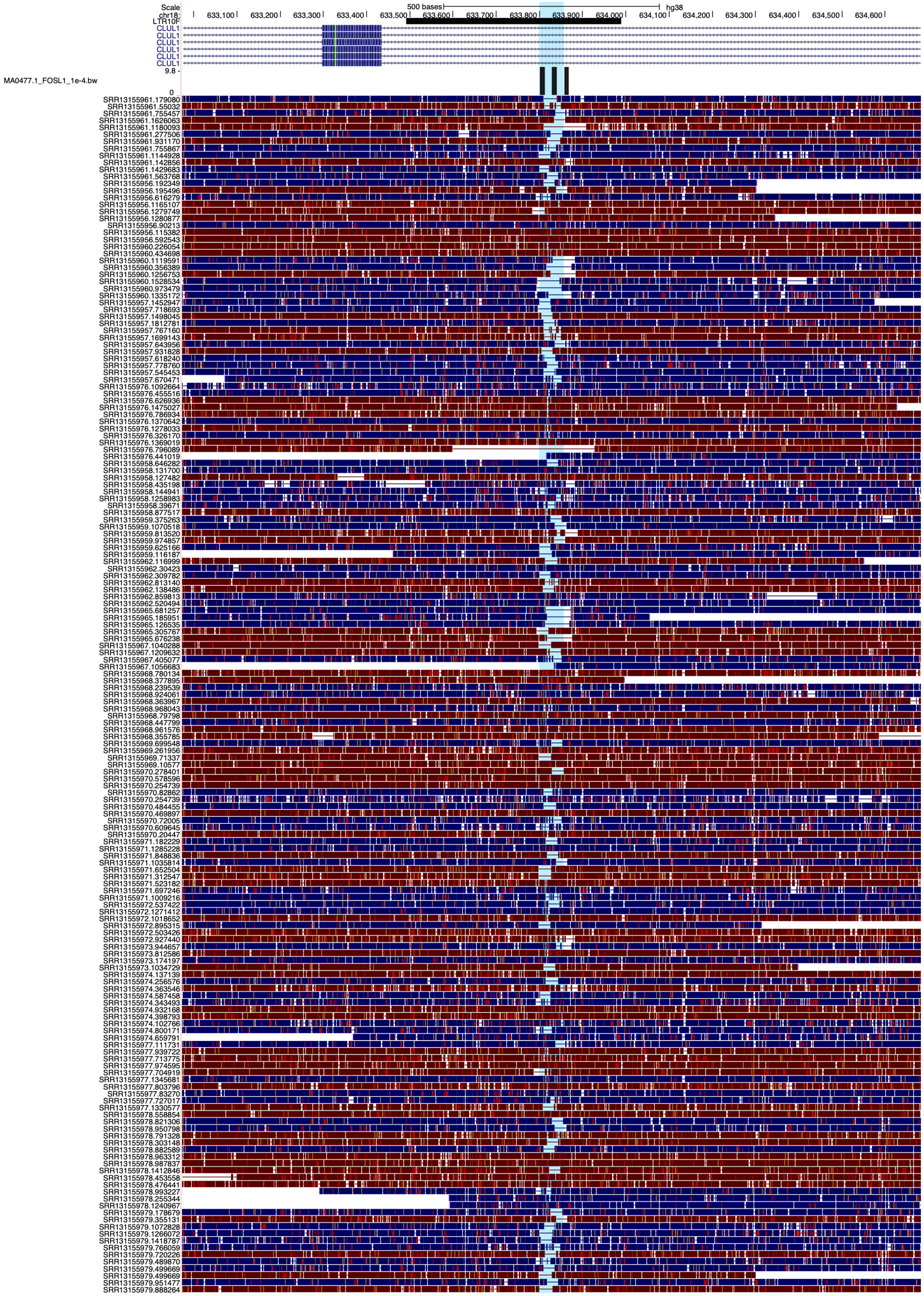

**Figure S6G:** Genome browser screenshot of a deletion within an LTR10F element within the CLUL1 gene, showing reads from a long-read dataset of 25 Asian individuals (Quan et al. 2021). The blue highlight shows the location of the reported deletion.

### Figure S6H

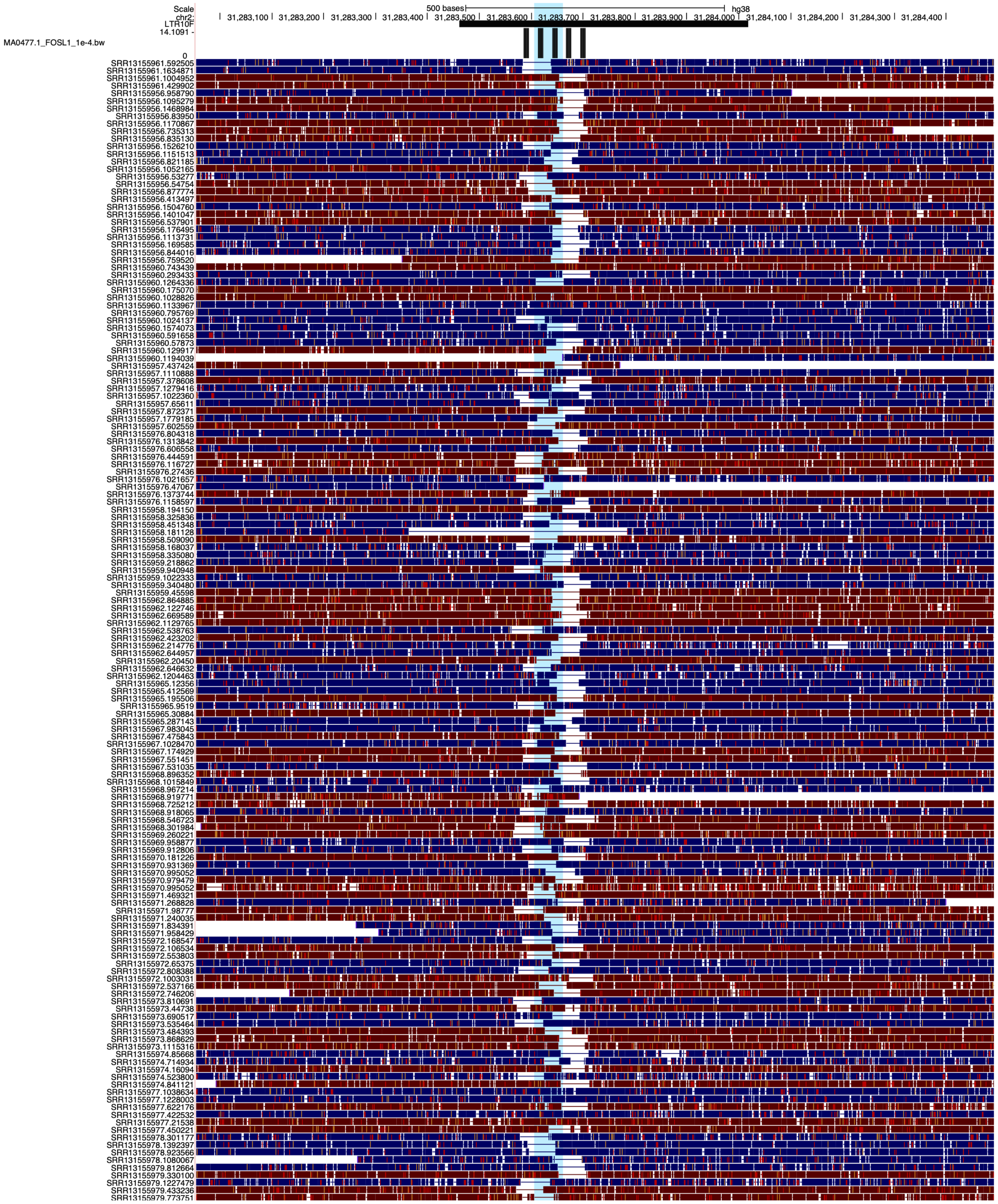


**Figure S6H:** Genome browser screenshot of a chr2 deletion within an LTR10F element, showing reads from a long-read dataset of 25 Asian individuals (Quan et al. 2021). The blue highlight shows the location of the reported deletion.

### Figure S6I

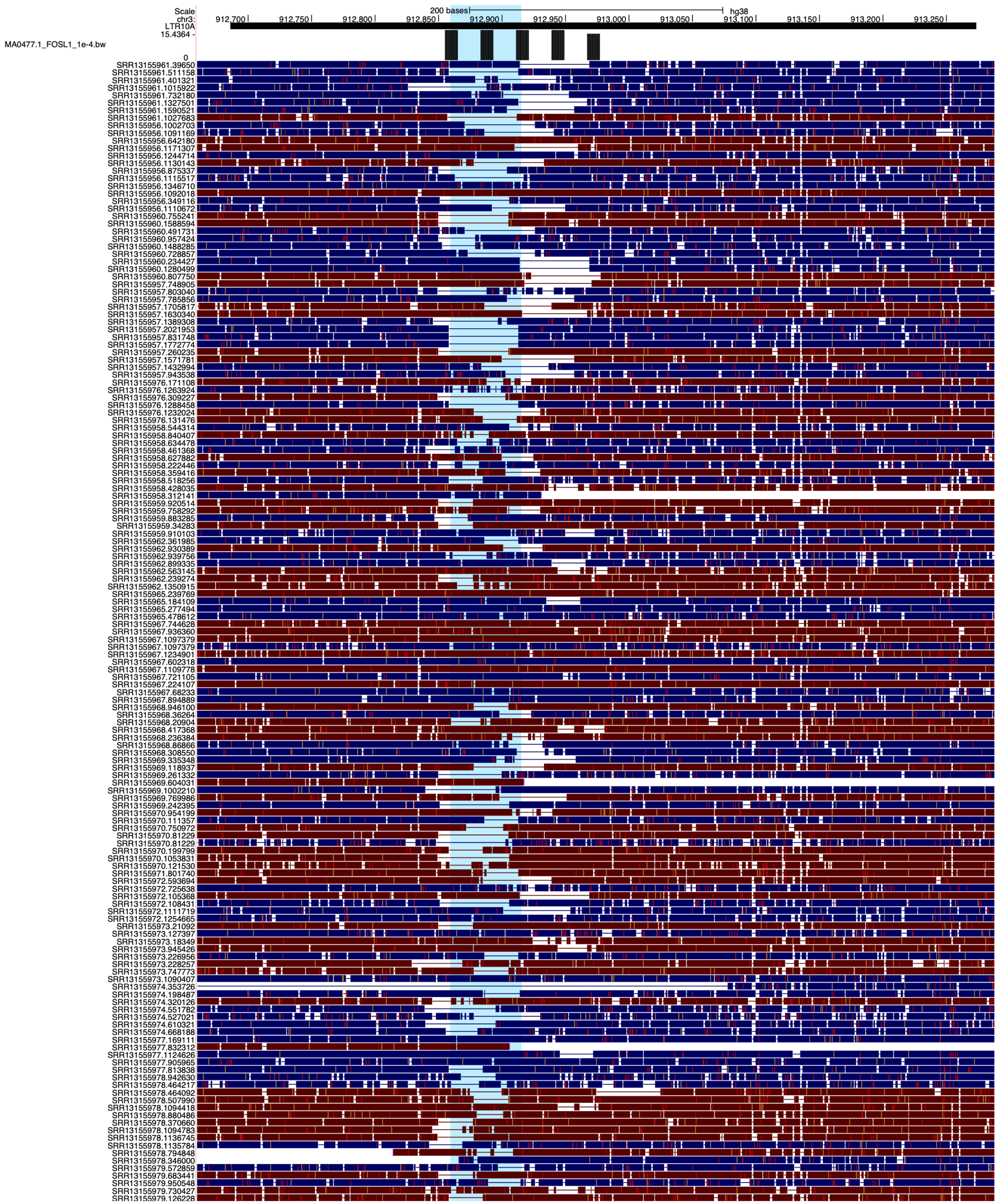

**Figure S6I:** Genome browser screenshot of a chr3 deletion within an LTR10A element, showing reads from a long-read dataset of 25 Asian individuals (Quan et al. 2021). The blue highlight shows the location of the reported deletion.

### Figure S6J

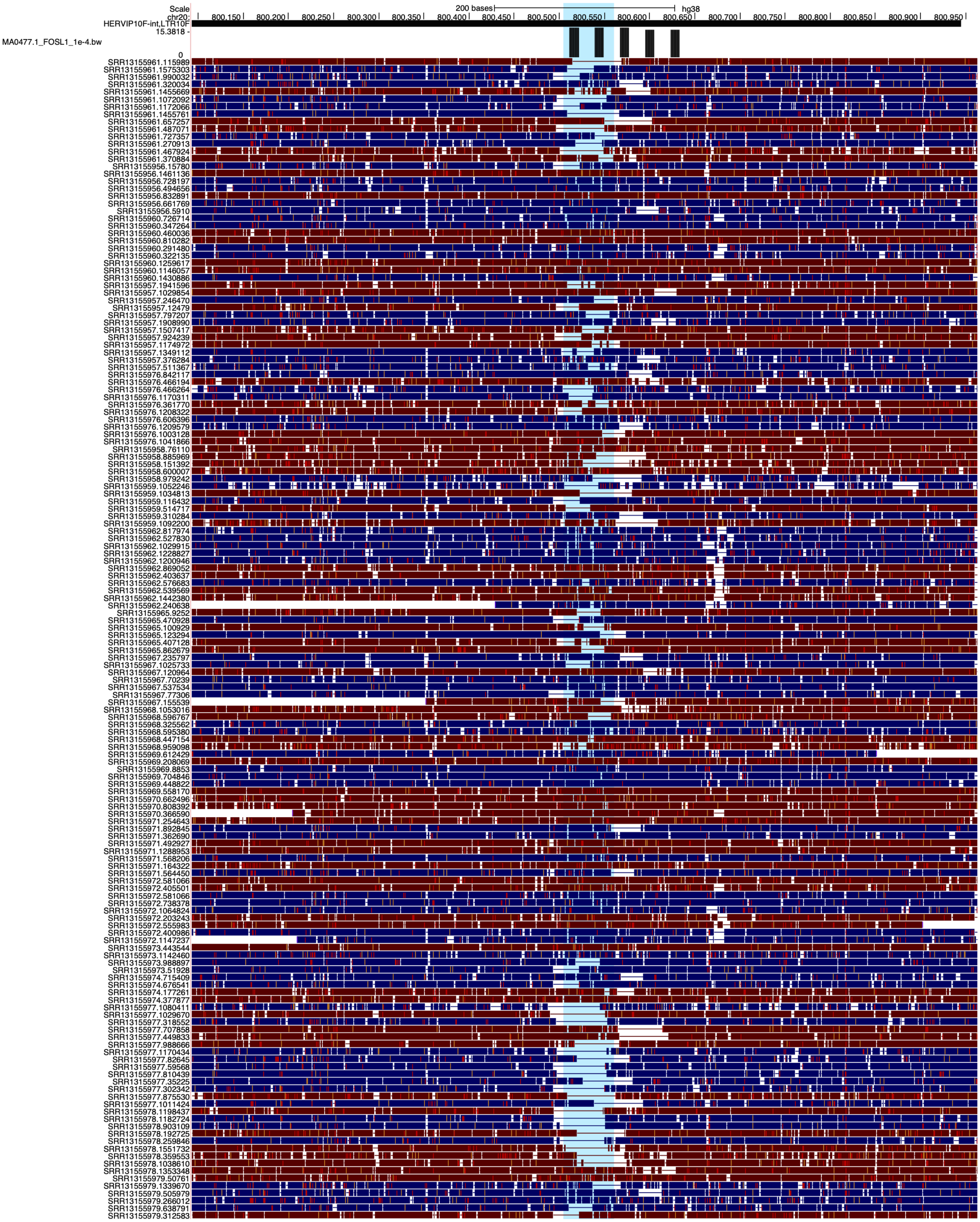


**Figure S6J:** Genome browser screenshot of a chr20 deletion within an LTR10F element, showing reads from a long-read dataset of 25 Asian individuals (Quan et al. 2021). The blue highlight shows the location of the reported deletion.

#### Figure S6K

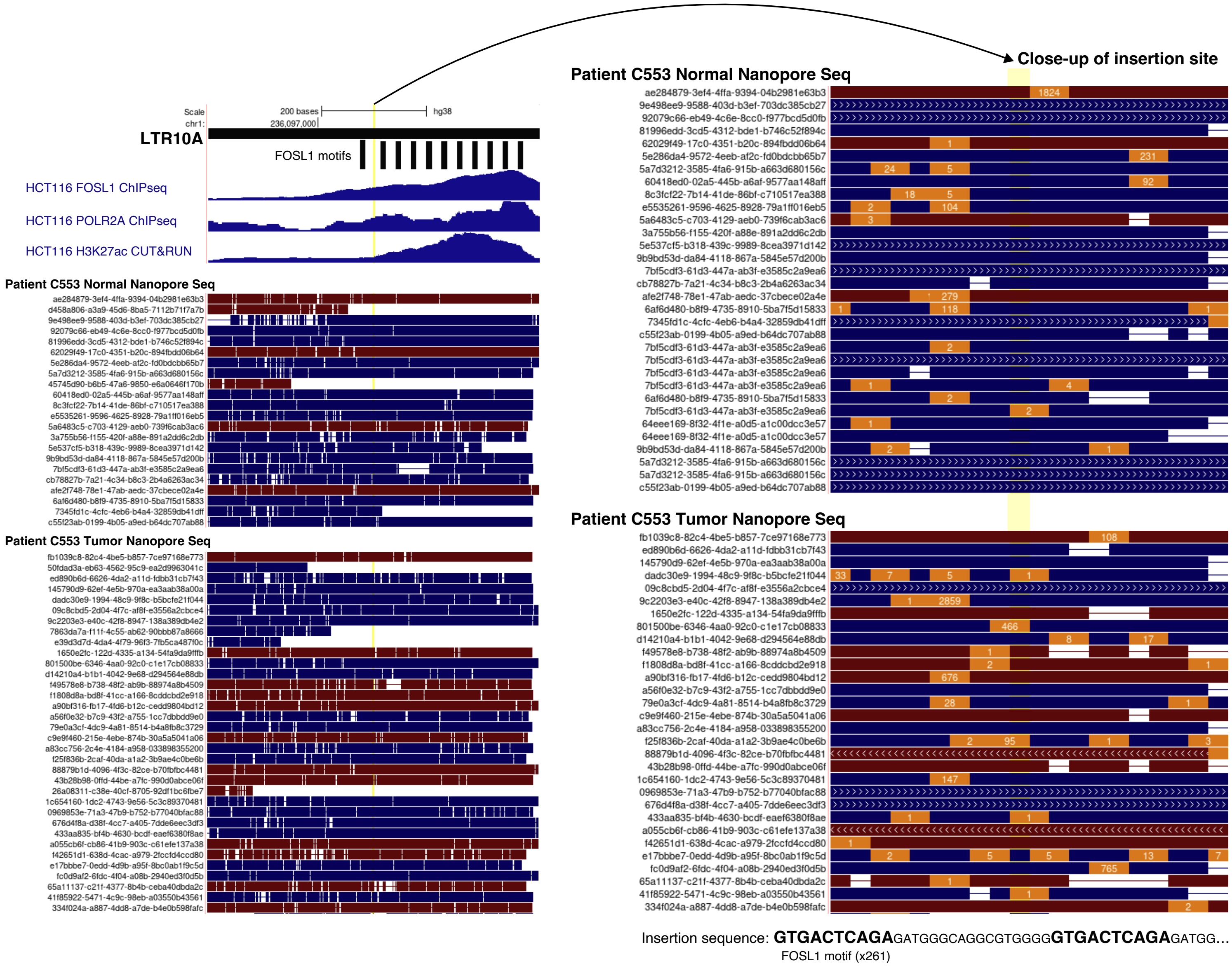


**Figure S6K:** Genome browser screenshots of a tumor-specific LTR10A VNTR expansion on chr1. Matched tumor/normal reads from patient C553 are derived from a long-read nanopore sequencing dataset of 20 patients with colorectal adenocarcinomas (Xu et al. 2023). The yellow highlight shows the approximate location of the predicted tumor-specific mosaic expansion, as called by Sniffles2 (Smolka et al. 2023). A close-up of reads at the insertion site is shown on the right, with orange boxes indicating insertions and numbers within orange boxes indicating insertion sequence lengths (in bp).

### Figure S6L

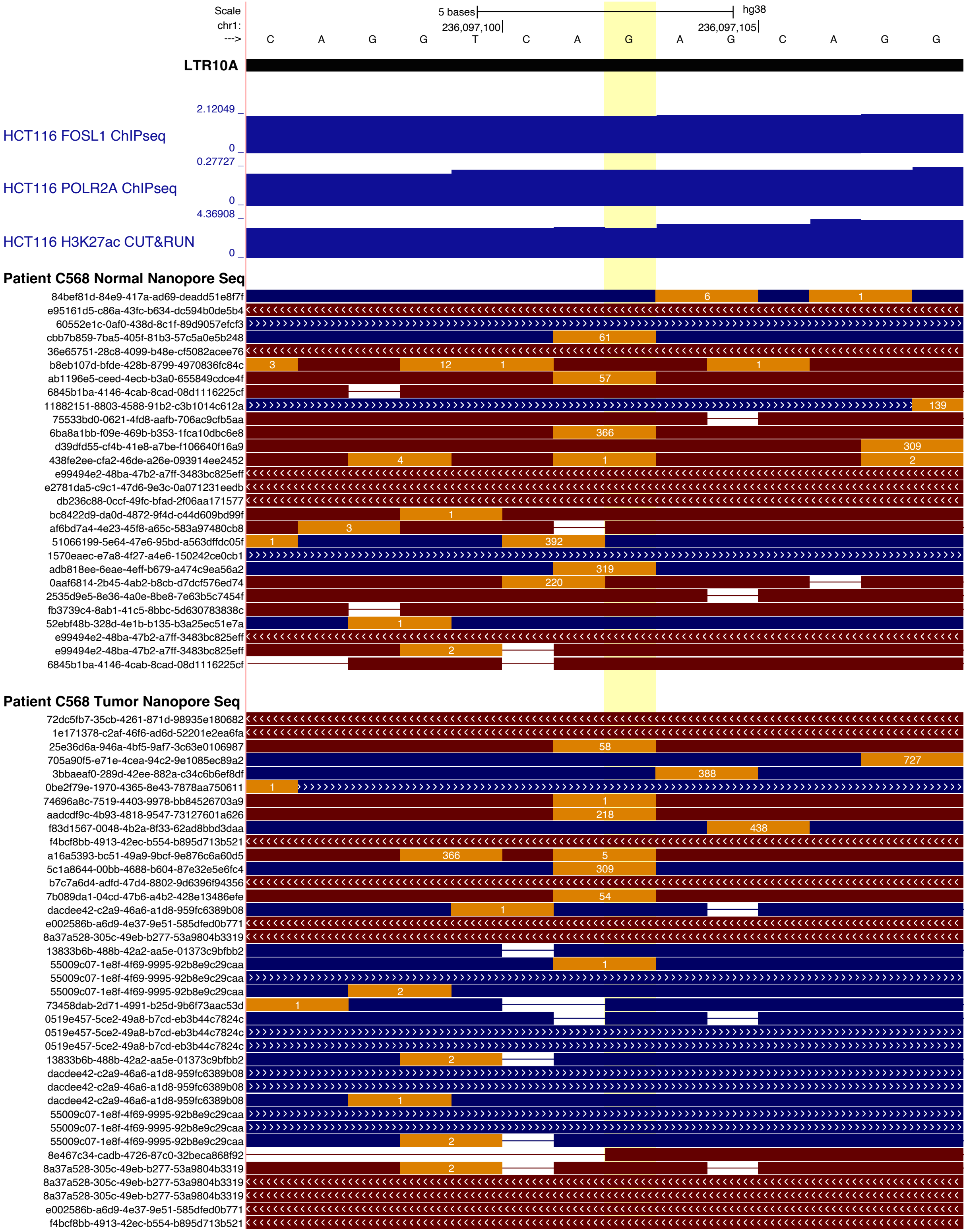


**Figure S6L:** Genome browser screenshot of a tumor-specific LTR10A VNTR expansion on chr1. Matched tumor/normal reads from patient C568 are derived from a long-read nanopore sequencing dataset of 20 patients with colorectal adenocarcinomas (Xu et al. 2023). The yellow highlight shows the approximate location of the predicted tumor-specific mosaic expansion, as called by Sniffles2 (Smolka et al. 2023). Orange boxes indicate insertions and numbers within orange boxes indicate insertion sequence lengths (in bp).

### Figure S6M

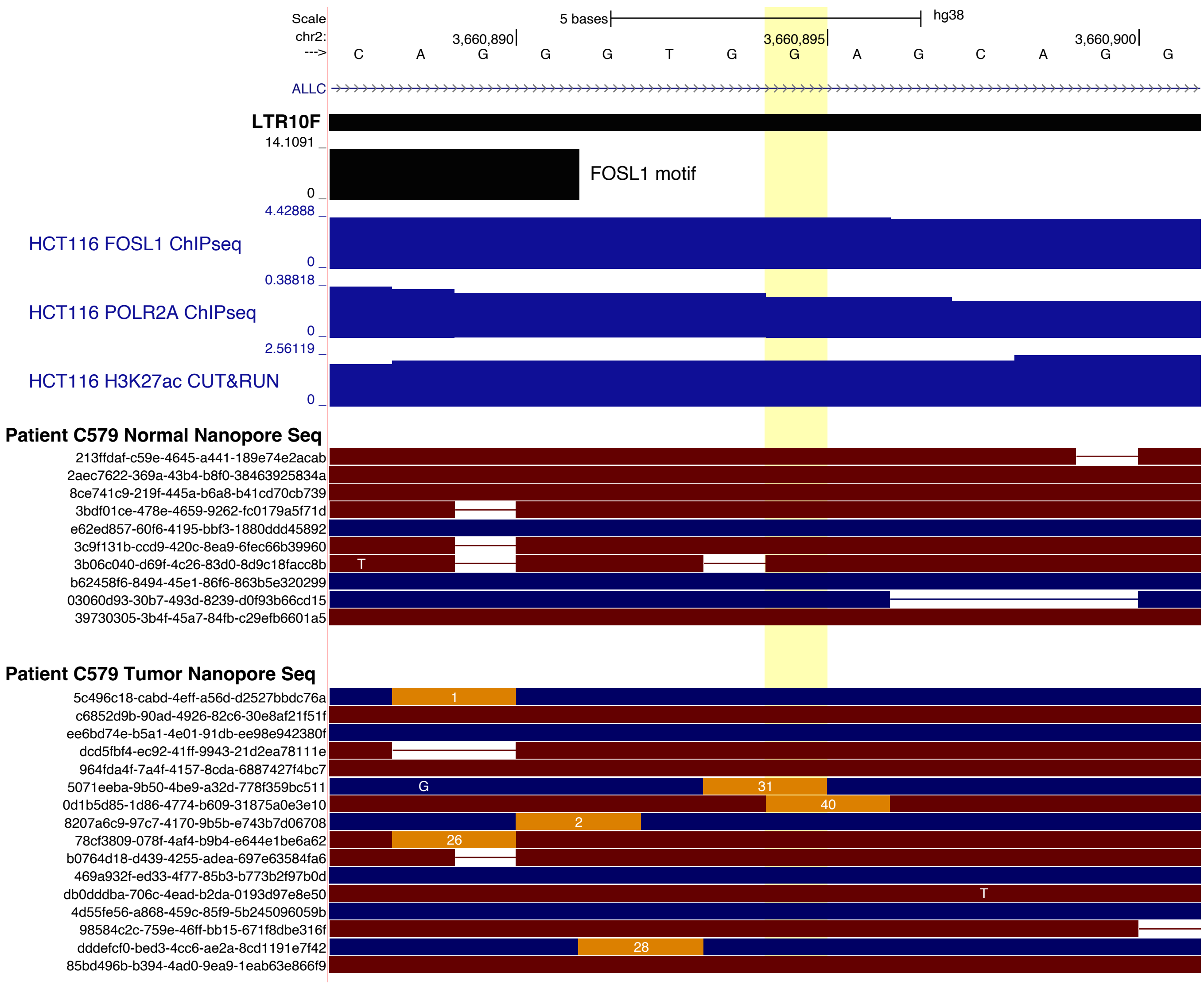


**Figure S6M:** Genome browser screenshot of a tumor-specific LTR10F VNTR expansion on chr2. Matched tumor/normal reads from patient C579 are derived from a long-read nanopore sequencing dataset of 20 patients with colorectal adenocarcinomas (Xu et al. 2023). The yellow highlight shows the approximate location of the predicted tumor-specific mosaic expansion, as called by Sniffles2 (Smolka et al. 2023). Orange boxes indicate insertions and numbers within orange boxes indicate insertion sequence lengths (in bp).

### Figure S6N

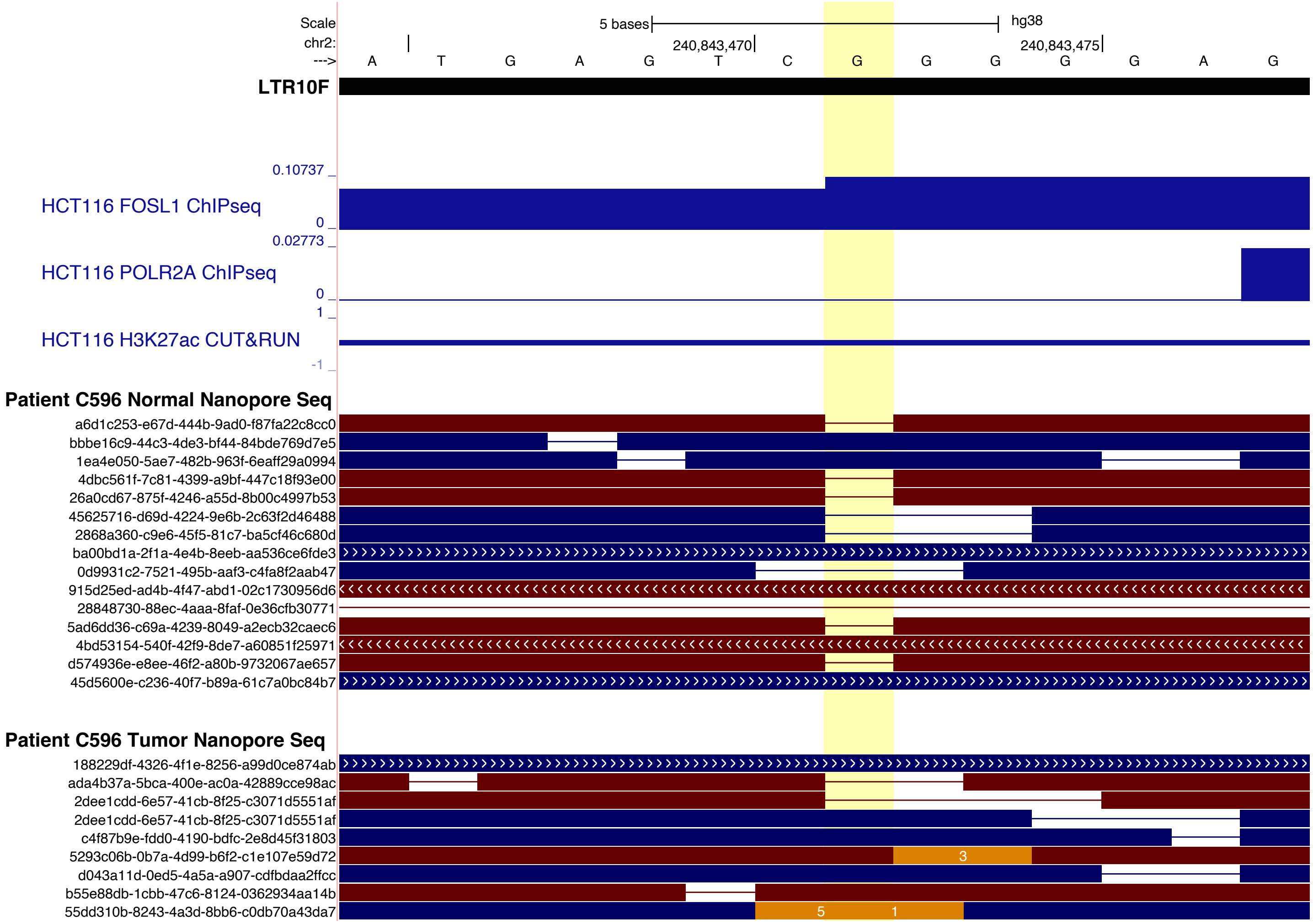


**Figure S6N:** Genome browser screenshot of a tumor-specific LTR10F VNTR expansion on chr2. Matched tumor/normal reads from patient C596 are derived from a long-read nanopore sequencing dataset of 20 patients with colorectal adenocarcinomas (Xu et al. 2023). The yellow highlight shows the approximate location of the predicted tumor-specific mosaic expansion, as called by Sniffles2 (Smolka et al. 2023). Orange boxes indicate insertions and numbers within orange boxes indicate insertion sequence lengths (in bp).

#### Figure S6O

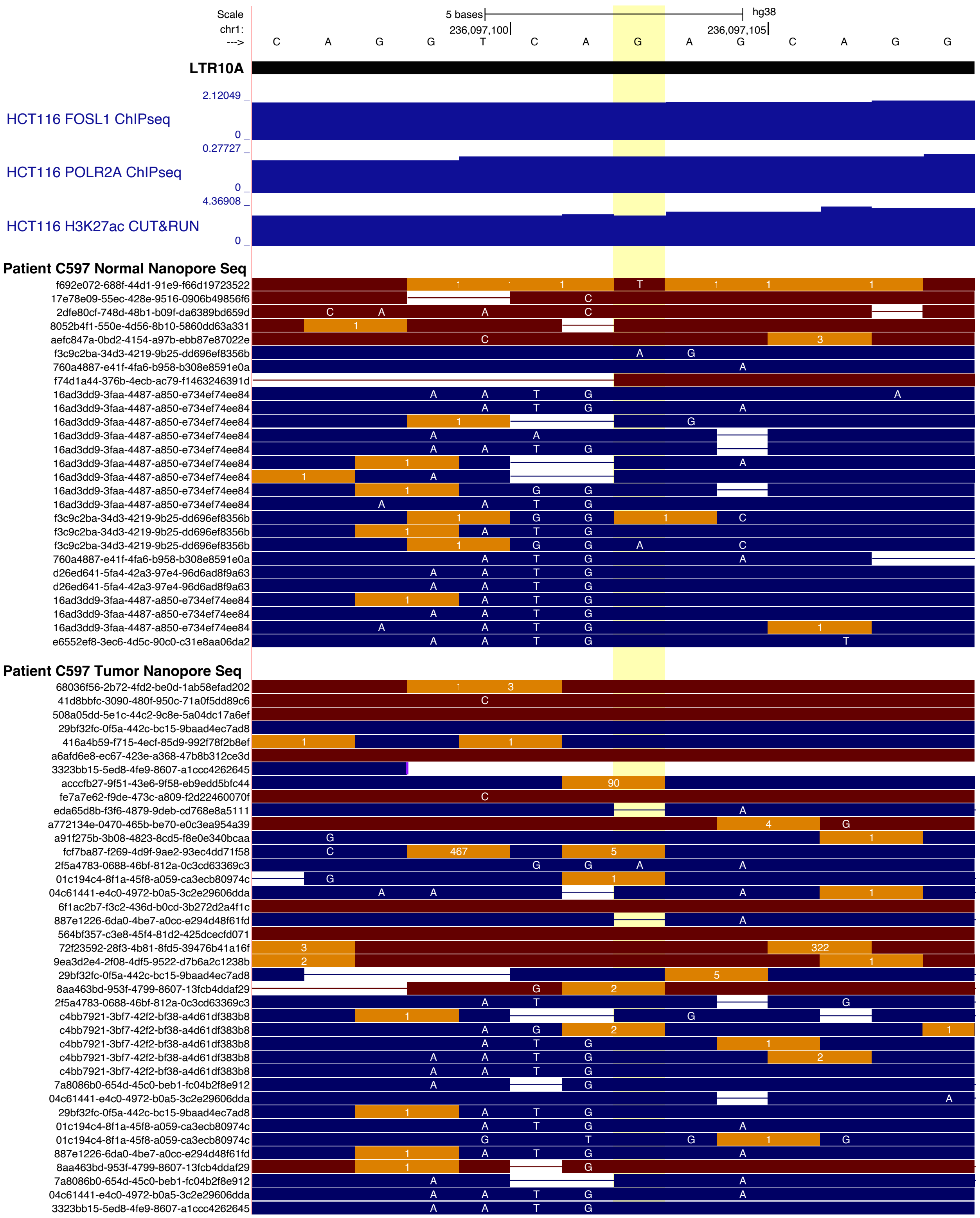


**Figure S6O:** Genome browser screenshot of a tumor-specific LTR10A VNTR expansion on chr1. Matched tumor/normal reads from patient C597 are derived from a long-read nanopore sequencing dataset of 20 patients with colorectal adenocarcinomas (Xu et al. 2023). The yellow highlight shows the approximate location of the predicted tumor-specific mosaic expansion, as called by Sniffles2 (Smolka et al. 2023). Orange boxes indicate insertions and numbers within orange boxes indicate insertion sequence lengths (in bp).

### Figure S6P

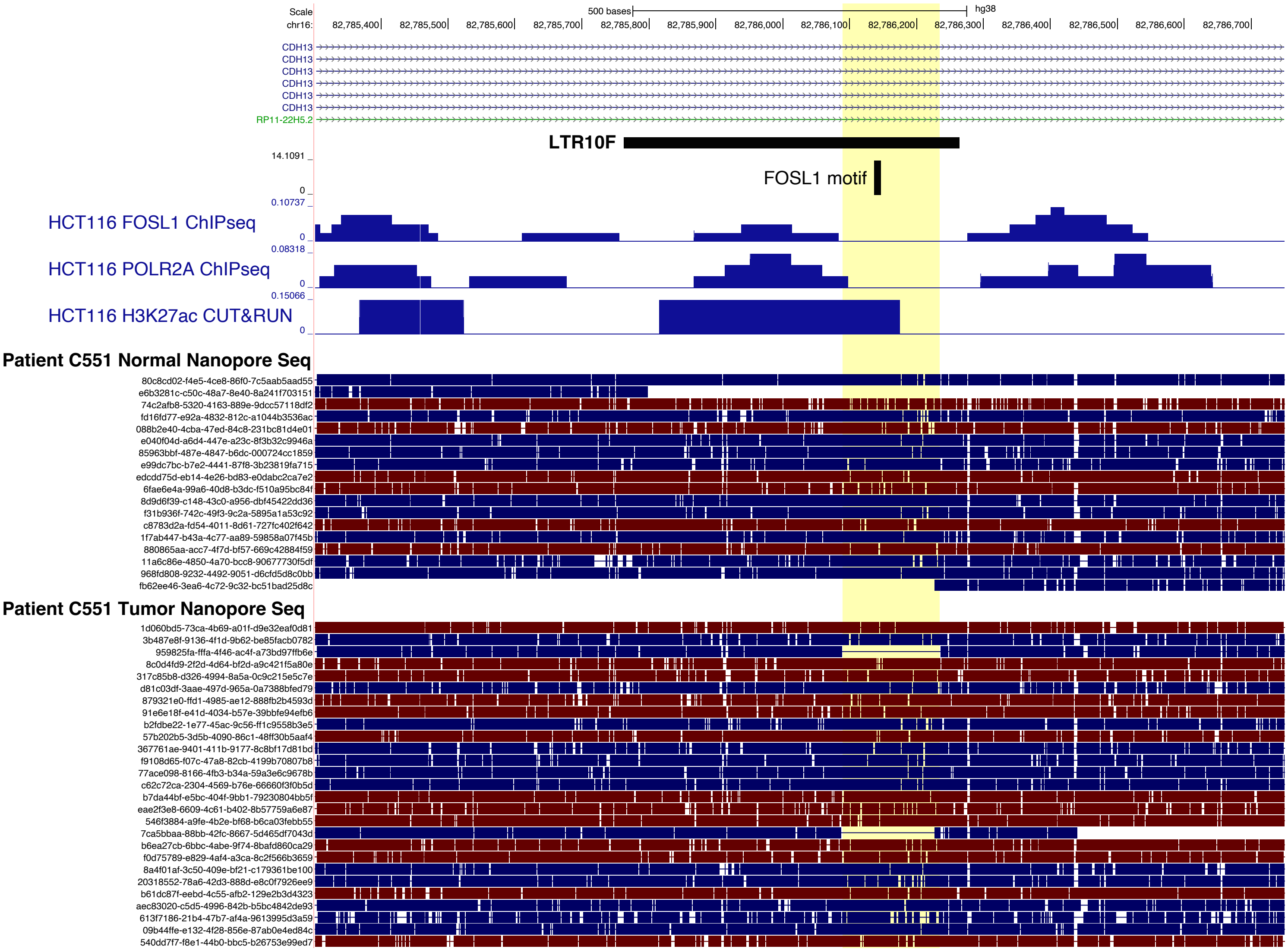


**Figure S6P:** Genome browser screenshot of a tumor-specific LTR10F deletion on chr16. Matched tumor/normal reads from patient C551 are derived from a long-read nanopore sequencing dataset of 20 patients with colorectal adenocarcinomas (Xu et al. 2023). The yellow highlight shows the location of the predicted tumor-specific mosaic deletion, as called by Sniffles2 (Smolka et al. 2023).

#### Figure S6Q

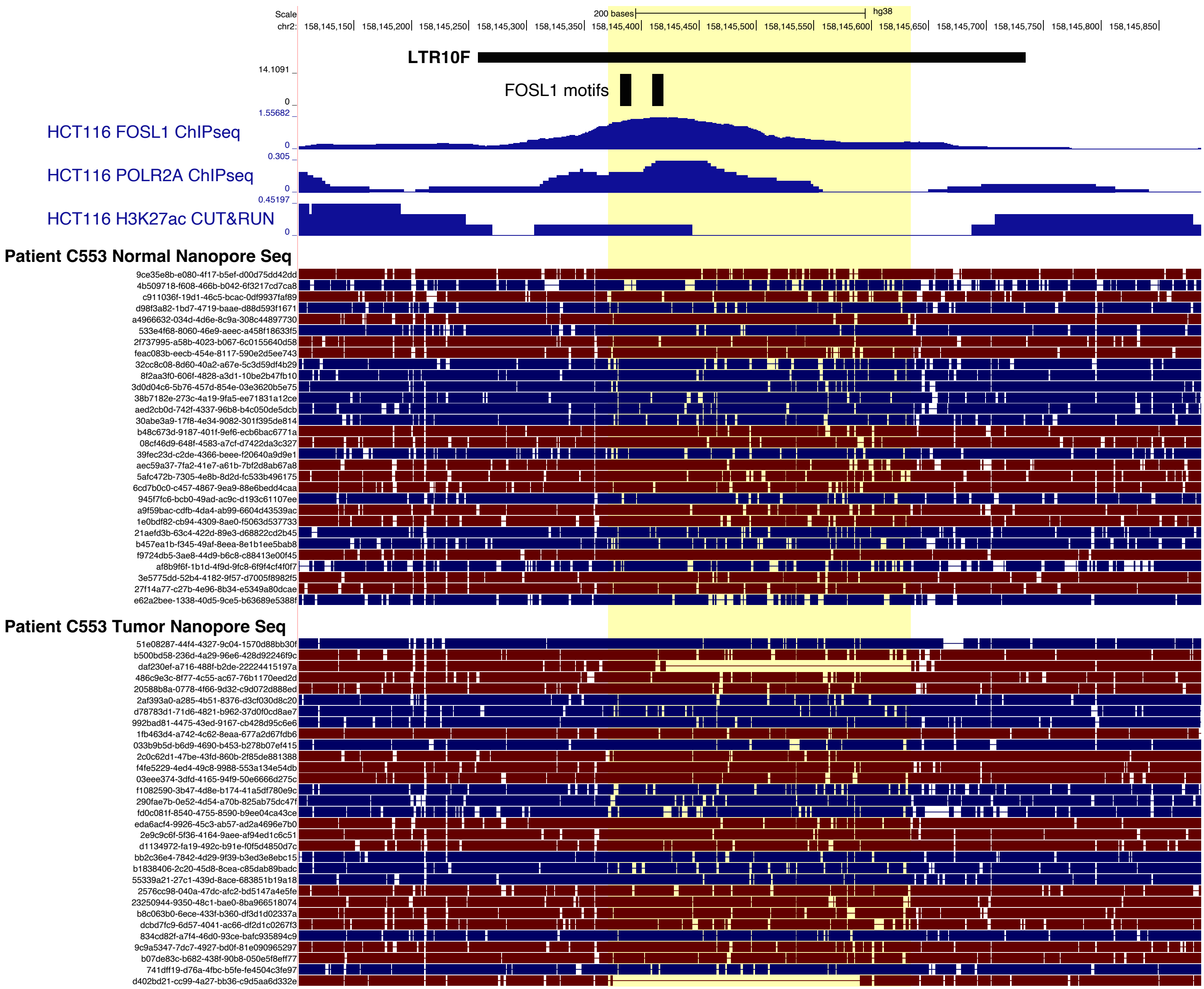


**Figure S6Q:** Genome browser screenshot of a tumor-specific LTR10F deletion on chr2. Matched tumor/normal reads from patient C553 are derived from a long-read nanopore sequencing dataset of 20 patients with colorectal adenocarcinomas (Xu et al. 2023). The yellow highlight shows the approximate location of the predicted tumor-specific mosaic deletion, as called by Sniffles2 (Smolka et al. 2023).

### Figure S6R

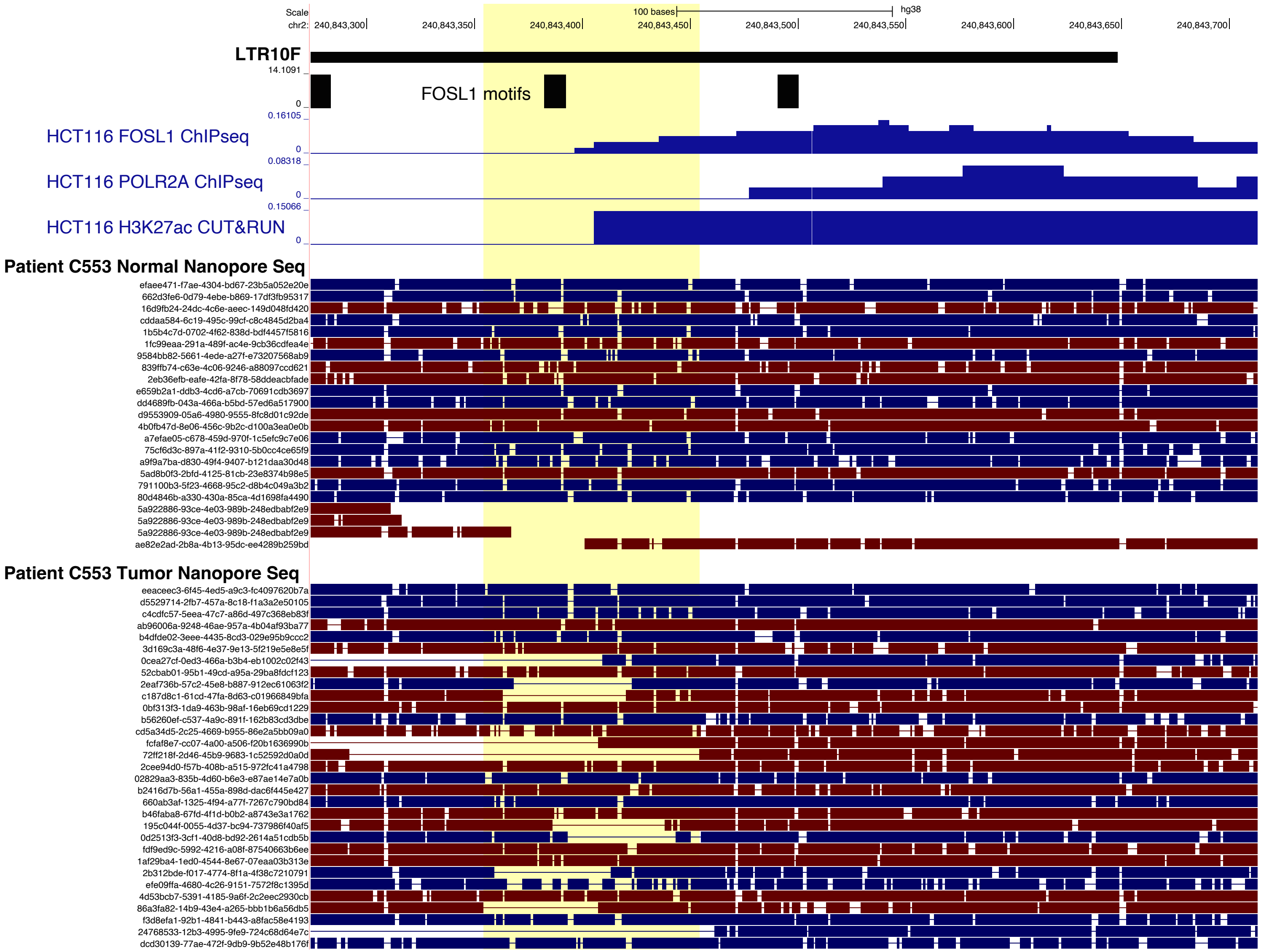


**Figure S6R:** Genome browser screenshot of a tumor-specific LTR10F deletion on chr2. Matched tumor/normal reads from patient C553 are derived from a long-read nanopore sequencing dataset of 20 patients with colorectal adenocarcinomas (Xu et al. 2023). The yellow highlight shows the approximate location of the predicted tumor-specific mosaic deletion, as called by Sniffles2 (Smolka et al. 2023).

### Figure S6S

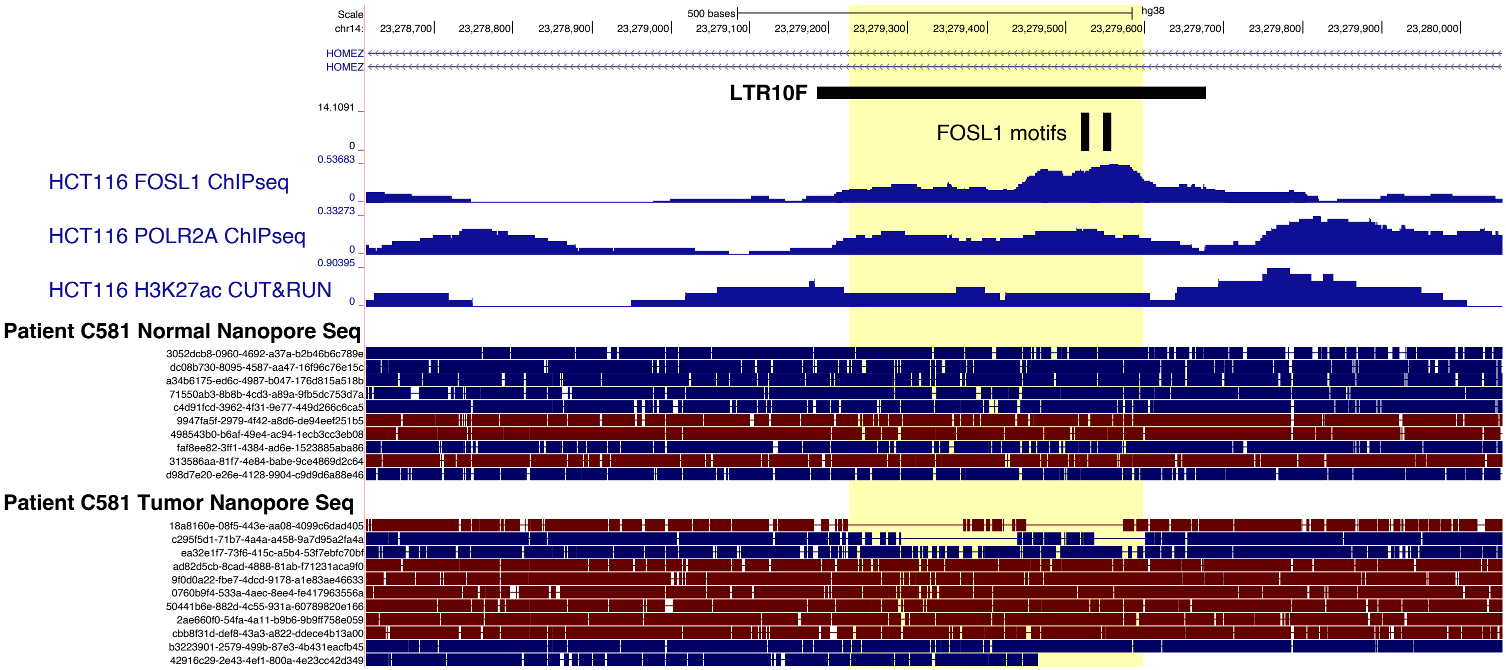


**Figure S6S:** Genome browser screenshot of a tumor-specific LTR10F deletion on chr14. Matched tumor/normal reads from patient C581 are derived from a long-read nanopore sequencing dataset of 20 patients with colorectal adenocarcinomas (Xu et al. 2023). The yellow highlight shows the approximate location of the predicted tumor-specific mosaic deletion, as called by Sniffles2 (Smolka et al. 2023).
